# Vanadium-dependent haloperoxidases from diverse microbes halogenate exogenous alkyl quinolone quorum sensing signals

**DOI:** 10.1101/2024.07.31.606109

**Authors:** Jackson T. Baumgartner, Catherine S. McCaughey, Hanna S. Fleming, Adam R. Lentz, Laura M. Sanchez, Shaun M. K. McKinnie

## Abstract

Site-selective vanadium-dependent haloperoxidases (VHPOs) are a unique enzyme family that catalyze selective halogenation reactions previously characterized within bacterial natural product biosynthetic pathways. However, the broader chemical roles and biological distribution of these halogenases remains to be explored. Using bioinformatic methods, we have defined a VHPO subfamily that regioselectively brominates alkyl quinolone (AQ) quorum sensing molecules. *In vitro* AQ halogenation activity was demonstrated from phylogenetically distinct bacteria lacking established AQ biosynthetic pathways and sourced from diverse environments. AQ-VHPOs show high sequence and biochemical similarities with negligible genomic synteny or biosynthetic gene cluster co-localization. Exposure of VHPO-containing microbes to synthetic AQs or established bacterial producers identifies the chemical and spatial response to subvert their bacteriostatic effects. The characterization of novel homologs from bacterial taxa without previously demonstrated vanadium enzymology suggests VHPO-mediated AQ bromination is a niche to manipulate the chemical ecology of microbial communities.

**Table of Contents Graphic:** 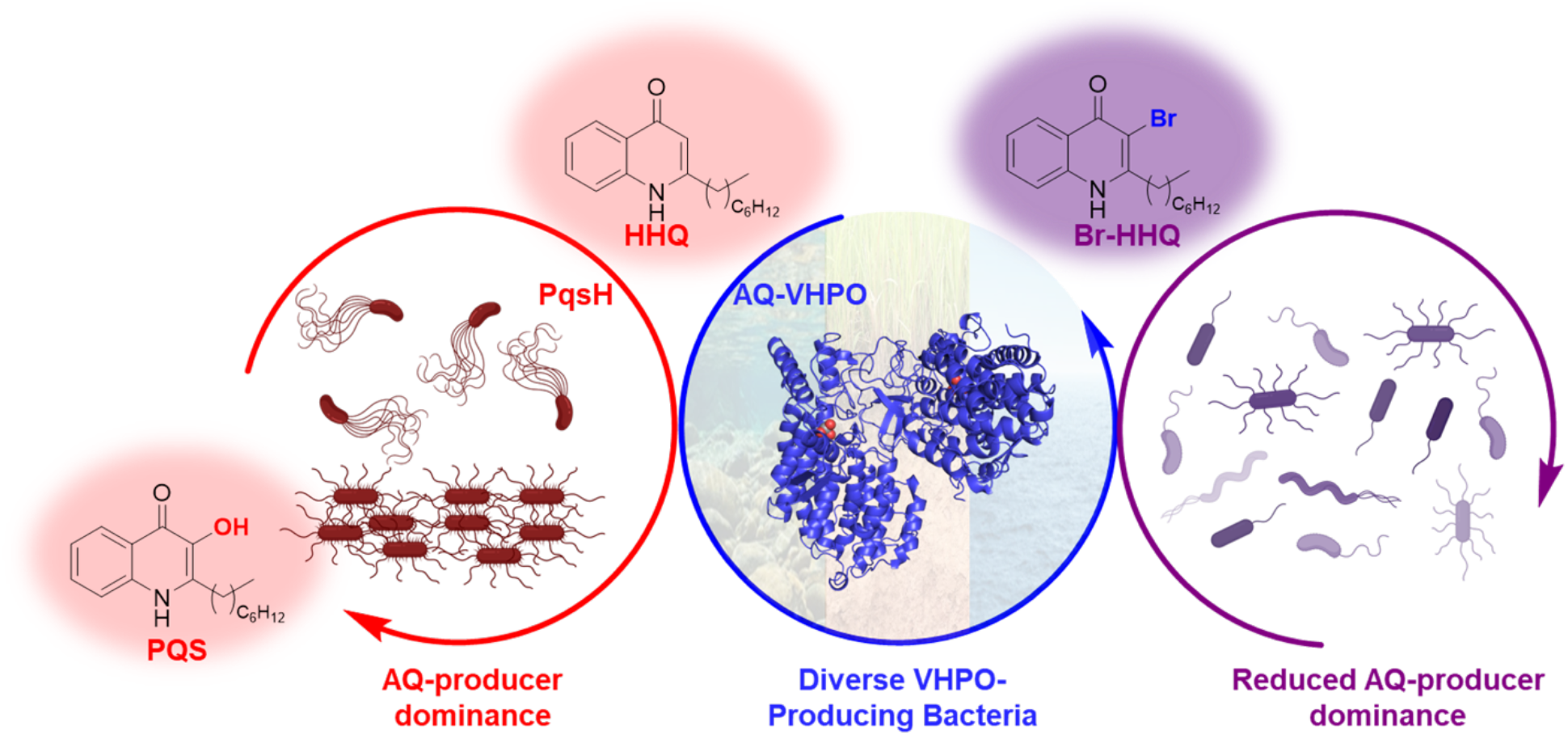

**Synopsis:** A broadly distributed subfamily of bacterial site-selective VHPOs provides protection against exogenously encountered bacteriostatic alkyl quinolone quorum sensing signals.

## Introduction

Vanadium-dependent haloperoxidases (VHPO) are a unique family of enzymes that use a histidine-bound vanadate cofactor and hydrogen peroxide to oxidize aqueous halide ions to hypohalous acid.^1–3^ Historically, this family of enzymes was believed to release diffusible hypohalous acid to spontaneously react with electron rich molecules.^1^ Macroalgal, cyanobacterial, and fungal VHPOs were rigorously characterized and are believed to play roles in antifouling mechanisms, dissolved organic matter degradation, haloform production, and halogenated natural product formation.^1^ For example, dpVHPO from *Delisea pulchra* provides antifouling activity via bromofuranone synthesis and acyl homoserine lactone (AHL) degradation.^4^ In contrast, characterized substrate-selective and stereospecific terpene cyclizing macroalgal and meroterpenoid biosynthetic streptomycete VHPOs display chemically complex reactions, making them intriguing candidates for biocatalytic investigation (Figure 1).^5–10^ Outside of natural product biosynthesis, the functional and ecological roles of site-selective bacterial VHPOs remains underexplored. Aside from the meroterpenoid biosynthetic VHPOs, only one site-selective bacterial VHPO has been characterized. HZ11-VHPO was isolated from the gammaproteobacterium *Microbulbifer* sp. HZ11 and found to regioselectively brominate the C3 position of alkyl quinolones (AQs), an important family of quorum sensing (QS) molecules.^11^ AQ C3-bromination conferred protection against the bacteriostatic effect of AQs to the VHPO-producing organism and modulated their toxicity against other environmental strains. Despite the lack of AQ production under laboratory conditions, *Microbulbifer* sp. HZ11 encodes an AQ biosynthetic gene cluster (BGC), suggesting the potential for the organism to endogenously produce QS molecules in Nature. However, AQs are also ubiquitous environmental virulence factors, implying a dual role for HZ11-VHPO in endogenous and exogenous substrate halogenation.^11^ The demonstrated activity against AQ toxicity suggests that HZ11-VHPO and its homologs represent a novel avenue to understand the ecological role of site-selective bacterial VHPOs.

**Figure 1.**
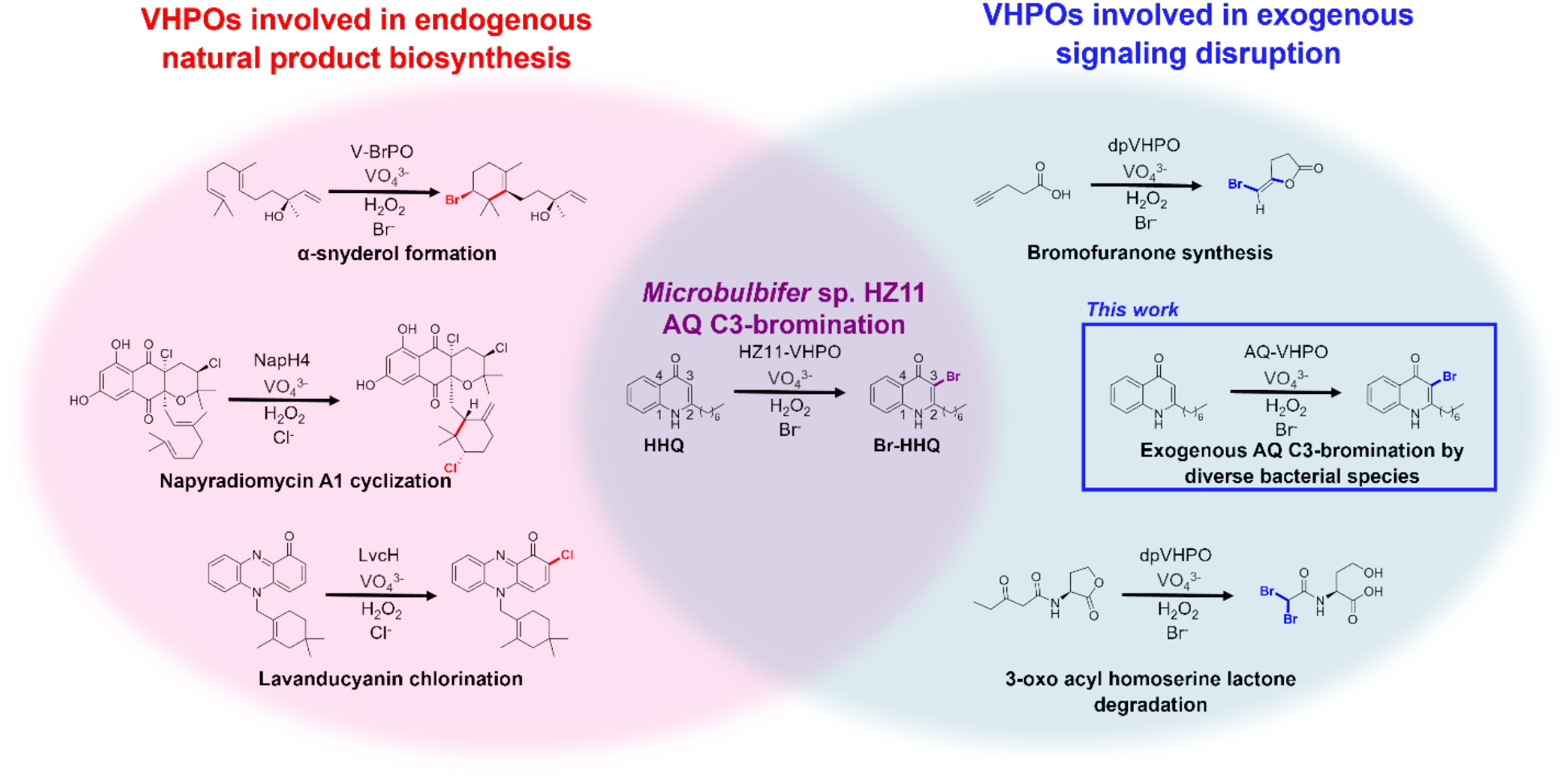
Comparison of the activity of VHPOs on small molecule substrates and the sources of the substrates. Endogenously produced bioactive natural product substrates in which the VHPO plays an established role in the biosynthetic pathway (red).^8–10^ Substrates encountered exogenous to the VHPO-producing organism and the respective chemical transformations catalyzed (blue).^4^ The reaction catalyzed by HZ11-VHPO has the potential to brominate both endogenously and exogenously produced AQs (purple).^11^ This work (blue box) expands the AQ-VHPO homologs as a broader subfamily that modify exogenously produced QS substrates.

Quorum sensing is a conserved bacterial communication system that coordinates activities including virulence factor production, biofilm formation, cell-to-cell interaction, and interkingdom interaction.^12–14^ In Gram-negative bacteria, small organic molecules (such as AHLs and AQs) are typically utilized as QS signals.^12^ The inverse is quorum quenching (QQ), wherein organisms utilize small molecule mimics or enzymatic modification to interfere with the QS pathways of other organisms.^15–17^ QQ provides alternative responses to wide ranging issues in antibiotic resistance, microbiomes, aquaculture, and wastewater treatment from traditional bactericidal means.^15,18–21^

Many AHL degradative enzymes and a variety of inhibitors of the corresponding LuxI/LuxR signaling system have been discovered and developed, however the degree of understanding on the QQs of other QS systems has lagged significantly.^16^ One such system is the AQ-based *pqs* signaling system, well known for its QS role in human pathogen *Pseudomonas aeruginosa* and other *Pseudomonas*/*Burkholderia* species. AQs also facilitate interkingdom communications and can elicit direct antibiotic or cytotoxic effects.^22–25^ The most prominent compounds of this pathway are 2-heptyl-4(1*H*)-quinolone (HHQ) and 2-heptyl-3-hydroxy-4(1*H*)-quinolone (commonly referred to as Pseudomonas quinolone signal, PQS), however many naturally occurring variations of these molecules have been isolated from diverse microbes.^25–27^ To the best of our knowledge, currently characterized AQ degradation/modification pathways begin with the C3-hydroxylated PQS, and result in oxidative ring cleavage, additional C2/C3 oxidation, and/or glucuronidation.^28–31^ To initiate the formation of PQS, which has substantial roles in *P. aeruginosa* QS, motility, and virulence, the flavin-dependent monooxygenase PqsH catalyzes the regioselective C3 hydroxylation of HHQ. Relatively conservative synthetic or enzymatic modifications to the C3 position in AQ derivatives largely reduce their antimicrobial and biofilm-inducing properties, thereby highlighting the significance of this position for bioactivity. This has been recently explored via CeO2 nanocrystal-catalyzed *in situ* hypobromous acid production, resulting in the non-enzymatic C3-bromination of 2-heptyl-4-hydroxyquinoline-*N*-oxide (HQNO) and reduction of *P. aeruginosa* biofilm growth.^32^ Furthermore, incubation of HHQ with *in situ* enzymatically (algal non-selective VHPO) or synthetically produced aqueous hypobromous acid have allowed for a proposed pathway of bromination triggered HHQ degradation.^33^ These cumulative findings, including HZ11-VHPO C3-bromination activity, the degradative effects triggered by C3-HHQ bromination, and reported activity of Br-HQNO suggest that AQ modification by selective bacterial VHPOs represent an underexplored QQ mechanism.

In this study, we show that AQ-modifying VHPOs are not exclusive to *Microbulbifer* sp. HZ11, but widespread throughout diverse bacterial taxa and environmental sources. These enzymes represent a conserved family utilizing rare vanadate biochemistry to provide defensive measures against environmentally encountered AQs. Additionally, we provide a basis for which this transformation can occur in the environment. Through these efforts, we have established: (1) VHPO-dependent AQ C3-bromination is present across multiple microbial species and environments; (2) organisms without established endogenous AQ genes are capable of brominating exogenous AQs to rescue bacterial growth; and (3) this modulation occurs *in situ* between pairwise microbial interactions of VHPO producers and an AQ-producing competitor bacterial species. Ultimately, this reveals VHPO-mediated AQ bromination as a widespread mechanism for modulating AQ signaling and toxicity in the polymicrobial milieu. Future ecological studies of this interaction will provide insight into the more general roles of VHPOs in interspecies interaction as well as into the variety of quorum quenching mechanisms utilized by bacterial species.

## Results

### AQ-VHPOS are distributed across diverse species and environments

We began our bioinformatic investigation into putative AQ-modifying VHPOs by constructing a manually curated Sequence Similarity Network (SSN) using the top 500 most similar sequences to HZ11-VHPO within NCBI and UniProt databases (Figure S1a). In the process of refining the network to contain distinct clusters or sub-clusters, several putative AQ-VHPOs that had clustered with the HZ11-VHPO sub-cluster under less strict alignment thresholds were excluded early in a secondary sub-cluster (Figures S1b and S1c). These VHPOs were included in subsequent analyses to ensure that putative AQ-modifying homologs were not eliminated through over-refinement. A total of 75 non-redundant putative AQ-modifying VHPOs from assembled and metagenomic sources were identified (Table S1). These genes are predominantly found in Gram-negative gammaproteobacteria; however, homologs from betaproteobacteria, deltaproteobacteria, acidobacteria, actinomycetes, and chloroflexi were also discovered. The source organisms are widely distributed across diverse marine, freshwater, and terrestrial environments (Figure S2). This represents an interesting departure from characterized site-selective meroterpenoid bacterial VHPOs isolated strictly from actinomycetes from predominantly marine environments. Moreover, this broad distribution encouraged an exciting opportunity to re-imagine the ecological role that these VHPOs may be playing. We next analyzed the lineages of these putative AQ-VHPOs by assembling a phylogenetic tree that formed distinct regions (Figure 2a). Regions 1 and 3 of the tree are distinguished by a high degree of source-organism homogeneity. Contrastingly, region 2 of the tree displays heterogeneous VHPO source organisms with prominent intermixing between Classes. Despite the formation of distinct leaves for gammaproteobacteria, *Vibrio*, and chloroflexota species, VHPOs from these taxonomic families also contribute to the heterogeneity of Region 2. Furthermore, the diversity within Region 2 suggests that additional gene acquisition methods or selection pressures likely occurred.

**Figure 2.**
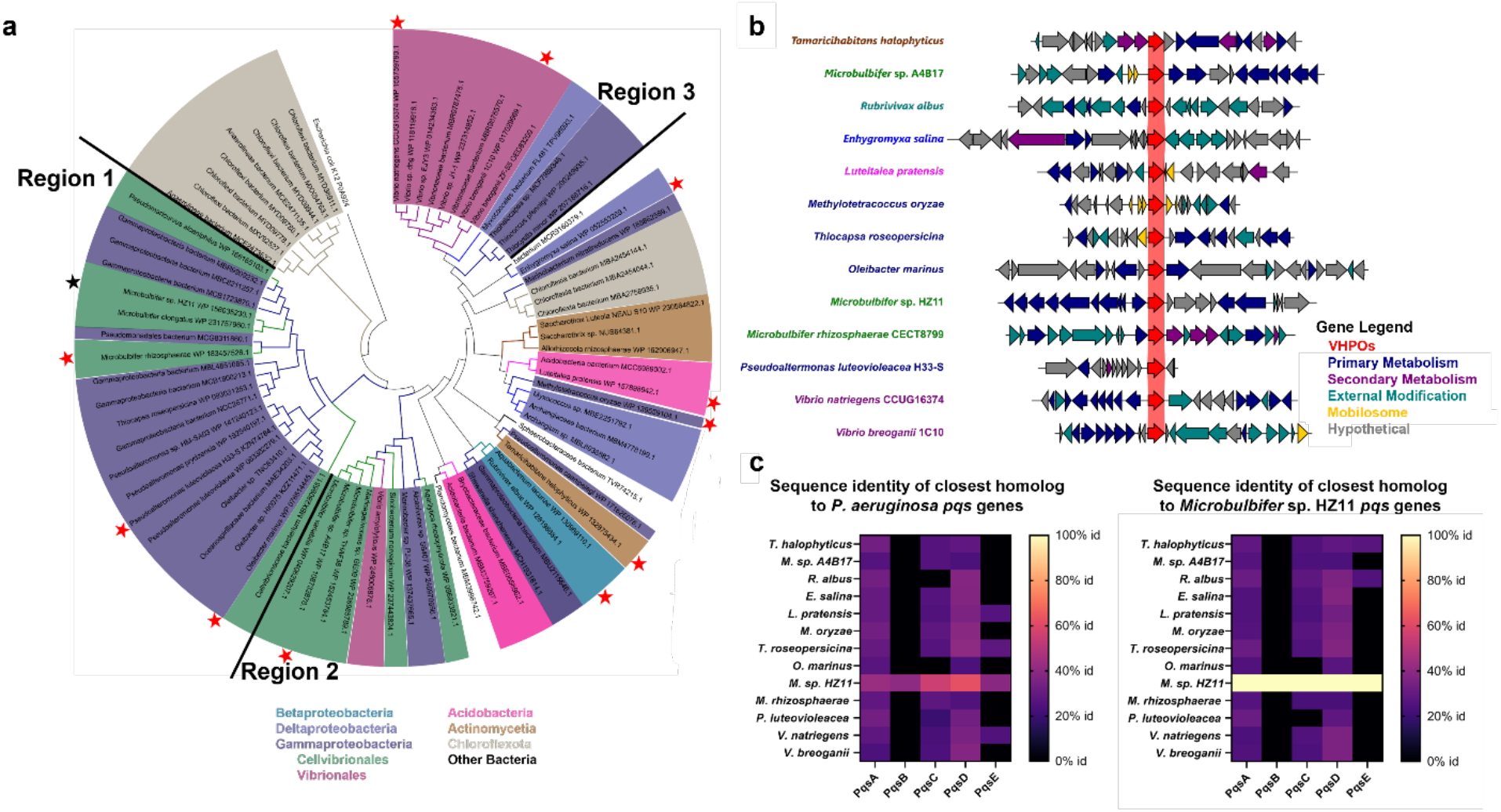
(a) Phylogenetic tree of putative AQ-VHPOs identified in this study. Tree is color coded by the Class of the source bacterium that the VHPO is derived from. Gammaproteobacteria is further split into Orders due to the large number of gammaproteobacterial source species. Previously characterized AQ-brominating HZ11-VHPO (black star)^11^ and the 13 novel AQ-VHPOs further explored in this study (red stars) are indicated. Distinct phylogenetic regions (1-3) are labeled. Bootstrap values have been removed for visual clarity. (b) The genomic contexts of 13 AQ-VHPOs (red arrow) selected to represent the diversity of microbial taxonomy and source environments identified in this study. Putative gene functions were defined based on COG definitions, classified into one of five general categories, and color coded accordingly. Shaded blocks connecting the arrows represent a percent sequence identity of 30%. (c) Heat map visualization of the general absence of alkyl quinolone biosynthetic pathways in species containing AQ-VHPOs, using either the *P. aeruginosa* (left) or *Microbulbifer* sp. HZ11 (right) *pqsABCDE* genes.

### AQ-VHPO homologs are found in variable and non-conserved genomic contexts

This inconsistent relation through common ancestry led us to investigate the genomic contexts of these VHPOs. Since most other selective bacterial VHPOs have been found in syntenic BGCs and characterized to produce secondary metabolites, we hypothesized that these VHPOs could be related through shared BGC contexts. We prioritized 13 VHPO genes from taxonomically divergent bacteria isolated from a range of source environments – however all VHPOs from assembled genomes were assessed bioinformatically (Table S2). Putative AQ-VHPOs were aligned and the adjacent genomic regions (10 genes upstream and downstream regardless of strand) were visualized using clinker^34^ (Figure 2b). Each gene was broadly attributed to essential cell functions, secondary metabolism, external response/modification, transposon/recombination-related elements, or hypothetical proteins based on BLAST annotations. Remarkably, putative AQ-modifying VHPOs were not generally present in secondary metabolism gene clusters and were in variable genomic environments, even within bacteria from similar genera. Additionally, 5 of the 13 aligned VHPOs were in regions with phage associated integrases or recombinases. In the case of *Methylotetracoccus oryzae*, 5 transposable elements immediately surrounded the AQ-VHPO. The presence of these elements suggests the involvement of horizontal gene transfer in the acquisition of these VHPOs in some species. Given the similar sequences of AQ-VHPOs to established meroterpenoid VHPOs (Figure S1), it was unexpected to observe negligible conserved biosynthetic gene co-localization. We have recently shown that biosynthetic VHPOs are not always present in BGCs via the discovery and characterization of LvcH, which catalyzes the regioselective chlorination of phenazine-derived meroterpenoid lavanducyanin.^9^ Moreover, the original AQ-VHPO in *Microbulbifer* sp. HZ11 was located 700 kbp away from *pqsABCDE* homologs.^11^ To investigate whether putative AQ-modifying VHPO bacterial strains have the biosynthetic capability to produce AQ molecules, we queried each assembled bacterial genome via antiSMASH 7.0^35^ for the complete *P. aeruginosa* PA14 *pqsABCDE* BGC (Figures 2c and S3) and manually BLAST searched for the presence of homologs required for AQ cyclization (*pqsBCE*) (Figure S3). No obvious *pqs* genes were observed using either of these bioinformatic queries except within *Microbulbifer sp*. HZ11 (Figure 2c, Table S2).^36^ Subsequent investigation using the *Microbulbifer* sp. HZ11 *pqs* genes failed to uncover more similar AQ biosynthetic machinery within putative AQ-VHPO containing microbes (Figure 2c). These bioinformatic results were counterintuitive to the current VHPO biosynthetic paradigm that implies co-localization within defined secondary metabolism BGCs and encouraged the investigation of AQ halogenation activity *in vitro*.

### VHPOs from diverse bacteria possess HHQ bromination activity

To interrogate the halogenation potential of these putative AQ-VHPOs, HZ11-VHPO and 13 novel genes from diverse bacterial taxa were codon optimized, assembled into pET28a(+) vectors, and expressed in *Escherichia coli* BL21(DE3) cells as N-terminal hexahistidine tagged constructs (Tables S3-S4). The putative AQ-VHPOs were prepared as crude lysates and screened for HHQ bromination activity following overnight incubation with aqueous buffered sodium orthovanadate, potassium bromide, and hydrogen peroxide. Lysate assays were analyzed by ultraperformance liquid chromatography mass spectrometry (UPLC-MS) and compared to synthetic 3-bromo-HHQ (Br-HHQ) by retention time and extracted ion chromatograms. Excitingly, Br-HHQ production was identified in 11 of the 14 putative VHPO homologs tested, including the previously established HZ11-VHPO^11^ (Figure 3a). Importantly, Br-HHQ was not observed in the absence of substrate, cofactors/co-substrates, or an empty pET28a(+) lysate control (Figures S4-S17). This lysate screen supports that VHPO-mediated HHQ bromination occurs across a wide range of bacterial taxa with varying levels of *in vitro* efficiency (Figure 3b). These AQ-VHPO genes are found in fresh water, soil, and marine microbes, showing a more widespread application of selective VHPO enzymology beyond the marine environment (Figure S18, Table S1). Given the diverse distributions of AQ-VHPOs across bacteria, we wanted to identify regions of residue conservation within the subfamily. Using the Alphafold 2.0 model of the most active homolog from the lysate screen, *Enhygromyxa salina* (esVHPO), we adapted the ConSurf server^37^ to predict the evolutionary variance of each spatially defined residue position across confirmed active AQ-VHPOs (Figure 3c). The internal core residues show high conservation across homologs while surface residues show more variability, suggesting the majority of AQ-VHPO functionality derives from internal residues. The previously proposed vanadate binding residues^1^ (K373, S427, R490, and H496) and the putative haloamine forming lysine^3^ (K329) are strictly conserved, reinforcing the selective VHPO activity of the homologs. The sequence and structural model alignments identified substantial variability in the N-terminal regions of AQ-VHPO homologs despite strong overall amino acid conservation (Figure 3c and Figure S19). SignalP 6.0^38^ analysis identified the presence of N-terminal secretion sequence signals in the majority of AQ-VHPO homologs, both within the subset we selected for experimental interrogation (Figure 3d) and all non-redundant sequences identified from bioinformatic databases (Figure S20). While signal peptides processed via the Sec/SPI pathway predominated, homologs with Sec/SPII and TAT/SPI pathway signals were also identified (Table S6). Furthermore, the presence of signal peptides appears primarily in bacterial VHPOs, while they do not appear in previously characterized eukaryotic or cyanobacterial homologs (Figure S21). These cumulative bioinformatic results imply that microbes containing AQ-VHPOs generally do not biosynthesize their AQ substrates directly, but instead secrete these enzymes to catalyze AQ halogenation reactions on exogenously encountered QS molecules. Moreover, the observed activities and residue conservation demonstrate that HHQ bromination activity is present across diverse species and environments.

**Figure 3.**
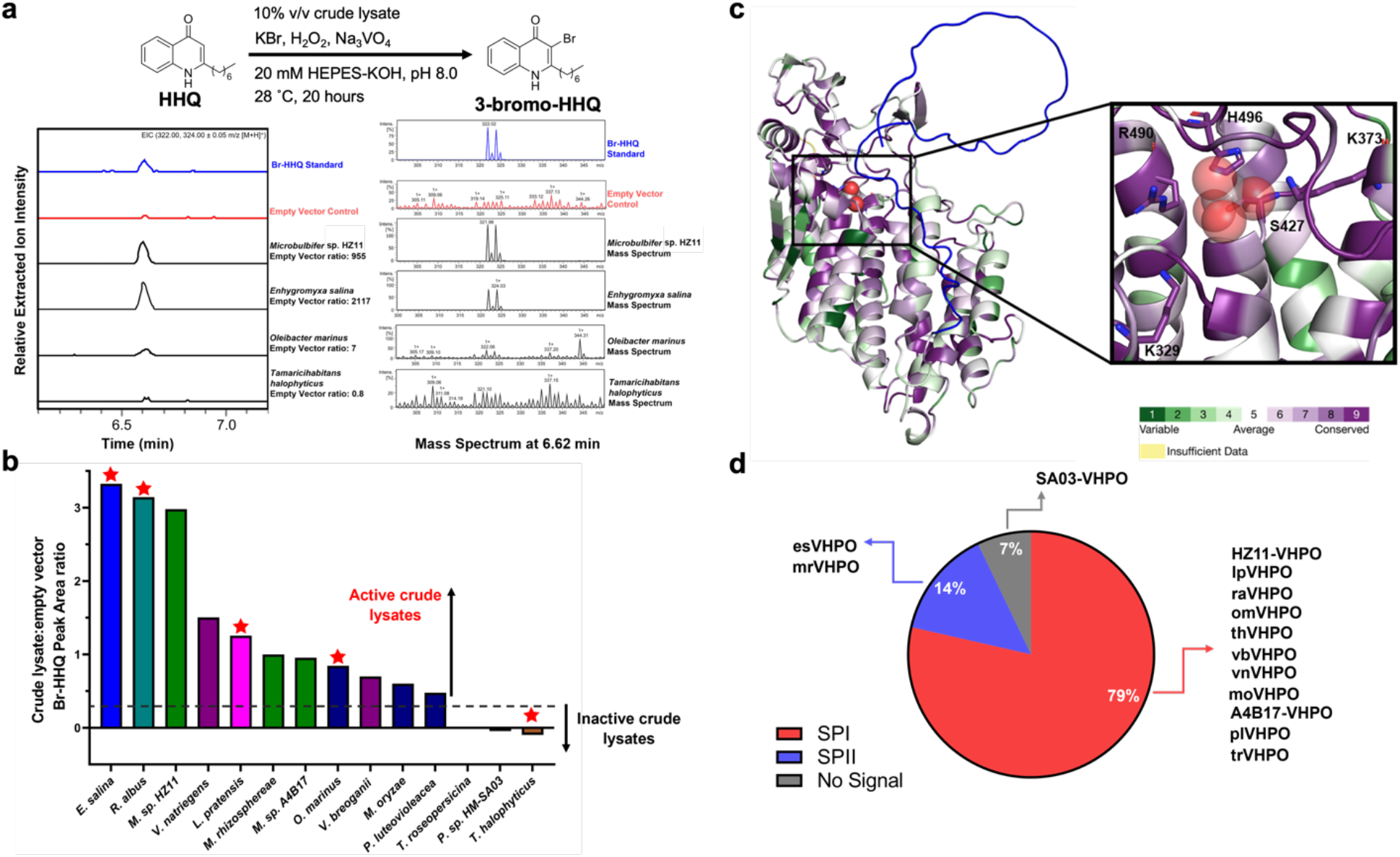
(a) Representative extracted ion chromatograms (EIC) traces of HHQ bromination activity by crude lysates of heterologously expressed AQ-VHPOs. Generalized reaction conditions, AQ substrate, and product are given above the traces. Synthetic Br-HHQ standard (blue), empty vector negative control (red), and experimental traces (black) are indicated. Empty Vector ratio represents the normalized size of the Br-HHQ peak area in the experimental condition compared to Empty Vector control. The MS for each extracted ion peak for each VHPO reaction is shown at the apex of its intensity. (b) HHQ bromination activity plot for all AQ-VHPOs heterologously expressed in crude lysates. Bars are color coded based on the phylogeny of the source organism defined in Figure 2 and red stars represent AQ-VHPOs that were further investigated through conventional protein purification. The y-axis plots the log scale ratio of the VHPO-containing crude lysate normalized Br-HHQ peak to the empty vector normalized Br-HHQ peak area. A ratio greater than 0.25 was arbitrarily considered active in this assay condition. (c) AlphaFold 2.0 model of esVHPO represented as a ribbon model. The vanadate ion (spheres) is overlaid from the crystal structure of the napyradiomycin A1 VHPO, NapH1 (PDBID: 3W36). Amino acid conservation grades were predicted by ConSurf software, where 1 (dark green) represents rapidly evolving positions, 5 (white) represents intermediate evolving positions, and 9 (dark purple) represents the most conserved positions. Insufficient data (yellow) results when fewer than 5 homologs have conserved residues in the region. The variable N-terminal region of esVHPO is highlighted (blue). The conserved vanadate pocket is zoomed in on and critical residues for selective VHPO activity are represented as sticks. (d) Distribution of predicted signal peptide categories based on SignalP 6.0 analysis for VHPOs with confirmed AQ-bromination activity.

### Selective C3-bromoperoxidase activity is conserved across AQ-VHPOs

To further interrogate AQ halogenation enzymology, four novel VHPOs that exhibited strong *in vitro* lysate activity were selected for protein purification, including homologs from: marine myxobacterium *Enhygromyxa salina* (esVHPO); soil acidobacteria *Luteitalea pratensis* H33-S (lpVHPO); freshwater betaproteobacterium *Rubrivivax albus* (raVHPO); and marine gammaproteobacterium *Oleibacter marinus* (omVHPO) (recently recommended to be renamed to *Thalassolituus maritimus*^39^). Despite previously reported solubility and purification challenges with *Microbulbifer* sp. HZ11-VHPO,^11^ we included it in our heterologous expression workflow as an established AQ-VHPO control. Lastly, the *Tamaricihabitans halophyticus* (thVHPO) homolog that did not demonstrate appreciable crude lysate activity was also selected to assess for any discrepancies in the initial lysate assay. Heterologous expression efforts following IPTG-induction and Ni-NTA affinity chromatography gave surprisingly variable results: esVHPO gave ∼5 mg/L purified product and small quantities (∼0.1 mg/L) of HZ11-VHPO were obtained; however, the other four constructs produced negligible soluble protein. Reassessment of the SignalP 6.0 results identified that esVHPO possesses a Sec/SPII signal peptide, while the other five homologs investigated were confidently Sec/SPI (Figure 3d). We hypothesized that this discrepancy could justify the variable heterologous expression results and inability to purify an N-terminally cleaved hexahistidine tagged protein that exhibited *in vitro* lysate activity. Recloning these four homologs with C-terminal hexahistidine tags and purifying from cell lysate (lpVHPO, raVHPO) or from culture media (omVHPO, thVHPO) gave improved quantities (0.1 – 2.5 mg/L) and purities to enable *in vitro* enzymology (Figure S22).

Our first step to characterize these novel VHPOs was to use the monochlorodimedone (MCD) assay to determine if the AQ-VHPOs generate diffusible hypobromous acid to react non-specifically with electron-rich substrates (Figure S23). Both HZ11-VHPO and esVHPO were unable to brominate MCD, suggesting that hypobromous acid is not being freely released. This contrasts with the robust MCD halogenation in the presence of bromide (but not chloride) ions by selective *Streptomyces* chloroperoxidase NapH1.^40^ This also deviates from previous QQ efforts performed using CeO2 nanocrystals and non-selective VHPOs that freely release oxidized halide. Upon its original characterization,^11^ HZ11-VHPO was shown to brominate a variety of AQ and heteroatom-containing aromatic substrates, with an *in vitro* preference for the longer alkyl chain 2-nonyl-4(1*H*)-quinolone (NHQ) and HHQ substrates. Our independent investigation with five naturally occurring AQ substrates (2-methyl-4(1*H*)-quinolone (MHQ), NHQ, HHQ, 2,4-dihydroxyquinolone (DHQ), and HQNO) mirrored the literature reports for HZ11-VHPO. However, different trends were observed for the four new AQ-VHPOs (Figure 4a). All five substrates were monobrominated by the novel VHPOs but exhibited an *in vitro* preference towards the shorter MHQ. Additional hydroxylation on either the quinolone ring (DHQ) or nitrogen (HQNO) generally reduced halogenation efficiency *in vitro* by these AQ-VHPOs. Subsequent characterization of enzymatic activities were carried out on MHQ due to its higher turnover. Incubation of each purified VHPO with all necessary cofactors and substrates resulted in the efficient halogenation of MHQ to 3-bromo-MHQ (Br-MHQ) following UPLC-MS analyses and comparison to synthetic standards (Figures 4b, S24). Subsequent co-substrate dependence of all novel purified VHPOs identified the strict requirement of bromide and hydrogen peroxide for Br-MHQ formation. In the absence of vanadate supplementation, esVHPO, lpVHPO, and thVHPO displayed negligible MHQ halogenation activity, however raVHPO and omVHPO showed robust production of Br-MHQ. Tryptic digestion and high resolution ICP-MS analyses of purified raVHPO measured ^51^V in medium resolution and identified quantifiable amounts of vanadate (∼1.3 ppb), whereas identical analyses with esVHPO were under the limit of detection. Previously characterized actinobacterial VHPOs require exogenous vanadate to reconstitute *in vitro* halogenation activity.^40^ This functional divergence suggests that certain AQ-VHPOs possess higher vanadate binding affinities capable of sequestering this trace mineral from heterologous growth conditions or require this cofactor for soluble folding. Next, we wanted to confirm the C3 regioselectivity of the novel VHPOs and the lack of ability to multiply brominate a single substrate. Synthetic standards of Br-MHQ and 3,6-dibromo-MHQ were prepared to benchmark the multiplicity of VHPO bromination (Figure 4c). esVHPO was selected as our model system and treated with excess potassium bromide and hydrogen peroxide concentrations. Even under superstoichiometric hydrogen peroxide concentrations, monobromination activity was retained without obvious MHQ overbromination or degradation (Figure 4c). A similar result was obtained for lpVHPO against an even more dramatic hydrogen peroxide gradient (Figure S25). In contrast, incubation of MHQ with increasing molar equivalents of chemical brominating reagent *N*-bromosuccinimide (NBS) showed production of the di- and tri-brominated products (Figure 4c). This highlights the high regioselectivity of esVHPO, and presumably the entire AQ-VHPO family. Finally, to assess the halide selectivity of novel AQ-modifying VHPOs, each novel purified homolog (esVHPO, lpVHPO, omVHPO, raVHPO) was assayed with MHQ while individually varying the halide source. Due to low soluble yields, thVHPO was omitted from this experiment. Following UPLC-MS analyses and comparison to synthetic 3-halogenated MHQ standards, all AQ-VHPOs showed both bromide and iodide oxidation and installation but no detectible chlorination, confirming their activities as bromoperoxidases (Figures 4d-f, S26-28). To further assess AQ-VHPO enzymes as a broader bacterial QQ mechanism, we next wanted to assess their activity within their microbial hosts.

**Figure 4.**
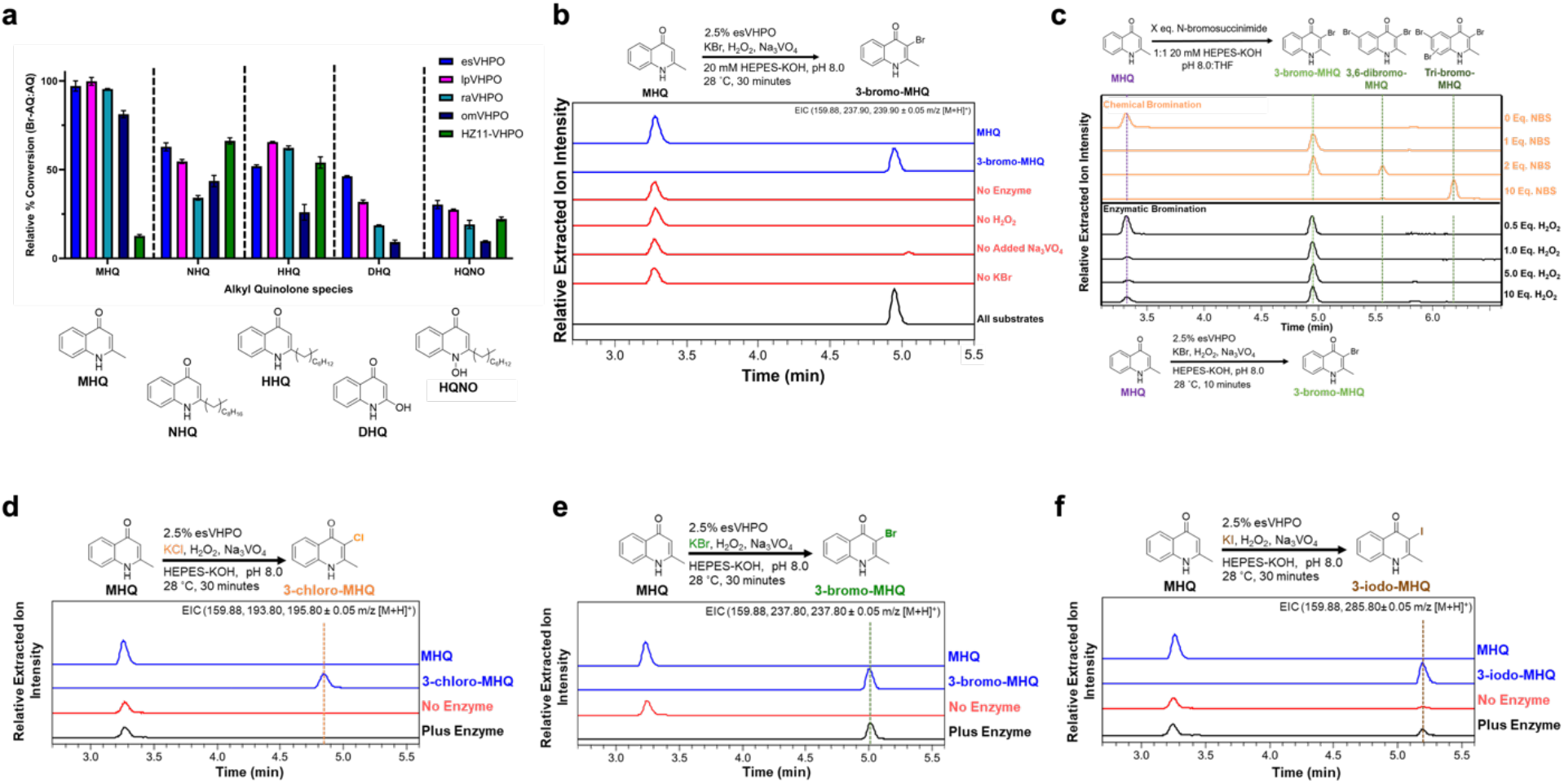
(a) Relative percent conversions of AQ to Br-AQ by each VHPO. Each bar represents the ratio of the normalized ion peak areas of Br-AQ to that of the unmodified AQ. Error bars represent the mean ± s.d. (n = 3 for all substrates and VHPOs). (b) Co-substrate dependencies of esVHPO. Synthetic standards (blue), negative control traces omitting the listed substrate (red), and the full enzyme reaction (black) traces are indicated. Extracted ions are given in the upper right corner of the figure. (c) EIC traces of MHQ chemical reaction with NBS at different stoichiometric equivalents (orange). EIC trace of MHQ enzymatic reaction with esVHPO in the presence of increasing stoichiometric equivalents of H_2_O_2_ (black). The mass ions for all 0 – 3 Br atom additions were extracted for each reaction condition trace. Halide oxidation profile of esVHPO with KCl (d), KBr (e), and KI (f). Generalized reaction conditions, AQ substrate, and products are given above EIC traces. Extracted ions are given in the upper right corner of each trace.

### omVHPO bromination of exogenous HHQ rescues growth in *Oleibacter marinus*

To confirm the sensitivity of AQ-VHPO containing strains to HHQ, we acquired three bacteria (*Oleibacter marinus* DSM 24913),^41^ *Microbulbifer rhizosphaerae* (DSM 28920),^42^ and *Vibrio natriegens* CCUG16374 (ATCC 33898)^43^ from strain repositories. Genomic DNA was extracted from each strain and the 16S rRNA and native VHPO genes were sequenced to confirm identity. Each genome was BLAST queried for alternative halogenases from major halogenase families (Table S8). For *O. marinus* and *V. natriegens*, the AQ-VHPO is the only predicted halogenase or haloperoxidase gene in their genomes; *M. rhizosphaerae* contains 8 putative FAD-dependent halogenases in addition to its one AQ-VHPO (Table S9). Because HHQ acts as a bacteriostatic agent,^22^ each organism was selected to represent a range of different bacterial growth speeds; with *V. natriegens, M. rhizosphaerae*, and *O. marinus* reaching stationary phases in 3 h, 15 h, and 48 h respectively (Figures 5a, S29, S30). We next tested the effects of HHQ and Br-HHQ on the growth rate of each organism. The addition of exogenous HHQ impacted each organism in a bacteriostatic manner that correlated to their unperturbed growth rates. *V. natriegens* displayed minimal sensitivity to HHQ, requiring 100 μM in culture conditions to observe any inhibitory growth effects (Figure S29). Under the same concentrations, the growth of *M. rhizosphaerae* was substantially slowed, indicating enhanced susceptibility to HHQ toxicity (Figure S30). *O. marinus* displayed strong dose-dependent sensitivity to HHQ, where comparatively low concentrations of HHQ dramatically arrested growth rates (Figure 5a). In contrast, incubation of each bacterium with equivalent concentrations of Br-HHQ showed no significant difference from the vehicle control, supporting previous reports of C3-bromination as a protective QQ modification (Figures 5b, S29-30).^8^ Repeating the same experiment with biofilm dispersing agent MHQ^44^ and its brominated variant, exhibited a modest dose-dependent decrease in *O. marinus* growth compared to the DMSO control (Figure S31). However, negligible differences were observed between the two compounds, showing that the halogenation of bacteriostatic AQs is likely more physiologically significant.

**Figure 5.**
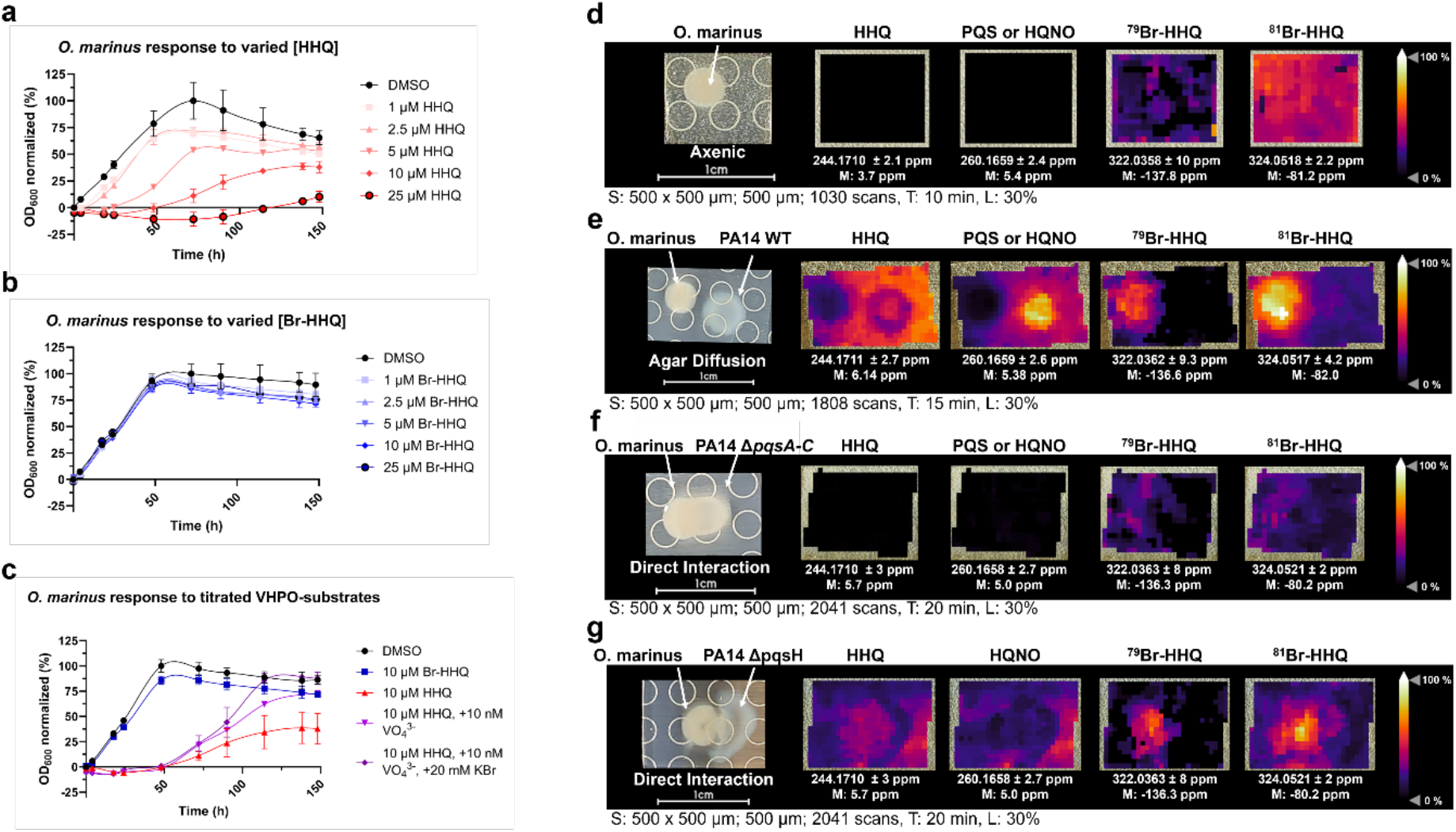
(a) Response of *O. marinus* to the increasing concentrations of synthetic HHQ. Error bars represent the mean ± s.d. (n=3). Lines do not represent a line of best fit and are added for visual clarity. Each [HHQ] growth curve is normalized to the DMSO growth curve. (b) Response of *O. marinus* to increasing concentrations of synthetic Br-HHQ. Data was analyzed and normalized as in Figure 5a. (c) Response of *O. marinus* to 10 μM HHQ in the presence of vanadate and bromide supplementation. Data was analyzed as in Figure 5a. Each media condition growth curve was normalized to the DMSO growth curve from the same media condition. (d) The MALDI-QqTOF MSI visualization of axenic *O. marinus*. Mass ions for HHQ, PQS/HQNO, ^79^Br-HHQ and ^81^Br-HHQ were identified and visualized. (e) The MALDI-QqTOF MSI visualization of *O. marinus* co-cultured with wild type *P. aeruginosa* PA14. (f) The MALDI-QqTOF MSI visualization of *O. marinus* co-cultured with the AQ knockout strain *P. aeruginosa* PA14 *ΔpqsA-C*. (g) The MALDI-QqTOF MSI visualization of *O. marinus* co-cultured with the PQS-deficient strain *P. aeruginosa* PA14 *ΔpqsH*. For MSI, signal intensity is displayed as a heatmap and shows the spatial distribution of the visualized mass ion. Spot raster; size; scan number (S), mass interval confidence (M), acquisition time (T), and laser power (L) are given in the figure. The annotation (A) for each experiment was targeted. The resolving power (R) of the instrument used for all MALDI-MSI data collection was 60000_FWMH_ at 1200 *m/z*.

To assess *in vivo* AQ halogenation, each strain was propagated in the presence of HHQ, sodium orthovanadate, and potassium bromide. Following ethyl acetate extraction and UPLC-MS analysis, Br-HHQ was successfully observed in the culture of *O. marinus* by retention time and extracted ion chromatogram comparison to a synthetic standard (Figure S32). This metabolite was not observed in extracts of the other two bacteria, possibly due to alternative AQ degradation or detoxification mechanisms present within faster growing microbes. The successful formation of Br-HHQ by *O. marinus* cultures encouraged us to further investigate this microbe for *in vivo* omVHPO activity. Quantitative polymerase chain reaction (qPCR) confirmed constitutive *omVHPO* transcription in axenic *O. marinus* culture, with a general decrease in transcripts in the vehicle control as the culture approached stationary phase (Figure S33). In contrast, *omVHPO* mRNA transcripts were either slightly increased or sustained in the presence of tolerated HHQ concentrations (1 or 10 μM) once normalized to the first collected time point. To influence omVHPO activity in culture, *O. marinus* was again incubated in HHQ gradients, however the culture media was now supplemented with AQ-VHPO co-factors and co-substrates (10 nM sodium orthovanadate and/or 20 mM potassium bromide). While each additive was individually effective at restoring microbial growth, the combination of both was the most impactful for *O. marinus*; this trend became more pronounced under higher AQ concentrations (Figures 5c, S34). In the absence of genetic manipulation methods, these microbiological interactions with VHPO substrates and HHQ suggests that the activity of omVHPO helps rescue the growth of *O. marinus* under bacteriostatic AQ pressure.

### AQs and Br-AQs localize differentially in agar co-culture between VHPO- and AQ-producing bacteria

To further explore the impact of *in vivo* AQ-VHPO enzymology, we turned to agar based microbial co-cultures between AQ-VHPO and AQ-producing bacteria, adapting matrix-assisted laser desorption/ionization – mass spectrometry imaging (MALDI-MSI) technology to visualize the spatial localization of metabolites. *O. marinus* was cultured on modified marine broth agar for 5 days before introduction of *P. aeruginosa* PA14. Experiments were conducted with either axenic *O. marinus*, or three co-culture experiments with *P. aeruginosa* PA14 strains, including: wild-type PA14 (WT); an AQ knockout strain (*ΔpqsA-C*); and a PQS deficient strain (*ΔpqsH*) previously used to study *pqs* control of phenazine production^41^ and recently adapted to study the biofilm inhibition of taurolithocholic acid.^42^ Microbes were co-cultured either in contact or separated by a short distance to allow chemicals to diffuse through the agar and were visually imaged (Figure S35) prior to MALDI-MSI analysis (Figure 5d-g). Co-culturing with both *P. aeruginosa* PA14 WT and *ΔpqsH* demonstrated a reduced *O. marinus* fitness consistent with the bacteriostatic effects of AQs. In contrast, incubation with AQ knockout strain *ΔpqsA-C* appeared visually consistent to the axenic culture (Figure S35). To identify the spatial location of metabolites, co-cultures were analyzed by MALDI-MSI and ion images corresponding to HHQ, PQS/HQNO, and both bromine isotopologs of Br-HHQ were analyzed (*m/z* [M+H]^+^: 244.1710, 260.1658, 322.0358, and 324.0518 respectively). Rationale for how particular mass ion peaks were selected for generating mass ion images, including low intensity peak-picking, are included in the Supporting Information for all MALDI-MSI images (Figures S37-S40, S43, S44, S46). Axenic *O. marinus* failed to give appreciable signal intensities for endogenous AQ production, consistent with the bioinformatic results that this microbe lacks the established biosynthetic machinery (Figure 5d). While a background ion was observed for ^81^Br-HHQ, it did not reproduce an equal intensity ^79^Br-HHQ isotopolog and is most likely attributed to a media component based on liquid culture analyses (Figure S32). This difference in bromine isotopolog intensities by MALDI-MSI was further supported by adding synthetic Br-HHQ directly onto the *O. marinus* colony prior to imaging (Figure S36a). Analysis of the *O. marinus* and *P. aeruginosa* PA14 WT co-culture identified the diffusion of HHQ from *P. aeruginosa* PA14 WT into the agar, while PQS/HQNO are observed localizing to the producer colony (Figure 5e and Figure S36b). However, both isotopologs of Br-HHQ are located within the *O. marinus* colonies, indicating that VHPO halogenation chemistry is occurring by the microbe. The absence of all AQ-derived signals in the *P. aeruginosa* PA14 *ΔpqsA-C* co-culture, including both Br-HHQ isotopologs, further supported that the brominated masses observed were due to modification of exogenous AQ from the Pseudomonad instead of the induction of a novel metabolite in *O. marinus* (Figure 5f). To serve as a control for the genetic methods, incubation with the *P. aeruginosa* PA14 *ΔpqsH* mutant capable of making AQs but unable to C3 hydroxylate them, restored all signals including Br-HHQ (Figure 5g). While colony-associated PQS appears to be ablated in this mutant, a remnant mass is still observed with greatest intensity at its farthest point from *O. marinus* and is presumably from *N*-hydroxylated regioisomer HQNO. Finally, Br-HHQ is again observed to be localized within *O. marinus*, further supporting microbial halogenation.

Despite the absence of Br-HHQ accumulation in liquid culture, we assessed solid agar co-cultures of *V. natriegens* CCUG 16374 or *M. rhizosphaerae* with *P. aeruginosa* PA14 by MALDI-MSI to determine if AQ halogenation is occurring in solid agar. Due to the faster growth rates of these microbes, all co-cultures were inoculated at the same time. Both *V. natriegens* and *M. rhizosphaerae* exhibited similar patterns of reduced fitness in the optical images when inoculated near *P. aeruginosa* PA14 strains capable of making AQs, with the *ΔpqsA-C* mutant mirroring axenic cultures (Figures S41, S45). Following MALDI-MSI analyses, both microbes generated Br-HHQ in solid culture and exhibited similar spatial organization of native and brominated AQs as the *O. marinus* co-cultures did (Figures S42, S46). A stronger media background was present in all experiments for the ^81^Br-HHQ ion, making the ^79^Br-HHQ isotopolog more diagnostic for the transformed AQ metabolite. Halogenated HHQ ions localized to the microbial interface between *V. natriegens*/PA14 and diffused throughout the colony region of *M. rhizosphaerae*; in all cases ions were notably absent in the *P. aeruginosa* PA14 colonies. Given the *in vivo* halogenation activity, we questioned what effect brominated AQs would have on *P. aeruginosa* PA14. We were unable to observe toxicity or growth rate reduction in PA14 cultures incubated with either HHQ or Br-HHQ (Figure S48) and saw a negligible impact on swarming motility (Figure S49). While the effect of Br-HHQ on swarming motility was subtle, it did appear to promote enhanced swarming areas, perhaps suggesting Br-HHQ interfering with the transition of *P. aeruginosa* PA14 from its planktonic to sessile state (Figure S49). These cumulative microbiological experiments suggest that VHPO-mediated AQ bromination occurs between bacteria when they meet in the environment, and has a more beneficial effect on the VHPO-producing bacteria.

## Discussion

Through our interdisciplinary approach to better understand AQ-VHPOs, we propose a model wherein these enzymes intercept and selectively brominate exogenously produced AQs to alter their biological effects. This work shows that AQ-VHPOs are present within the genomes of diverse bacterial species sourced from variable environments, notably many without previously characterized vanadium-dependent homologs. The previously reported effects of Br-AQs on environmental and pathogenic bacteria,^11,47–49^ combined with our observations of exogenous HHQ bromination in *Oleibacter marinus* cumulatively support that C3 AQ halogenation is a modification shared by many types of microbes to alter the impact of AQ toxicity and signaling. Previous literature investigations into the *in vitro* or *in situ* bromination of HHQ and HQNO suggest that this modification can begin the process of AQ degradation. While our study focuses on the formation of these halogenated metabolites instead of their decomposition, additional efforts are underway to further understand this unusual mechanism for QQ. The reported effects of (halogenated) AQ molecules on algal blooms and other marine ecosystems reinforce the idea that these molecules are being produced, modified, and degraded in variable environments.^50–53^ In our study, we saw alternative responses by three different bacteria to AQs produced by *Pseudomonas aeruginosa*, implying a deeper complexity to how environmental organisms interact with this family of molecules. Additional study of how these microbes and their environmental co-localizers respond to AQ C3-bromination will provide insight into QQ mechanisms utilized by bacteria, how these VHPO-induced modifications affect their microbial communities, and give additional perspective into underexplored AQ degradation strategies.

Our study reveals that AQ-VHPOs represent a novel subfamily of site-selective VHPO with heterogeneous phylogeny, broad environmental distribution, implied roles external to the cell, and intriguing physiological activities that set them apart from previously characterized macroalgal, fungal and streptomycete VHPOs. This work further expands the defined roles that bacterial VHPOs play beyond natural product biosynthesis^5–7^ and haloform production.^54^ Despite improvements in our understanding of their diverse ecological roles, their exciting selective-transformations, and their high biocatalytic compatibility, much about the functionality, distribution, and substrate profile of site-selective VHPOs remains unknown.^1,2,55^ Efforts to better understand these enzymes have been historically limited by their perception as solely sources of diffusible hypohalous acid, despite early work indicating they catalyze enantiospecific reactions.^10^ Structural insights on the naphthoquinone-derived meroterpenoid homologs, NapH1 and NapH3, have identified underlying features of actinobacterial VHPOs that differentiate them from their non-selective counterparts.^3^ Additionally, recent efforts with fungal and cyanobacterial VHPOs have begun to unveil modes of preferential substrate halogenation in VHPOs historically characterized as non-selective.^56,57^ Our characterization of AQ-VHPO activity provides a new perspective to better understand the enzymology of site-selective VHPOs. To that end, we are leveraging their unique biochemical features to interrogate the mechanisms of selectivity and general functionality of this enzyme family.

## Supporting information

AQ-VHPO-SI-PDF

## Supporting Information

- Experimental protocols, sequence similarity networks, crude lysate HHQ bromination MS chromatograms, AQ-VHPO sequence alignment, signal peptide identities, characterized VHPO phylogenetic tree and signal peptides presence, SDS-PAGE gel, monochlorodimedone assay and other purified AQ-VHPO substrate dependency assays, growth curves for bacteria in the presence of AQs and Br-AQs, *in vivo* MS chromatogram for *O. marinus* Br-HHQ production, optical images for *P. aeruginosa* co-cultures, additional MALDI-MSI images and MS spectra, *P. aeruginosa* swarming motility, synthetic procedures for HHQ and X-AQs, and tables of all bioinformatically identified AQ-VHPO, biosynthetic gene clusters for assembled genomes of AQ-VHPO organisms, DNA and amino acid sequences for heterologously expressed AQ-VHPOs, PCR primers, predicted signal sequences, VHPO accession numbers for phylogenetic tree construction, and halogenase tables for AQ-VHPO organism genome queries (PDF).
- FAIR data, including primary NMR FID files, qPCR ΔΔCt values, and MALDI-MSI raw spectral data.

## Acknowledgements

We gratefully acknowledge financial support from the National Institutes of Health (R35-GM147235-01 to S. M. K. M), National Science Foundation grant (IOS-2220510 to L. M. S.), Allen Distinguished Investigator Award, a Paul G. Allen Frontiers Group advised grant of the Paul G. Allen Family Foundation (L. M. S.), and the University of California, Santa Cruz for startup funding (S. M. K. M. and L. M. S). We appreciate the contributions of H.-W. Lee, J. MacMillan, and B. Rabbits (University of California, Santa Cruz) for assistance and maintenance of nuclear magnetic resonance, high-performance liquid chromatography, and plate-reader instrumentation, respectively. We acknowledge B. Dreyer (University of California, Santa Cruz Plasma Analytic Laboratory, RRID: SCR_02195) for assistance with and preparation of ICP-MS samples. We also would like to acknowledge Prof. Lars Dietrich (Colombia University) for supplying *P. aeruginosa* PA14 *pqs* genetic deletion strains.

## Author Contributions

J.T.B. and S.M.K.M conceived the project. J.T.B designed all experiments, analyzed data, and compiled the figures. C.S.M and L.M.S. helped with the preparation and interpretation of MALDI-MSI experiments and *P. aeruginosa* experiments. H.S.F. and A.R.L. helped purify proteins and generate data for proof-of-concept experiments. J.T.B. and S.M.K.M. wrote the manuscript with input from all authors.

## Notes

### Competing Interest Statement

The authors have declared no competing interest.

