## Supplementary material for "Vanadium-dependent haloperoxidases from diverse microbes halogenate exogenous alkyl quinolone quorum sensing signals": AQ-VHPO-SI-PDF

**This PDF file includes:**

Methods

Figures S1 to S49

Chemical Synthesis

Tables S1 to S9

References

**Table of Contents**


1. **Methods and Experimental** **S4-S13**

General Information S4
Bioinformatic and Genomic Methods S6
Biochemical Methods S7
Bacterial *in vivo* Methods S10

2. **Supplementary Figures** **S14-S63**

Figure S1. HZ11-VHPO Sequence Similarity Networks S14
Figure S2. Taxonomic and environmental sources of putative AQ-VHPO containing organisms S15
Figure S3. General HHQ biosynthetic pathway S16
Figure S4. HZ11-VHPO crude lysate HHQ bromination activity S17
Figure S5. esVHPO crude lysate HHQ bromination activity S18
Figure S6. lpVHPO crude lysate HHQ bromination activity S19
Figure S7. moVHPO crude lysate HHQ bromination activity S20
Figure S8. mrVHPO crude lysate HHQ bromination activity S21
Figure S9. A4B17-VHPO crude lysate HHQ bromination activity S22
Figure S10. omVHPO crude lysate HHQ bromination activity S23
Figure S11. plVHPO crude lysate HHQ bromination activity S24
Figure S12. SA03-VHPO crude lysate HHQ bromination activity S25
Figure S13. raVHPO crude lysate HHQ bromination activity S26
Figure S14. thVHPO crude lysate HHQ bromination activity S27
Figure S15. trVHPO crude lysate HHQ bromination activity S28
Figure S16. vbVHPO crude lysate HHQ bromination activity S29
Figure S17. vnVHPO crude lysate HHQ bromination activity S30
Figure S18. Environmental distribution of confirmed AQ-brominating VHPOs S31
Figure S19. Sequence alignment of AQ-VHPOs S32
Figure S20. Conservation of secretion signals across all AQ-VHPOs S33
Figure S21 Phylogenetic distinction between the presence and absence of signal peptides in characterized VHPOs S34
Figure S22. SDS-PAGE of purified AQ-VHPOs S35
Figure S23. MCD assay of AQ-VHPOs demonstrates absence of diffusible hypobromous acid production S36
Figure S24. Co-substrate and co-factor dependency of purified AQ-VHPOs S37
Figure S25. lpVHPO control of MHQ bromination S38
Figure S26. lpVHPO halide oxidation profile S39
Figure S27. raVHPO halide oxidation profile S40
Figure S28. omVHPO halide oxidation profile S41
Figure S29. Response of *V. natriegens* CCUG 16374 to exogenous HHQ and Br-HHQ S42
Figure S30. Response of *M. rhizosphaerae* to exogenous HHQ and Br-HHQ S42
Figure S31. Response of *O. marinus* to exogenous MHQ and Br-MHQ S43
Figure S32. *In vivo* biotransformation of exogenous HHQ by *O. marinus* S44
Figure S33. *O. marinus* omVHPO qPCR S45
Figure S34. *O. marinus* growth curves for all HHQ concentrations in each media condition S46
Figure S35. Optical images of *O.marinus* and *P. aeruginosa* PA14 co-cultures used for MALDI-MSI S47
Figure S36. Additional experiments for *O. marinus* MSI S48
Figure S37. Analysis of contribution to $^{79}$Br-HHQ MALDI mass ion intensity by competing background peaks S49
Figure S38. Specific MS1 Spectra chosen for MALDI imaging of Figure 5d and Figure S35ab S50

Figure S39. Specific MS1 Spectra chosen for MALDI imaging of Figure 5e S51
Figure S40. Specific MS1 Spectra chosen for MALDI imaging of Figure 5fg S52
Figure S41. Optical images of *V. natriegens* and *P. aeruginosa* PA14 co-cultures used for MALDI-MSI S53
Figure S42. Communal interaction between *V. natriegens* and *P. aeruginosa* PA14 S54
Figure S43. Specific MS1 Spectra chosen for MALDI imaging of Figure S42 (Panels a and b) S55
Figure S44. Specific MS1 Spectra chosen for MALDI imaging of Figure S42 (Panels c and d) S56
Figure S45. Optical images of *M. rhizosphaerae* and *P. aeruginosa* PA14 co-cultures used for MALDI-MSI S57
Figure S46. Communal interaction between *M. rhizosphaerae* and *P. aeruginosa* PA14 S58
Figure S47. Specific MS1 Spectra chosen for MALDI imaging of Figure S46 S59
Figure S48. Optical images of *P. aeruginosa* PA14 swarming motility S60
Figure S49. *P. aeruginosa* PA14 swarm areas S62

3. **Chemical Synthesis** **S63-S67**
Synthesis of 2-heptyl-4(1H)-quinolone (HHQ) S63
Synthesis of 2-heptyl-3-bromo-4(1H)-quinolone (Br-HHQ) S64
Synthesis of 2-methyl-3-chloro-4(1H)-quinolone (Cl-MHQ) S65
Synthesis of 2-methyl-3-bromo-4(1H)-quinolone (Br-MHQ) and 2-methyl-3,6-dibromo-4(1H)-quinolone ($Br_2$-MHQ) S66
Synthesis of 2-methyl-3-iodo-4(1H)-quinolone (I-MHQ) S67

4. **Tables** **S68-S105**
Table S1. All putative alkyl quinolone modifying VHPOs and their source organisms S68
Table S2. Biosynthetic potential of alkyl quinolone VHPO containing strains S74
Table S3. *E. coli* codon optimized DNA sequences for selected alkyl quinolone VHPOs S77
Table S4. Amino acid sequences for selected alkyl quinolone VHPOs S87
Table S5. Primers used in this study S91
Table S6. Predicted secretion signal peptides S96
Table S7. Representative VHPO accession numbers from across kingdoms S102
Table S8. Halogenases used to query bacterial genomes for halogenation potential S104
Table S9. Halogenases in *Oleibacter marinus*, *Vibrio natriegens*, and *Microbulbifer rhizosphaerae* S105

5. **References** **S106-S107**

### Methods and Experimental

#### General Information

##### Materials and instruments

All chemicals, media, and solvents used were purchased from commercial suppliers (Acros, Sigma Aldrich, Difco, Fisher, Oakwood, Cayman Chemical Company, or Alfa Aesar) and used as received unless otherwise noted. All tested AQs were purchased as commercial standards. HHQ, Br-HHQ, Cl-MHQ, Br-MHQ, and I-MHQ were prepared synthetically (Chemical Synthesis Section). PCR was run using a Mastercycler nexus – PCR Thermal Cycler (Eppendorf). All FPLC-based protein purification was performed on an ÄKTAGo instrument (GE Healthcare) equipped with buffers filtered through a 0.22 μM nitrocellulose membrane. FPLC data was analyzed with UNICORN v 7.5 (GE Healthcare). Centrifugations were performed with a Centrifuge 5424 R (Eppendorf) (FA-45-24-11 rotor), a Sorvall Lynx 6000 (Thermo) (Fiberlite F9-6x1000 Lex rotor), and/or a Centrifuge 5804 R (Eppendorf) (S-4-72 rotor or FA-45-6-30 rotor). A BioSpectrometer kinetic (Eppendorf) and a NanoDrop One$^{C}$ (Thermo) were used to collect UV-vis and optical density ($OD_{600}$) readings. Solid media bacterial cultures were grown in a Heratherm Compact Microbiological Incubator (Thermo) and liquid media bacterial cultures were grown in a New Brunsweick Scientific Excella E24 Shaker (Eppendorf) or a New Brunswick Inova44 (Eppendorf).

##### Bacteria strain acquisition and growth conditions

Plasmid propagation and heterologous expression were performed in *Escherichia coli* DH10β competent cells (Invitrogen) and in *E. coli* BL21(DE3) competent cells (New England Biolabs), respectively. *E. coli* was cultured in either Lysogeny Broth (LB) or Modified Terrific Broth (TB) (Fisher). *Pseudomonas aeruginosa* PA14 WT and its genetic knockouts (ΔpqsA-C, and ΔpqsH) used in this study were prepared previously.[1] *P. aeruginosa* PA14 was cultured in LB media, except where otherwise noted. *Oleibacter marinus* (DSM 24913)[2] *and Microbulbifer rhizosphaerae* (DSM 28920)[3] were obtained from the German Collection of Microorganisms and Cell Cultures (DSMZ). *Vibrio natriegens* CCUG 16374 (ATCC 33788)[4] was obtained from the American Type Culture Collection (ATCC). All environmental strains were revived and cultured precisely with the handling instructions and media conditions provided by the strain repositories, unless otherwise noted. All liquid media cultures were shaken at 200 RPM. All media was sterilized in a SX-500 autoclave (TOMY).

**Caution!** *P. aeruginosa* PA14 was handled using BSL2 protocols.

##### Analytical Methods

NMR Spectra were collected using an Avance III HD Spectrometer (Bruker) equipped with a BBFO SmartProbe at 500 MHz ($^{1}$H NMR) or 125 MHz ($^{13}$C NMR) using MeOD and $CDCl_3$ as solvents. Chemical shifts are reported in ppm and referenced to the CDCl3 solvent signal (δ = 7.26 ppm for $^{1}$H, δ = 77.2 ppm for $^{13}$C NMR. NMR splitting data are reported as follows: singlet (s), doublet (d), triplet (t), quartet (q), pentet (p), multiplet (m), coupling constant (*J*, Hz). Synthetic reaction progress was monitored via LCMS and/or precoated thin-layer chromatography (TLC, Merck, 60 $F_{24}$). High resolution mass spectrometer (HRMS) data were collected via MALDI-QqTOF on a TIMSTOFfleX (Bruker) in positive mode with a laser power of 35%, frequency of 1000 Hz, and 2000 shots per sample. A 1:1 mixture of recrystallized δ-cyano-4-hydroxycinnamic acid (CHCA) and 2,5-dihydroxybenzoic acid (DHB) (Sigma) was used as MALDI matrix and Phosphorous Red (Sigma) was used as calibrant. UPLC-MS data was collected on an Elute UPLC instrument (Bruker) coupled to an Amazon ESI-ion trap mass spectrometer (Bruker). Compounds were separated in reversed-phase chromatography using 0.1% formic acid (Solvent A) and acetonitrile + 0.1% formic acid (solvent B) with a Bruker Intensity Solo C18(2), 2 mm x 100 mm column. The following LC separation method was used: 10% to 20% solvent B over 3 minutes, 20% to 100% solvent B over 3 minutes, 100% solvent B over 2 minutes, 100% to 90% solvent B over 1 minute, 90% to 10% solvent B over 2 minutes. All MS data was collected in positive mode. LC components were controlled with Compass Hystar 5.1 (Bruker), MS was controlled with ESI Compass trapControl v 8.0 (Bruker), and LCMS data was analyzed with Compass DataAnalysis v 5.2 (Bruker). For normalization of alkyl quinolone ion peak areas, an internal standard of 250 nM nalidixic acid was added to all MS samples.[5] Calibration curves for HHQ, Br-HHQ, MHQ, and Br-MHQ were generated to confirm that there was no significant difference in ionization efficiency between AQs and their C3 brominated derivatives. Commercial AQs and synthesized Br-HHQ and Br-MHQ were used as standards. The reaction outputs of the established C3-bromination enzyme HZ11-VHPO with NHQ, DHQ, and HQNO were used as standards for bromination activity on said substrates. Ion peak areas were collected by extracting the relevant $[M+H]^+$ ion for each AQ species; for chlorinated and brominated species the M and M+2 peaks were extracted for. All chromatograms were analyzed in BPC mode to confirm that side reactions or additional products were not being produced. All mass spectrum chromatograms were exported to InkScape for graphical workup. All protein quantification was performed with the Bradford method using Quick Start Bradford 1x Dye Reagent (Bio-Rad Laboratories) and a standard curve generated with Bovine Serum Albumin (Thermo Scientific). All data was collected at least in duplicate.

**General Biological Methods**

Plasmids were transformed into competent *E. coli* strains with the following protocol. Chemically competent *E. coli* 100 µL aliquots were removed from -70 °C and thawed on ice. Plasmids were directly pipetted into the aliquot (1 µL of pure 40-60 ng/µL plasmid) and then gently tapped to mix. The mixture was then incubated on ice for 30 minutes before being placed at 42 °C for 45 seconds. After heat shock, cells were kept at room temperature for 2 minutes before transferring to 700 µL LB media and shaking for 1 hour at 37 °C. 200 µL of the cells were then plated on LB Agar supplemented with 50 µg/mL Kanamycin. Plates were incubated for 16-20 hours at 37 °C. Single colonies were selected for all future applications.

Protein was expressed with the following protocol except where otherwise noted. Freshly transformed *E. coli* BL21(DE3) containing the plasmid of interest was inoculated into LB media (either 5 mL or 25 mL, depending on the scale of expression) supplemented with 50 µg/mL Kanamycin and incubated for 16-20 hours at 37 °C to generate starter cultures. TB media supplemented with 50 µg/mL Kanamycin was then inoculated with 1% v/v of the starter culture. Cultures were then incubated at 37 °C until $OD_{600}$ reached 0.5-0.7 (typically 4-6 hours). Cultures were then placed at 18 °C for 1 hour and then 100 µM isopropyl-β-D-thiogalactopyranoside (IPTG) was added to induce protein expression. Cultures were then incubated for 20 hours at 18 °C before harvesting. Cultures were harvested by centrifuging at 2500 x g for 30 minutes at 4 °C. Supernatant was poured off and the pellets were re-suspended in 30 mL Buffer A per liter of culture (50 mM Tris-HCl pH 8.0, 20 mM Imidazole, 500 mM NaCl). This sub-family of VHPOs do not appear to consistently purify or display activity when frozen before purification so all purifications or other lysate applications were performed on the day of harvest.

**Enzymatic Assay Components and Stocks**

All AQ substrate stocks (HHQ, MHQ, DHQ, NHQ, HQNO) were prepared as 10 mM stocks in DMSO and stored at -20 °C. 10 mM $Na_3VO_4$, 2 M KBr, 2 M KCl, and 2 M KI stocks were prepared in MilliQ water and stored at 4 °C. 10 mM $H_2O_2$ stocks were prepared fresh for each assay from a 30% $H_2O_2$ solution stored at 4 °C. Reactions were ran in the following buffer conditions: 20 mM HEPES-KOH pH 8.0 and 10% glycerol. Standard reactions contained 100 µM $Na_3VO_4$, 20 mM halide, 1 mM $H_2O_2$, 100 µM AQ substrate (equates to 1% DMSO in all reactions), and 2.5 µM VHPO. All reaction components, aside from enzyme, were allowed to warm to room temperature before initiating reactions. Typical reactions were initiated with the addition of $H_2O_2$. All reactions were incubated at 28 °C and 200 RPM shaking. Reactions were run at 100 µL scale and quenched with an equal volume of methanol. Before preparing MS samples, reactions would be centrifuged for 5 minutes at 11000 x g to remove precipitated protein. Reactions would be further diluted for MS to avoid ion suppression, when necessary. Nalidixic acid internal standard was added after quenching or after all subsequent dilutions had been performed.

#### Bioinformatic and Genomic Methods

##### EFI-EST Sequence Similarity Network Generation

The sequence similarity network was produced using Enzyme Function Initiative-Enzyme Similarity Tool (EFI-EST).[6,7] NCBI-BLAST was used to collect the top 500 most similar sequences to HZ11-VHPO. Accession IDs were batch downloaded and used in the "Accession IDs" option with all default options maintained. Networks were generated from alignment thresholds of 130, 140, and 150. The generation of previously observed clusters and sub-clusters aside from the putative AQ-VHPO cluster ensured that the collection of 500 sequences was sufficient. Networks were visualized in Cytoscape 3.5.1 and Organic yFiles were used to visualize the networks.[8]

##### Phylogenetic analysis of putative AQ-VHPOs

Putative AQ-VHPO accession numbers and sequences were extracted from the SSN clusters associated with HZ11-VHPO. Redundant and incomplete sequences were culled from the accession list, resulting in 75 unique putative AQ-VHPOs (including HZ11-VHPO) (Table S1). The source organism, status of genomic assembly (assembled vs. metagenomic), and environmental source were acquired from NCBI. A Pap2 phosphatase family enzyme (Uniprot ID: P0A924) from *E. coli* K12 was selected to act as an outgroup. Sequence alignment and phylogenetic tree generation was performed in Mega11.[9] Sequences were aligned in the program using built-in default ClustalW settings. Phylogenetic analysis was then performed on the alignment using Maximum Likelihood as the statistical method, 100 bootstraps, the Jones-Taylor-Thornton (JTT) model, using all sites for Gap/Missing data treatment, and the Nearest-Neighbor-Interchange heuristic method. The tree was exported to and enhanced in Adobe Illustrator to improve image quality and visual comprehension.

##### Phylogenetic analysis of characterized VHPOs from across kingdoms

Amino acid sequences of characterized VHPOs were acquired from NCBI, UniProt, or JGI IMG (Table S7). The VHPOs selected for this include a sample of putative AQ-VHPOs representing the diversity of taxa, the characterized actinobacterial meroterpenoid VHPOs, and non-selective VHPOs from flavobacteria, fungi, brown algae, red algae, and cyanobacteria. A Pap2 phosphatase family enzyme (Uniprot ID: P0A924) from *E. coli* K12 was selected to act as an outgroup. The tree was generated as above.

##### Genomic context analysis of AQ-VHPOs

The genomes of selected strains were accessed through DOE JGI-IMG or NCBI. Genbank files of the VHPO genomic context were generated in SnapGene Software (www.snapgene.com). Genomic contexts were visualized as 10 coding genes upstream and downstream of the VHPO. Genomic neighborhoods were aligned using Clinker.[10] COG definitions were used to attribute the likely cell functions of each gene, aside from the VHPO.[11] Essential cellular functions involve genes related to primary metabolism, replication and division, cellular structure, and translation. Secondary metabolism or Defense involves genes related to natural product biosynthesis, anti-toxins, or other protective molecule production. External modification or response involves genes related to signaling, transcription factors, quorum sensing, or secreted molecules/biomolecules. Mobilosome or recombination related genes involves genes related to integrases, prophage proteins, heterologous recombination, or other mobile genetic elements. Hypothetical proteins involved genes with unknown function, no COG definition, or chance coding sequences. All strains for which assembled genomes were available (Table S2) were analyzed initially by antiSMASH.[12] The *pqsABCDE* protein sequences from *P. aeruginosa* PA14 and from *Microbulbifer* sp. HZ11 were used to query all assembled genomes for the presence of AQ biosynthetic machinery using NCBI BLAST.

##### Sequence alignment and structural mapping of AQ-VHPOs

Amino acid sequences for activity tested AQ-VHPOs (Table S4) were aligned with the default settings of Clustal Omega[13] and subsequently visualized with ESPript 3.0.[14] The AlphaFold model of esVHPO was generated via ColabFold v1.5.5[15] and aside from relaxing the top seed, all default settings were selected. The top selected seed was chosen as the model for esVHPO. The sequence alignment of the AQ-VHPOs was mapped to the esVHPO AlphaFold model using ConSurf server.[16]

#### Biochemical Methods

##### Heterologous AQ-VHPO Cloning

The VHPO sequences selected for heterologous expression and *in vitro* activity testing are listed in Table S3**.** The VHPO sequences were codon optimized for *E. coli* expression (Table S4 and purchased (Twist Bioscience). The purchased genes were then amplified by PCR using primers (Table S5) with Gibson overhangs designed for NdeI/XhoI insertion (N-terminal hexahistidine tag) or NcoI/XhoI insertion (C-terminal hexahistidine tag) into pET28a(+) backbone. Genes were amplified with Prime Star HS DNA Polymerase (TaKaRa) using manufacturer supplied buffers/reagents, 200 nM primers, 1 ng/25 µL gene template, and 3% DMSO. The following two step PCR thermocycling method was used to amplify gene inserts: initial denaturation at 98 °C (120 seconds), 35 cycles of melt at 98 °C (10 seconds) and anneal/elongate at 72 °C (90 seconds), final elongation at 72 °C (120 seconds). Gene inserts were then purified through gel extraction using a QIAquick Gel Extraction kit (Qiagen). pET28a(+) was linearized for N-terminal hexahistidine constructs by PCR using the primers in Table S5 and Prime star Max DNA Polymerase 2x master mix (TaKaRa), 200 nM primers, 1 ng/25 µL gene template, and 3% DMSO. The following PCR thermocycling method was used to amplify linear backbone: initial denaturation at 98 °C (120 seconds); 35 cycles of melt at 98 °C (10 seconds); anneal at 64.4 °C (5 seconds), elongate at 72 °C (150 seconds); final elongation at 72 °C (120 seconds). Linear backbone was then purified through gel extraction using a QIAquick Gel Extraction kit (Qiagen). pET28a(+) vector was linearized for C-terminal hexahistidine constructs by restriction digest of empty pET28a(+) plasmid with NcoI and XhoI (New England Biolabs) at 37 °C (30 minutes) and then at 80 °C (20 minutes). Digested vector was then worked up with QIAquick PCR Purification Kit (QIAgen). Gene inserts and linearized pET28a(+) vector were ligated using NEBuilder HIFI DNA Assembly Master Mix (New England Biolabs), using 50 ng of vector at a 2:1 molar ratio of gene insert:vector. Gibson assembly mix was incubated at 50 °C for 30 minutes and then placed on ice for subsequent use. 10 µL of Gibson reaction was transformed into chemically competent *E. coli* DH10β. Individual colonies were then used to inoculate 5 mL LB media supplemented with 50 µg/mL Kanamycin and incubated for 16-20 hours at 37 °C. Overnight cultures were then centrifuged at 2500 x g for 30 minutes at 4 °C and the supernatant discarded. Plasmids were then purified from the pellet using the manufacturer instructions from QIAprep Spin Miniprep Kit (QIAgen). Purified plasmids were confirmed to have the correct VHPO construct by Sanger sequencing (Azenta Life Sciences).

##### Crude Lysate Preparation and Assay Conditions

The VHPOs listed in Table S4 and empty pET28a(+) were cultured at small scale for the detection of AQ-bromination activity. The general expression protocol was followed on a 50 mL TB scale. After centrifugation and the removal of the supernatant, cell pellets were re-suspended in 500 µL of Buffer A. Resuspended pellets were sonicated on ice (Model 50 Sonic Dismembrator (Fisherbrand), 1/8 in. probe, 40% amplitude, 10 s pulse on/10 s pulse off, 1 minute “on” time). Crude lysates were then immediately tested for bromination activity. 100 µL reactions with standard assay components, KBr as the halide source, and HHQ as the AQ substrate were prepared. 10% v/v crude lysate was added to each reaction. Reactions were incubated at 28 °C and 200 RPM for 20 hours. Reactions were then quenched and prepared for LCMS analysis. A small amount of spontaneous bromination was detected in the empty vector control, so a two-fold greater degree of bromination was set as the threshold for activity (roughly equivalent to a log scale ratio of 0.25).

##### Heterologous AQ-VHPO Expression and Purification

The general expression method was followed for: HZ11-VHPO, esVHPO, IpVHPO, and raVHPO. Each VHPO was expressed on a 2 L TB media scale. Resuspended pellets were sonicated on ice (Q500 Sonicator (QSonica), 6.4 mm probe, 40% amplitude, 15 s pulse on/45 s pulse off, 5 minutes “on” time). The lysate was then centrifuged for one hour at 16,000 x g and 4 °C or until supernatant had clarified. Clarified supernatant was loaded onto a 5 mL HisTrap FF Column (GE Healthcare Life Sciences) that was pre-equilibrated with at least 5 CV of Buffer A at a flow rate of 2 mL/min. After all clarified supernatant was loaded, column was rinsed with buffer A until UV absorbance had returned to baseline. The column was washed with 10% Buffer B (50 mM Tris-HCl pH 8.0, 500 mM Imidazole, 500 mM NaCl) for at least 5 CV or until UV absorbance had returned to baseline. Hexahistidine tagged protein was eluted with a linear gradient from 10% Buffer B to 100% Buffer B over 60 mL, while collecting 5 mL fractions. Fractions were assessed for purity through SDS-PAGE (10% acrylamide) and were combined if at least 90% pure. Combined fractions were concentrated to 2.5 mL using a 30 kDa cutoff Amicon Ultra-15 concentrator. Protein was buffer exchanged into Cold Storage Buffer (25 mM HEPES-KOH pH 8.0, 300 mM KCl, 10% glycerol) using a pre-equilibrated PD-10 gravity flow column (GE Healthcare Life Sciences). Protein concentration was then quantified with Bradford reagent and further concentrated if necessary. Each protein was stored at -70 °C for further purification. Each VHPO was purified using a HiLoad 16/600 Superdex 200 pg (GE Healthcare Life Sciences) column using GF Buffer (20 mM HEPES pH 8.0 and 300 mM KCl). Each VHPO eluted at a retention time that corresponded to the molecular weight of the dimer. esVHPO (57 kDa, 5 mg/L) solubly purified as an N-terminal hexahistidine tagged construct. lpVHPO (51 kDa, 2 mg/L) and raVHPO (52 kDa, 1.5 mg/L) solubly purified as C-terminal hexahistidine tagged constructs with cleaved N-terminal signal sequences, based off SDS-PAGE migration (Figure S17). HZ11-VHPO did not elute in the Buffer B gradient fractionation but was observed as a major component of the 10% Buffer B wash. The wash fraction was collected, de-salted, and further purified with gel filtration as above. The eluted protein was then purified a second time with a 10/300 Superdex 200 Increase (Cytiva) to about 50% homogeneity (0.10 mg/L).

omVHPO and thVHPO appeared to be secreted by the heterologous system so their expression and purification protocol was modified. omVHPO was grown at 6 L scale, while thVHPO was grown at 1 L scale. The general expression protocol was followed until the induction stage. Once induced, cultures were shaken at 200 RPM and 18 °C for 40 hours. Cultures were harvested by centrifuging at 2500 x g for 30 minutes at 4 °C. Supernatant was retained and the pellets were discarded. 5 mL of batch Ni-NTA resin (GE Healthcare Life sciences) pre-equilibrated in Buffer A was added to the supernatants and was incubated for 1 hour at 50 RPM and 4 °C. Supernatant was then passed through an empty 1 x 50 cm Econo-Column (Bio-Rad Laboratories) and the resin collected on the frit. Once all clarified supernatant had been passed through, resin was washed with Buffer A until no more protein eluted (detected by reaction with Bradford reagent). Resin was then washed with 5 CV 10% Buffer B or until no more protein eluted. Resin was then eluted with 5 CV of 100% Buffer B to collect hexahistidine tagged VHPO. Fractions were assessed for purity through SDS-PAGE (10% acrylamide). From this point, the protocol returned to the general purification protocol. omVHPO (52 kDa, 0.25 mg/L) and thVHPO (53 kDa, 0.1 mg/L) solubly purified as C-terminal hexahistidine tagged constructs with cleaved N-terminal signal sequences, based off SDS-PAGE migration (Figure S17) and their localization to the culture media.

**Monochlorodimedone (MCD) Assay of esVHPO and HZ11-VHPO**

The conditions for the MCD assay were adapted from previous work on selective VHPOs.[17] Reactions were performed in a quartz cuvette and monitored at 290 nm for 30 minutes with the absorbance recorded every 10 seconds. For each reaction, the spectrophotometer was blanked with the following mixture: 50 mM HEPES-KOH pH 8.0, 200 mM KCl, 200 mM KBr, 1 mM $H_2O_2$, 10 µM $Na_3VO_4$, and 10 µg/mL VHPO. For each VHPO, 1 mL reactions containing 50 mM HEPES-KOH pH 8.0, 50 µM MCD, 200 mM KCl, 10 µM $Na_3VO_4$, and 10 µg/mL VHPO were prepared. To establish a baseline reading, reaction was monitored for 2 minutes. Then, 1 mM $H_2O_2$ was added to assess for possible non-selective chloroperoxidase activity. After 5 minutes, 200 mM KBr was added to the reaction and monitored until 30 minutes was reached. The same reaction was performed at pH 6.0 using 50 mM MES pH 6.0 in place of HEPES-KOH pH 8.0. The selective chloroperoxidase NapH1, which diffuses hypobromous acid but not hypochlorous acid, was used as a positive control.[18]

**AQ substrate preferences of AQ-VHPOs**

Purified HZ11-VHPO, esVHPO, lpVHPO, raVHPO, and omVHPO were tested against a panel of 5 AQ substrates: HHQ, MHQ, NHQ, HQNO, and DHQ. 100 µL reaction mixes were prepared including all reaction components except for the AQ. Reactions were initiated by the addition 1 µL of the appropriate AQ. Reactions were incubated for 30 minutes at 28 °C before quenching and preparing for LCMS analysis. 10-fold dilutions were performed to avoid ion suppression effects. Relative conversion percentages were calculated as a ratio of Br-AQ peak area to AQ peak area added to the Br-AQ peak area, after normalization against the internal standard. Bar charts were prepared in GraphPad Prism version 8.0.0 for windows.

**Co-substrate and Co-factor dependency of AQ-VHPOs**

For each VHPO, 5 100 µL reactions with standard assay components were prepared with the following alterations: "No enzyme" omitted the addition of 2.5 µM VHPO, "No $H_2O_2$" omitted the addition of 1 mM $H_2O_2$, "No Added $Na_3VO_4$" omitted the addition of 100 µM $Na_3VO_4$, "No KBr" omitted the addition of 20 mM KBr, and "All substrates" was as standard conditions. All reactions utilized MHQ as the AQ substrate. Each condition was incubated for 30 minutes at 28 °C before quenching and LCMS analysis. Due to low purification yields, omVHPO and thVHPO assays were ran with 20 µM MHQ and 0.5 µM VHPO.

**ICP-MS of esVHPO and raVHPO**

esVHPO and raVHPO were digested with pancreatic trypsin (Sigma). 100 µg of lyophilized trypsin was resolubilized in 1 mM HCl to a concentration of 1 mg/mL. Prior to incubation with trypsin, both VHPOs were denatured by heating at 75 °C for 15 minutes in the following conditions: 100 mM Tris-HCl pH 8.0, 1 mM DTT, and 1 mg/mL VHPO. Denatured VHPOs were then brought to room temperature before adding 0.05 mg/mL trypsin (1:20 ratio of trypsin to VHPO) (total volume of 400 µL). Digest reactions were then incubated at 37 °C and 120 RPM for 20 hours. Reactions were then diluted to a volume of 3 mL with 1% $HNO_3$. Digests were centrifuged for 5 minutes at 3500 x g and supernatants were collected. Samples were analyzed at the UCSC Plasma Analytical Lab (RRID: SCR_02195) using an Element XR high-resolution ICP-MS (Thermo) instrument.

**Excess Hydrogen Peroxide Assays**

Purified esVHPO was reacted under standard conditions with MHQ as the AQ substrate, except for hydrogen peroxide concentration. Four separate concentrations of $H_2O_2$ were assayed: 50 µM $H_2O_2$ (0.5 stoichiometric equivalents of AQ substrate), 100 µM $H_2O_2$ (1 stoichiometric equivalent of AQ substrate), 500 µM $H_2O_2$ (0.5 stoichiometric equivalents of AQ substrate), and 1 mM $H_2O_2$ (10 stoichiometric equivalents of AQ substrate). The same assay was prepared for lpVHPO with the following concentrations of $H_2O_2$: 50 µM $H_2O_2$ (0.5 stoichiometric equivalents of AQ substrate), 200 µM $H_2O_2$ (2 stoichiometric equivalent of AQ substrate), 1 mM $H_2O_2$ (10 stoichiometric equivalents of AQ substrate), and 10 mM $H_2O_2$ (100 stoichiometric equivalents of AQ substrate). The reaction was incubated at 28 °C and 200 RPM for ten minutes before quenching and LCMS analysis.

**NBS bromination of MHQ**

This protocol was adapted from previous non-enzymatic bromination approximating enzymatic conditions.[19] MHQ (5 mg, 0.03 mmol) was dissolved into a solution of 2 mL tetrahydrofuran (THF) and 2 mL enzyme assay reaction buffer (20 mM HEPES-KOH, 10% glycerol). Four reaction conditions with different equivalents of recrystallized NBS were prepared with the following additions of NBS: 0 equivalents NBS (no NBS added), 1 equivalent NBS (5.3 mg, 0.03 mmol), 2 equivalents NBS (10.6 mg, 0.06 mmol), and 10 equivalents NBS (53 mg, 0.3 mmol). Reactions were secured to a 200 RPM shaker at 28 °C and reacted for 5 hours. Reactions were diluted 1:50 into methanol and analyzed by LCMS. Initial analysis was performed through comparison of the 0 equivalent NBS BPC chromatogram to the others and then extracted ion chromatograms were prepared based on ions only present in plus NBS conditions. The following ions representing their species were extracted for all chromatogram traces: MHQ (160.07 ± 0.1 m/z $[M+H]^+$), Br-MHQ (237.98, 239.98 ± 0.1 m/z

$[M+H]^+$), $Br_2$-MHQ (315.89, 317.89, 319.89 ± 0.1 m/z $[M+H]^+$), and $Br_3$-MHQ (393.80, 395.80, 397.80, 399.79 ± 0.1 m/z $[M+H]^+$).

**Halide specificity**

100 µL reactions with standard assay components were prepared. For each halide assay only one halide (KCl, KBr, or KI) was added. Each halide was added to the relevant assay at a concentration of 20 mM. Each halide condition was incubated in the reaction buffer with all relevant components aside from the VHPO to assess for spontaneous hypohalite production in the presence of hydrogen peroxide. Each VHPO was buffer exchanged into reaction buffer (20 mM HEPES-KOH pH 8.0 and 10% glycerol) prior to assay to ensure added buffer salts did not interfere. The reaction was incubated at 28 °C and 200 RPM for thirty minutes before quenching and LCMS analysis.

**Bacterial *in vivo* Methods**

**Environmental bacteria culture conditions**

*O. marinus* was cultivated in the following media: 3.7 g/L Marine Broth (Millipore), 5.0 g/L sodium pyruvate, and 1.5% agar (for solid media). The nutrient rich components were suspended in 250 mL MilliQ water per intended liter of media. 750 mL of artificial sea salt solution per intended liter of media was prepared with the following components: 21.2 g/L NaCl, 3.5 g $Na_3SO_4$, 0.6 g/L KCl, 0.175 g/L $NaHCO_3$, 0.9 g/L KBr, 23 mg/L boric acid, 3 g NaF, 9.6 g/L $MgCl_2·6\ H_2O$, 1.3 g/L $CaCl_2·6\ H_2O$, 22 mg/L $SrCl_2$, 0.1% 2.3 g/L $NaNO_3$, 0.1% 0.16 g/L $NaH_2PO4·H_2O$, 0.1% 0.09 g/L $FeCl_3·6\ H_2O$, 0.1% 0.28 g/L $Na_2EDTA·2\ H_2O$, 0.1% 3.6 mg/L $ZnSO_4·H_2O$, 0.1% 0.5 mg/L $CoSO_4·7\ H_2O$, 0.1% 27 mg/L $MnSO_4 · H_2O$, 0.1% 0.07 mg $Na_2MoO_4·2$ H2O, 0.1% 0.09 µg/L $Na_2SeO_3$, 0.1% 0.075 mg/L $NiCl_2·6\ H_2O$, 0.1% 5 mg/L Thiamine-HCl, 0.1% 0.1 mg/L Biotin, and 0.1% 0.05 mg/L Vitamin B12. The two mixtures were autoclaved separately and mixed prior to use to generate modified marine broth plus pyruvate media (MMBP media). Media for *V. natriegens* and *M. rhizosphaerae* were prepared similarly, with the exception of using 5.0 g/L of Marine Broth (Millipore) as the rich nutrient source (MB media). All cultures were prepared on 1.5 mL, 25 mL, or 50 mL scale depending on application. All cultures were grown at 28 °C and 200 RPM shaking (if applicable). Prior to its use in experiments, each strain would be inoculated from glycerol stock and grown to stationary phase (overnight for *V. natriegens* and *M. rhizosphaerae*; 3 days for *O. marinus*). This starter culture would then be used to inoculate additional cultures used in experiments. *O. marinus* viability in the presence of 10 nM, 20 nM, 30 nM, and 40 nM added $Na_3VO_4$ was tested and found to be tolerant of 10 nM $Na_3VO_4$ but was less tolerant for concentrations above that. *O. marinus* tolerance to additional KBr was tested and was found to be viable in concentrations excess of 20 mM KBr. To maintain parity between strains and culture conditions, 10 nM $Na_3VO_4$ and 20 mM KBr were used when relevant for all strains. Vanadate and additional KBr were not added as a part of routine culture.

**Br-HHQ extraction protocol cultures with added HHQ**

Media for each strain was prepared at 25 mL scale and then DMSO, 1 µM, 10 µM, and 25 µM HHQ was added to separate preparations. 10 nM $Na_3VO_4$ and 20 mM KBr were added to each culture. Bacteria were inoculated into the cultures at a 1:10 dilution from starter cultures. 1 mL aliquots were taken every 24 hours, quenched with an equal volume of methanol, and analyzed by LCMS. Once a culture had reached stationary phase, it would be incubated for another 24 hours and then inactivated with an equal volume of ethyl acetate. The organic layer would then be removed, and the aqueous phase washed two more times ethyl acetate. Organic solvent was removed *in vacuo* and metabolites were re-suspended in methanol to be analyzed on UPLC-MS.

***O. marinus* response to HHQ, Br-HHQ, MHQ, and Br-MHQ**

50 mL MMBP media was inoculated from an *O. marinus* starter culture and grown to an $OD_{600}$ of 0.5. This culture was then diluted in fresh MMBP media to an $OD_{600}$ of 0.05. 1.5 mL aliquots of this suspension were transferred to a sterile 24 well tissue culture plate (Falcon). AQs were added to intended final concentrations by the addition of 15 µL AQ from 100x stock solutions in DMSO. The vehicle control was prepared by adding 15 µL

DMSO to the appropriate wells. The plates were sealed with Breathe-Easy sealing membranes (Sigma) to prevent contamination between wells. Plates were secured to a 200 RPM shaker at 28 °C and incubated for 7 days. Regular measurements of the $OD_{600}$ were recorded with an EnVision 2105 multimode plate reader (Perkin Elmer). Each replicate was blank corrected against sterile media wells with the same concentration of AQ. The growth curves for each concentration of AQ were normalized to the DMSO growth curve with the following equation:

$$\% \text{ Normalized } OD_{600} = ({}_{[AQ]}OD_{600} - {}_{DMSO}\text{Minimum}_{OD600})/({}_{DMSO}\text{Maximum}_{OD600} - {}_{DMSO}\text{Minimum}_{OD600}) * 100.$$

The response to different concentrations of AQs was then defined as the percentage $OD_{600}$ absorbance of the vehicle control. Data plots were prepared in GraphPad Prism version 8.0.0 for Windows. Lines connecting the points were added for visual clarity by Spline fitting and do not represent a line of best fit.

##### *V. natriegens* response to HHQ and Br-HHQ

50 mL MB media was inoculated from a *V. natriegens* starter culture and grown to an $OD_{600}$ of 0.5. This culture was then diluted in fresh MB media to an $OD_{600}$ of 0.05. 1.5 mL aliquots of this suspension were transferred to a sterile 24 well tissue culture plate (Falcon). AQs were added to intended final concentrations by the addition of 15 µL AQ from 100x stock solutions in DMSO. The vehicle control was prepared by adding 15 µL DMSO to the appropriate wells. The plates were sealed with Breathe-Easy sealing membranes (Sigma) to prevent contamination between wells. Plates were secured to a 200 RPM shaker at 28 °C and incubated for 10 hours. Regular measurements of the $OD_{600}$ were recorded with an EnVision 2105 multimode plate reader (Perkin Elmer). Each replicate was blank corrected against sterile media wells with the same concentration of AQ. The growth curves for each concentration of AQ were normalized to the DMSO growth curve with the following equation:

$$\% \text{ Normalized } OD_{600} = ({}_{[AQ]}OD_{600} - {}_{DMSO}\text{Minimum}_{OD600})/({}_{DMSO}\text{Maximum}_{OD600} - {}_{DMSO}\text{Minimum}_{OD600}) * 100.$$

The response to different concentrations of AQs was then defined as the percentage $OD_{600}$ absorbance of the vehicle control. Data plots were prepared in GraphPad Prism version 8.0.0 for Windows. Lines connecting the points were added for visual clarity by Spline fitting and do not represent a line of best fit.

##### *M. rhizosphaerae* response to HHQ and Br-HHQ

AQ response growth curves for *M. rhizosphaerae* were performed as for *V. natriegens*, however the $OD_{600}$ values were monitored for 15 hours.

##### Titration of VHPO substrates in *O. marinus* culture

50 mL MMBP media was inoculated from an *O. marinus* starter culture and grown to an $OD_{600}$ of 0.5. 50 mL preparations of MMBP media with the appropriate concentrations of added $Na_3VO_4$ and KBr were inoculated with *O. marinus* to an $OD_{600}$ of 0.05. Plates with AQ concentration gradients for each media condition were then prepared, incubated, and monitored as above. [AQ] growth curves were normalized to the DMSO control growth curve of the same media condition. Data plots were prepared in GraphPad Prism version 8.0.0 for Windows. Lines connecting the points were added for visual clarity by Spline fitting and do not represent a specific fit.

##### omVHPO qPCR

50 mL MMBP media was inoculated from an *O. marinus* starter culture and grown to an $OD_{600}$ of 0.5. This starter culture was diluted into fresh MMBP media supplemented with 10 nM $Na_3VO_4$ and 2 mM KBr to an $OD_{600}$ of 0.02 and aliquoted into 40 mL cultures. AQs were added to intended final concentrations by the addition of 40 µL AQ from 100x stock solutions in DMSO. The vehicle control was prepared by adding 40 µL DMSO to the appropriate wells. Flasks were incubated at 28 °C and 200 RPM for 7 days. At regular intervals, 1 mL of culture was removed for total RNA extraction and 0.5 mL was removed for $OD_{600}$ monitoring. RNA was extracted in an RNAse free environment following the manufacturer instructions with the PureLink RNA Mini Kit

(Invitrogen). RNA quality and concentration was checked by NanoDrop. All RNA extracts were diluted to 10 ng/µL in RNAse free water. cDNA conversion was performed immediately after RNA extraction following the manufacturer specifications using a High-Capacity cDNA Reverse Transcription Kit (Thermo). 10 µL of 10 ng/µL RNA was used as the template for the reaction and reaction was performed in a thermocycler. No Reverse Transcriptase (NRT) controls were simultaneously prepared by omitting the reverse transcriptase. 25 µL qPCR reactions were prepared with a QuantiTect SYBR Green PCR kit (Qiagen) according to manufacturer instructions and aliquoted into 0.2 mL Non-skirted Low Profile 96-well PCR Plates (Thermo) and sealed with Microseal 'B' seals (Bio-rad). Primers for qPCR are listed in Table S5. Each primer pair was prepared with NCBI Primer-BLAST and checked against the genome of *O. marinus* for gene specificity. No template controls were prepared for each primer pair and primer pairs were confirmed to have comparable replication efficiencies. Two technical replicates were prepared for each biological replicate. PCR was run on a CFX Connect Real Time PCR Detection System (Bio-Rad Laboratories) and controlled with CFX maestro 2.0 (Bio-Rad Laboratories) with the following thermocycling method: initial denaturation at 95 °C (15 minutes), 40 cycles of melt at 94 °C (15 seconds) and anneal at 53 °C (30 seconds), elongate at 72 °C (30 seconds), and record fluorescence. After 40 cycles had completed, a melt curve of 60 °C to 95 °C with 0.2 °C increments every 15 seconds would be run for each plate. Fold change of gene expression data was refined using CFX Maestro 2.0. All VHPO amplification was referenced to the amplification of both 16s rRNA (NCBI: NR_112787.1) and DNA gyrase, gyrA (NCBI: WP_076516024.1). The fold VHPO expression from each condition was normalized to the first collected timepoint. Data was prepared as a box-and-whisker plot in GraphPad Prism version 8.0.0 for Windows.

**Mass Spectrometry Imaging Experiments**

Sterilized 10 mL MMBP + 1.0% agar was poured into 100 mm petri dishes. Plates were left slightly open for 30 minutes in a laminar flow hood and then closed and left in laminar flow hood for 16 hours before being used immediately. $Na_3VO_4$ was not added to solid media due to it causing severe restriction of *P.* aeruginosa PA14 growth. O. *marinus* cultures were grown to an $OD_{600}$ of 0.5 and then supplemented with 10 nM $Na_3VO_4$ and 2 mM KBr. 3 µL of culture was gently pipetted onto MMBP agar and allowed to dry. Several spots were placed on the plates at regular intervals. For plates intended for co-culture, spots were restricted to one half of the plate. Plates were then placed at 28 °C for 5 days to allow for adequate density of *O. marinus* to grow. *P. aeruginosa* PA14 wild type, *ΔpqsA-C*, and *ΔpqsH* were inoculated from glycerol stocks into 5 mL LB media and grown at 30 °C and 200 RPM for 20 hours. *P. aeruginosa* PA14 cultures were then diluted to an $OD_{600}$ of 0.5 and gently spotted as 3 µL aliquots next to or touching established *O. marinus* colonies. In the case of Br-HHQ spiking of *O. marinus* colonies, 3 µL of 50 µM Br-HHQ was directly spotted on top of the *O. marinus* colony. All plates were allowed to dry and then returned to 28 °C for 48 hours. Following 48 hours of *P. aeruginosa* PA14 growth, optical images of the plates were taken. Colonies were excised with a razor blade and transferred to an MSP 96-target ground steel plate (Bruker Daltonics) and optical images were taken. A 1:1 mixture of recrystallized CHCA and DHB was used as MALDI matrix. The target plate was evenly coated with matrix using a 53 µm stainless steel sieve (Hogentogler, Inc) and then placed in a 37 °C oven. After 30 minutes, plates were removed from oven and matrix was re-applied. Plates were replaced in oven and left to dry until agar was fully desiccated (about 4 hours). After agar had fully desiccated, excess matrix was removed from the plate with a stream of air. Plates were then placed in a vacuum chamber with Drierite overnight. Optical images of the matrix-coated desiccated colonies were taken prior to loading the target plate into the MALDI-QqTOF MS (Bruker timsTOF fleX) The plates were analyzed using timsControl v.4.1.12 and FlexImaging v.7.2 software. The detector gain and laser power were set to 3.0x and 30% respectively. The detection range was set from 100-600 Da with ion suppression set at 50 Da in positive reflectron mode. The laser set to 500 µm raster size and 200 shots per raster point. SCiLS software (Bruker, v 2023b) was used to analyze and visualize collected data. Ion bin was set to 1,000,000 during the import. Data was normalized with root-mean-square (RMS), an error of ±0.2 Da, and weak denoising was applied. Ions for HHQ, HQNO/PQS, and Br-HHQ were checked. Data was visualized using magma heatmap coloration. All mass annotations were targeted. MALDI-MSI data was reported following SMART protocols.[20] Exact masses used to calculate mass confidence intervals for each compound follow: 244.1696 (HHQ), 260.1645 (PQS/HQNO), 322.080 ($^{79}$Br-HHQ), 324.0781 ($^{81}$Br-HHQ).A similar protocol was followed for *V. natriegens* and *M. rhizosphaerae* co-cultures with the following alterations. 10 mL LB media + 1.5% agar was used as media but prepared the same otherwise. The two strains and *P. aeruginosa* PA14 were inoculated at the same time and grown over the course of 48 hours.

**P. aeruginosa PA14 swarming motility assay**

Swarming motility assay conditions were based on previous work done with *P. aeruginosa* PA14 and AQs.[21,22] LB media with 0.5% agar was used for all swarming plates. The swarming motility of *P. aeruginosa* PA14 was tested in the presence of DMSO, ciprofloxacin, PQS, HHQ, and Br-HHQ. 100x stock solutions of ciprofloxacin, PQS, HHQ, and Br-HHQ were prepared at the appropriate concentrations. Compounds were added to molten LB agar immediately before pouring plates. All plates were prepared by pouring 20 mL of LB agar + compound into 100 mm petri dishes. Plates were allowed to set for 30 minutes and were then flipped and left in laminar flow hood for 15 hours. 10 µM ciprofloxacin, 10 µM PQS, 10 µM HHQ, 10 µM Br-HHQ, 50 µM HHQ, 50 µM Br-HHQ, 100 µM HHQ, and 100 µM Br-HHQ conditions were prepared. *P. aeruginosa* PA14 WT was inoculated from glycerol stock into 5 mL LB media and incubated for 20 hours at 37 °C. *P. aeruginosa* PA14 culture was adjusted to an $OD_{600}$ of 1.0 and each plate was inoculated in the center with 4 µL of culture. Inoculant was allowed to dry and then incubated overnight at 30 °C. Extent of swarming was recorded at 16 hours and 24 hours by optical imaging. Swarm areas were measured using Image J.[23]

#### Supplementary Figures

##### Figure S1: HZ11-VHPO Sequence Similarity Networks

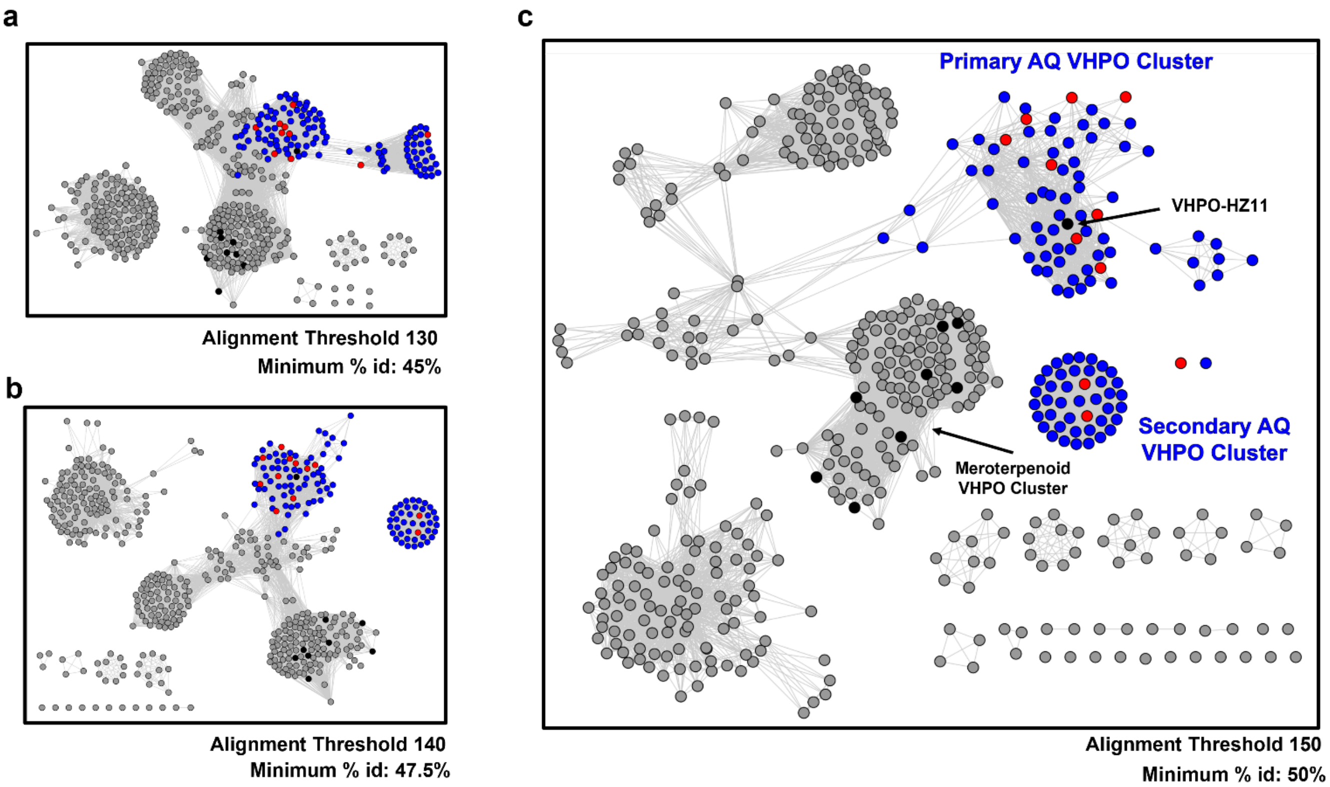


Sequence Similarity Network (SSN) of the top 500 most similar protein sequences to HZ11-VHPO. Each circle (node) represents a unique protein sequence and the lines that connect them (edge) represent a minimum sequence identity determined by the alignment threshold. Previously characterized selective VHPOs (black circles), putative meroterpenoid VHPO cluster (orange circles), putative AQ-VHPOs from the main AQ cluster identified in this study (blue circles), putative AQ-VHPOs from the secondary AQ cluster identified in this study (blue circles), VHPOs with AQ-bromination activity confirmed in this study (red circles), and uncharacterized VHPOs (grey circles) are indicated. The defined formation of the meroterpenoid VHPO cluster was used to aid in defining minimal alignment thresholds to refine the putative AQ-VHPO cluster. (a) The alignment threshold of 130 roughly equates to a minimum sequence identity of 45. (b) The alignment threshold of 140 roughly equates to a minimum sequence identity of 47.5%. (c) The alignment threshold of 150 roughly equates to a minimum sequence identity of 50%.

**Figure S2: Taxonomic and environmental sources of putative AQ-VHPO containing organisms**

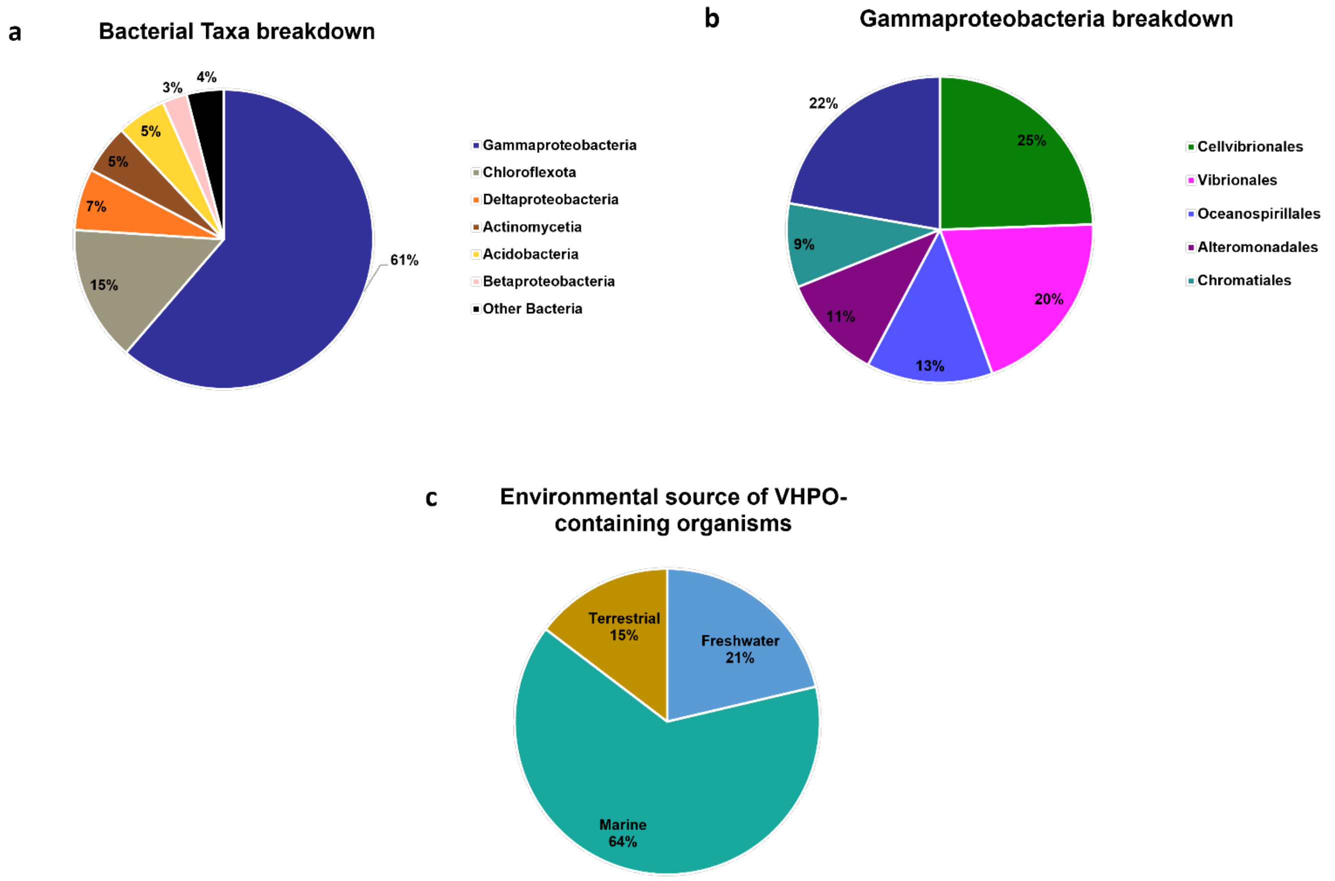


(a) Visual representation the distribution across bacterial Orders of all identified putative AQ-VHPO containing source organisms listed in Table S1. (b) Visual representation the distribution across varied gammaproteobacterial classes that contain identified putative AQ-VHPOs. (c) Visual representation of the environmental sources of putative AQ-VHPO containing organism. Environmental sources were defined by the bacteria's original source of isolation, accessed through NCBI. Marine environments include open ocean, marine sediment, marine organism associated symbiotes, and salt marshes. Terrestrial environments include soil and terrestrial rhizospheres. Freshwater environments include ponds, lakes, wastewater, and groundwater.

**Figure S3: General HHQ biosynthetic pathway**

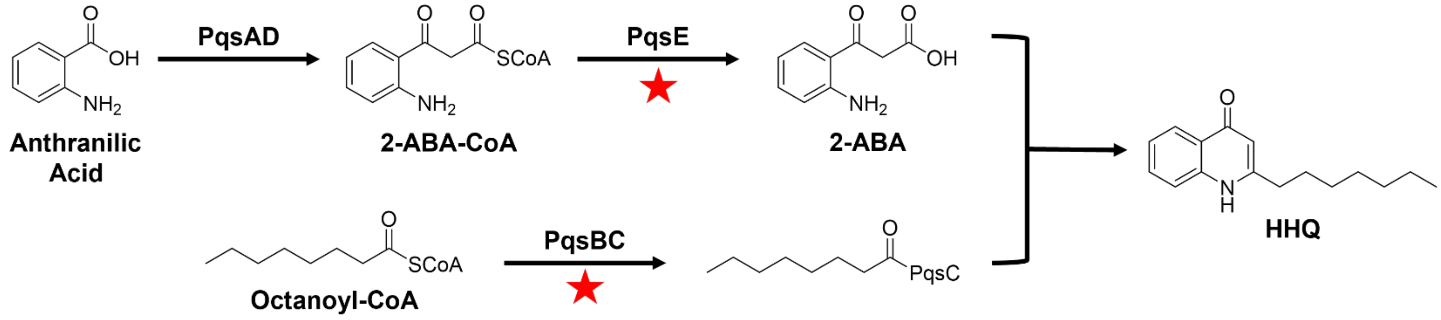


General biosynthetic pathway of HHQ, as characterized in *Pseudomonas aeruginosa*.[24,25] Required genes specific to the pathway are highlighted with red stars.

**Figure S4: HZ11-VHPO crude lysate HHQ bromination activity**

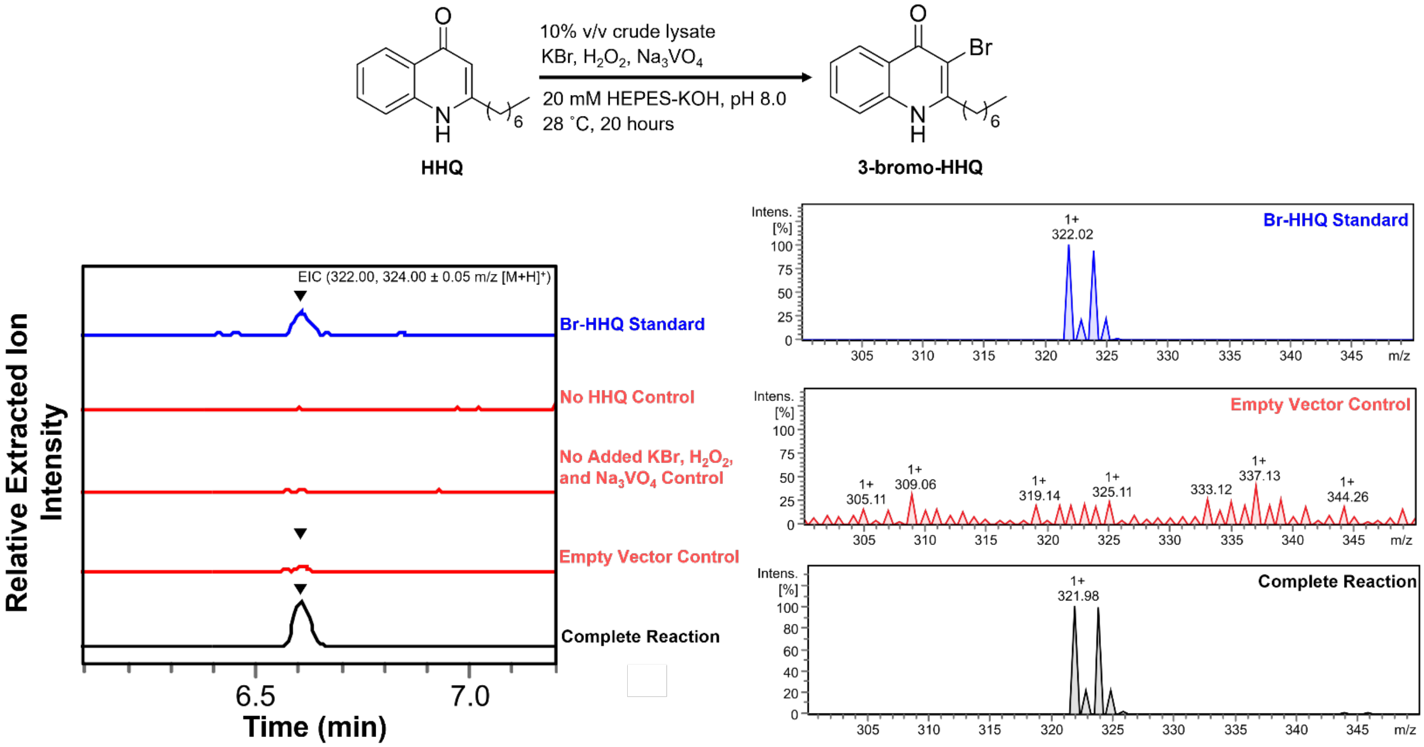


Enzyme:Empty Vector peak area ratio: 955

Extracted ion chromatogram (EIC) traces of HHQ bromination activity by the crude lysate of heterologously expressed HZ11-VHPO, sourced from *Microbulbifer* sp. HZ11. Extracted ions are given in the upper right of the trances. Generalized reaction conditions, AQ substrate, and product are given above the traces. Synthetic standard traces (blue), negative control traces (red), and experimental traces (black) are indicated. Enzyme:Empty Vector ratio represents the internal standard normalized size of the Br-HHQ peak area in the experimental condition compared to Empty Vector control. The MS for the extracted ion’s peak at its apex of intensity is given for each VHPO’s reaction

**Figure S5: esVHPO crude lysate HHQ bromination activity**

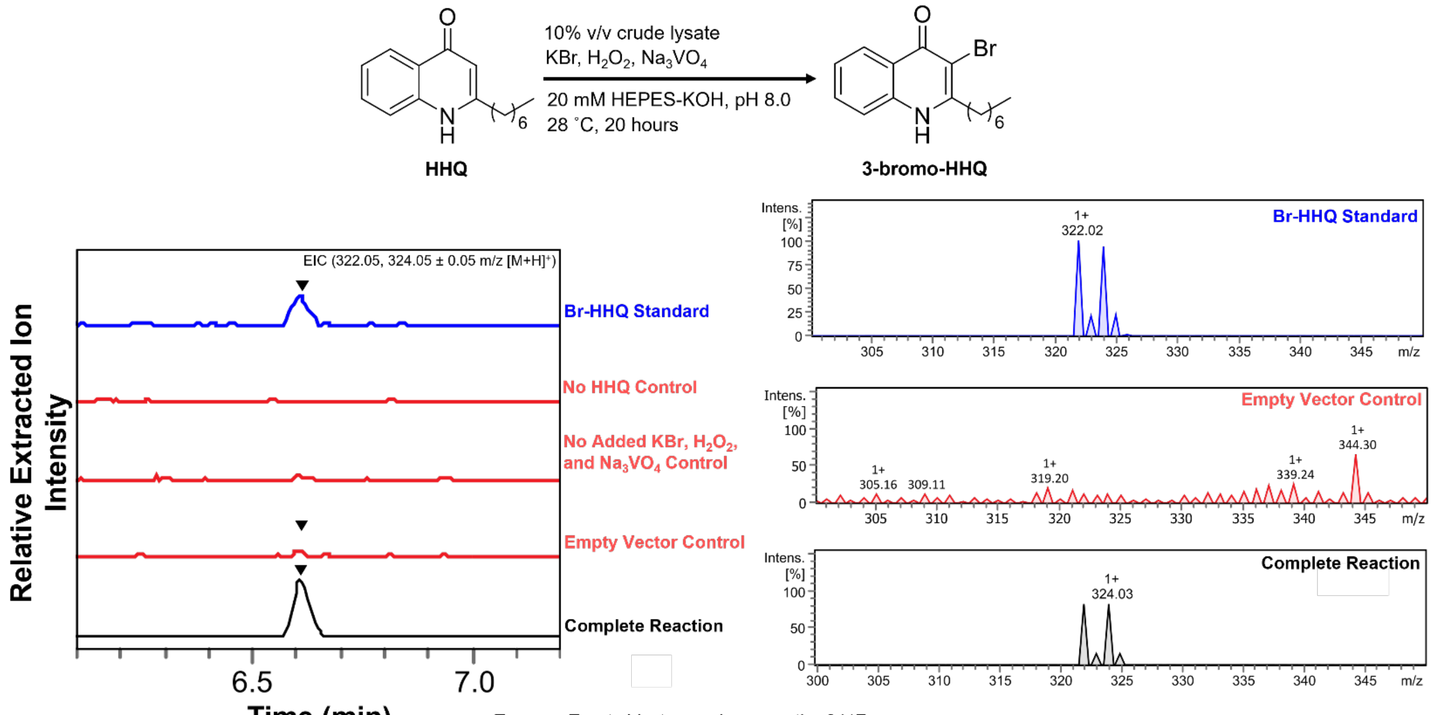


Enzyme:Empty Vector peak area ratio: 2117

Extracted ion chromatogram (EIC) traces of HHQ bromination activity by the crude lysate of heterologously expressed esVHPO, sourced from *Enhygromyxa salina*. All other components are as SI Fig. 3.

**Figure S6: lpVHPO crude lysate HHQ bromination activity**

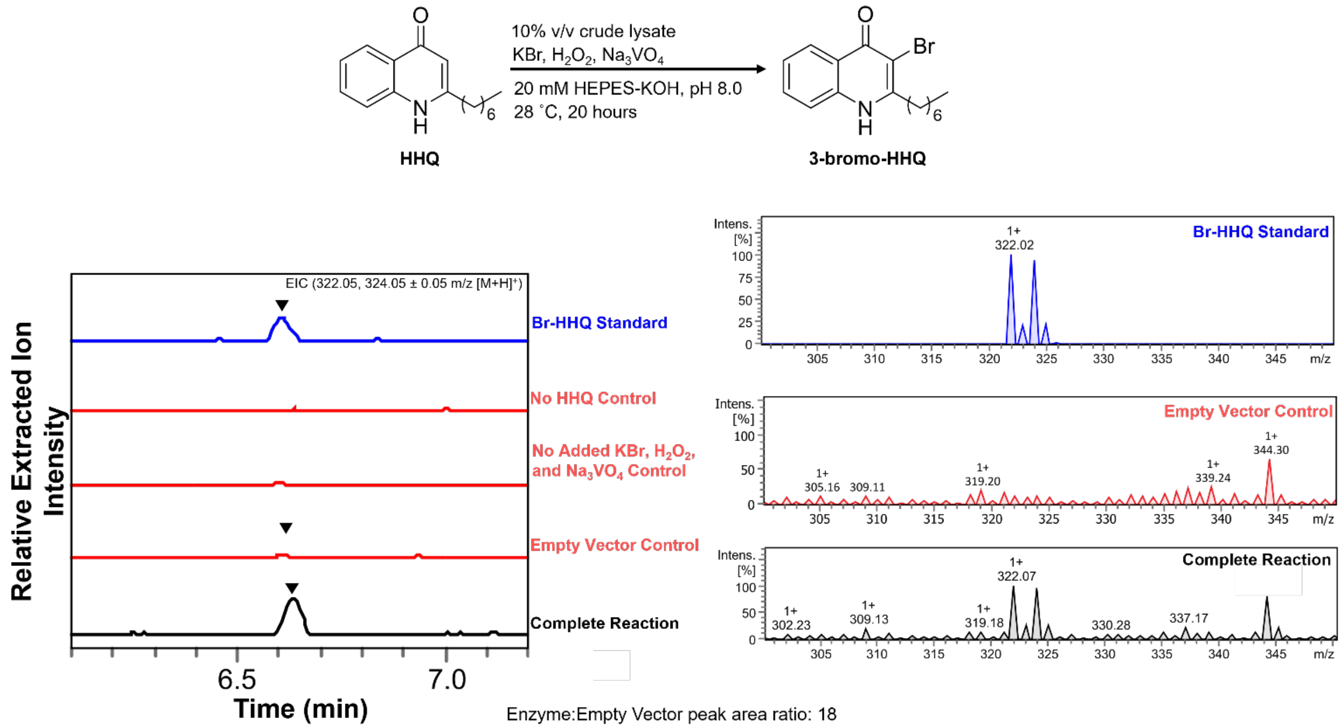


Extracted ion chromatogram (EIC) traces of HHQ bromination activity by the crude lysate of heterologously expressed lpVHPO, sourced from *Luteitalea pratensis*. All other components are as SI Fig. 3.

#### Figure S7: moVHPO crude lysate HHQ bromination activity

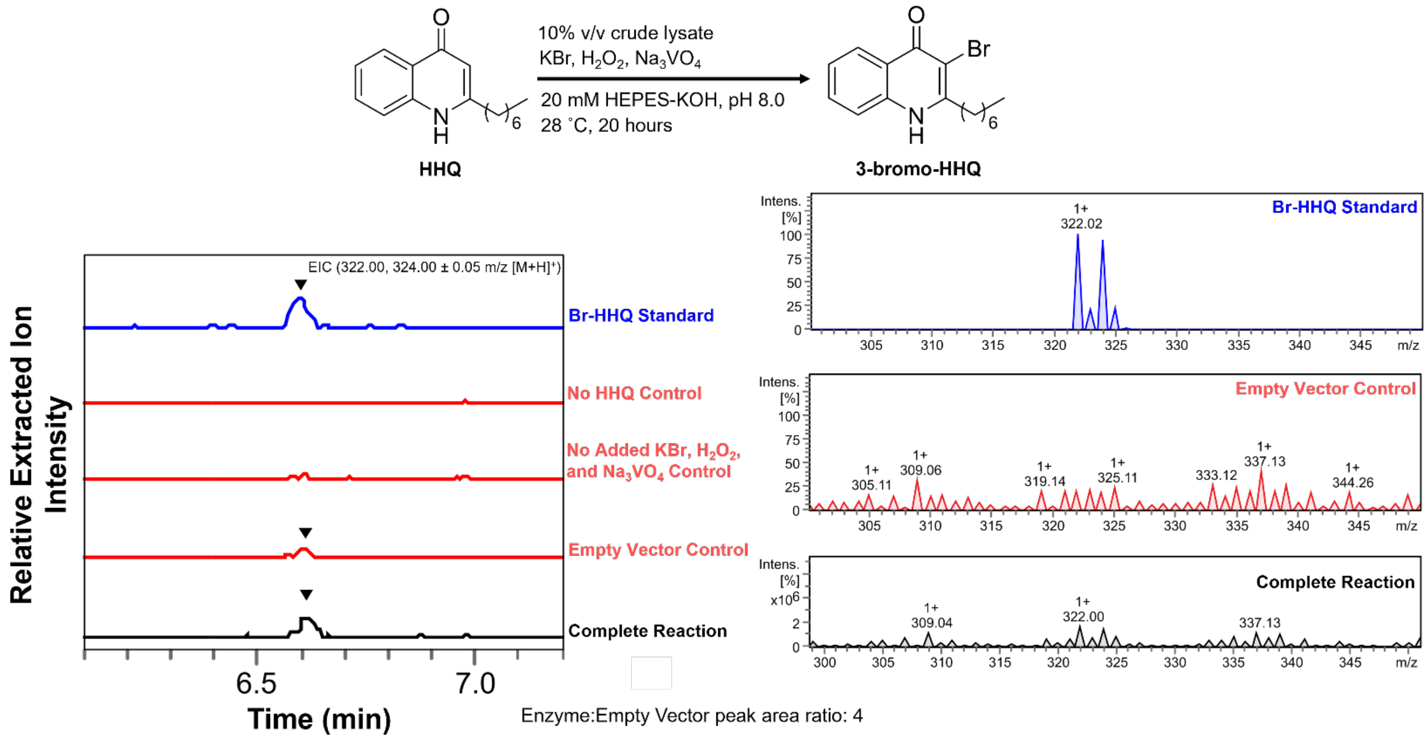


Extracted ion chromatogram (EIC) traces of HHQ bromination activity by the crude lysate of heterologously expressed moVHPO, sourced from *Methylotetracoccus oryzae*. All other components are as SI Fig. 3.

**Figure S8: mrVHPO crude lysate HHQ bromination activity**

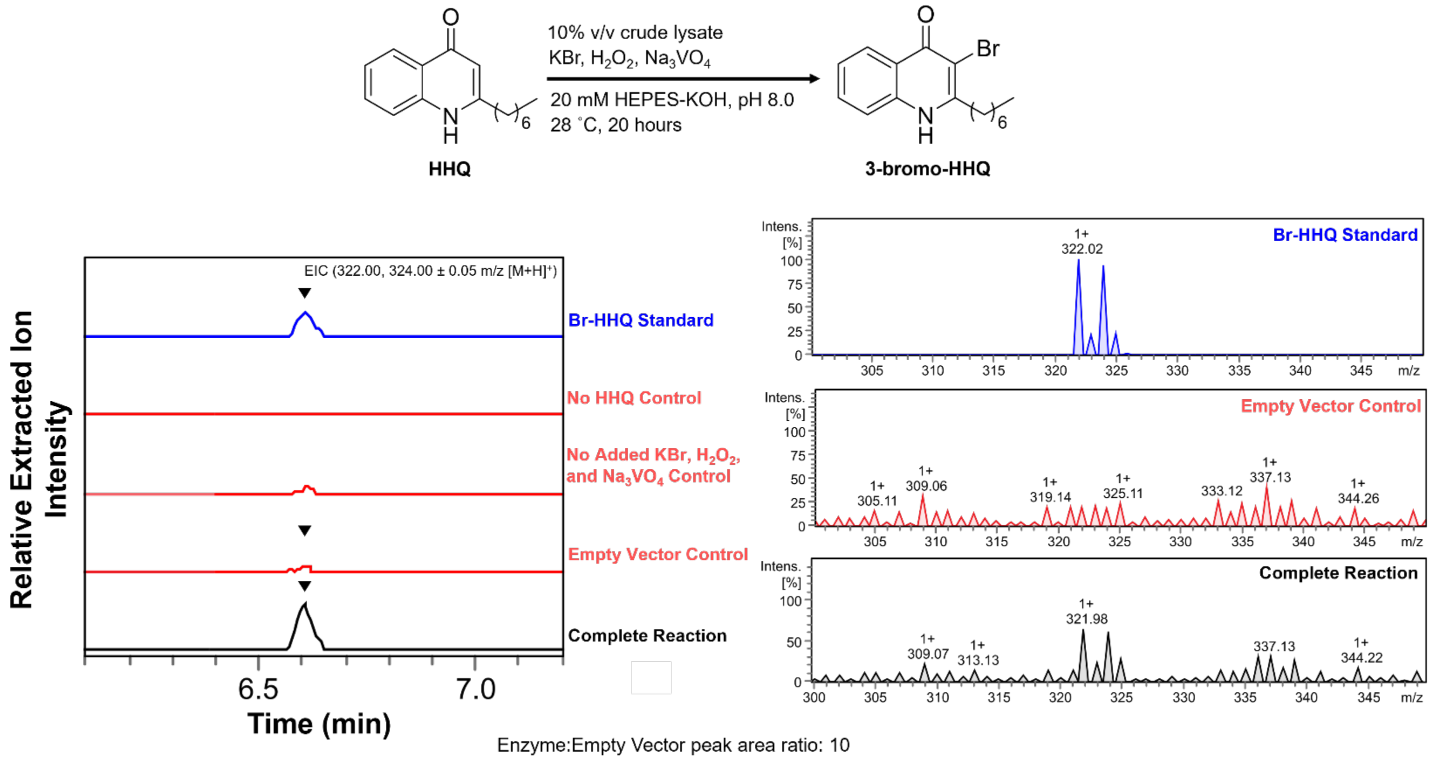


Extracted ion chromatogram (EIC) traces of HHQ bromination activity by the crude lysate of heterologously expressed mrVHPO, sourced from *Microbulbifer rhizosphaerae*. All other components are as SI Fig. 3.

**Figure S9: A4B17-VHPO crude lysate HHQ bromination activity**

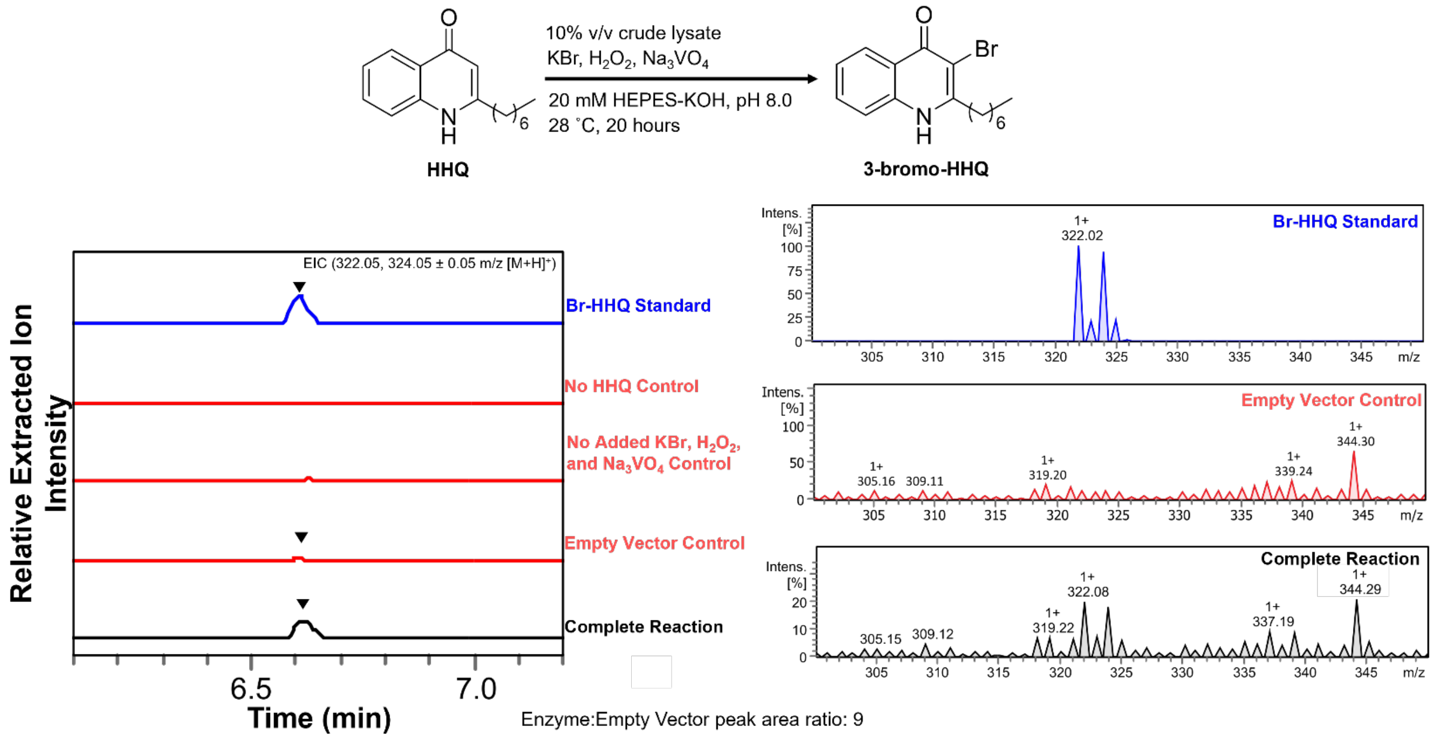


Extracted ion chromatogram (EIC) traces of HHQ bromination activity by the crude lysate of heterologously expressed A4B17-VHPO, sourced from *Microbulbifer* sp. A4B17. All other components are as SI Fig. 3.

**Figure S10: omVHPO crude lysate HHQ bromination activity**

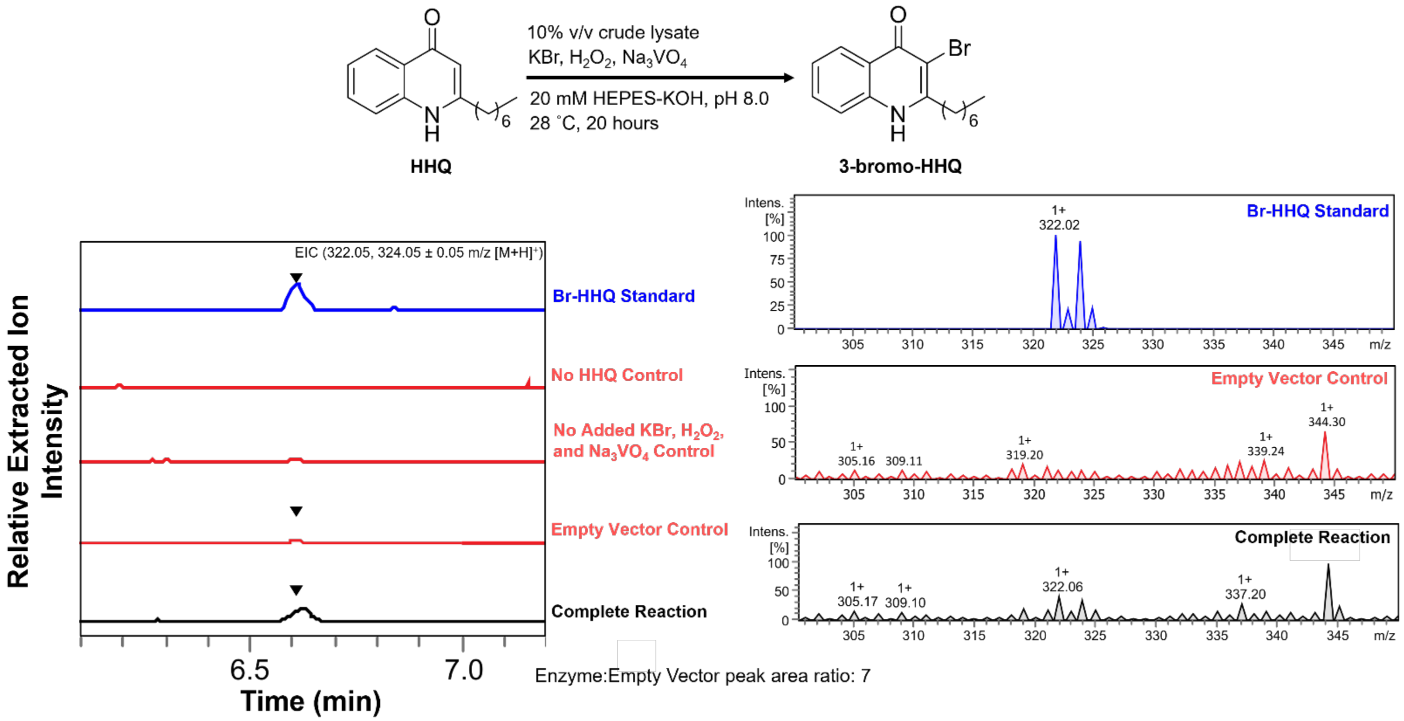


Extracted ion chromatogram (EIC) traces of HHQ bromination activity by the crude lysate of heterologously expressed omVHPO, sourced from *Oleibacter marinus*. All other components are as SI Fig. 3.

**Figure S11: plVHPO crude lysate HHQ bromination activity**

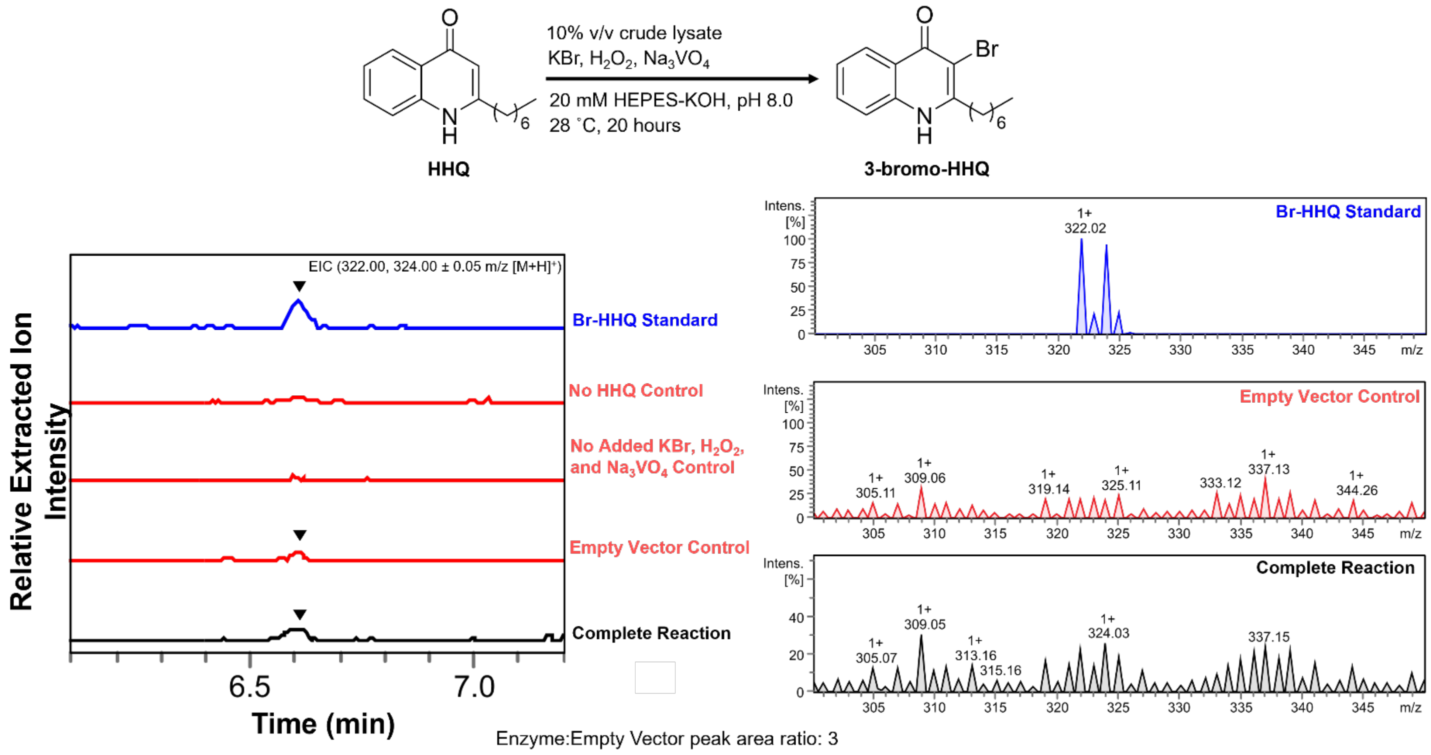


Extracted ion chromatogram (EIC) traces of HHQ bromination activity by the crude lysate of heterologously expressed plVHPO, sourced from *Pseudoalteromonas luteovioleacea* H33-S. All other components are as SI Fig. 3.

#### Figure S12: SA03-VHPO crude lysate HHQ bromination activity

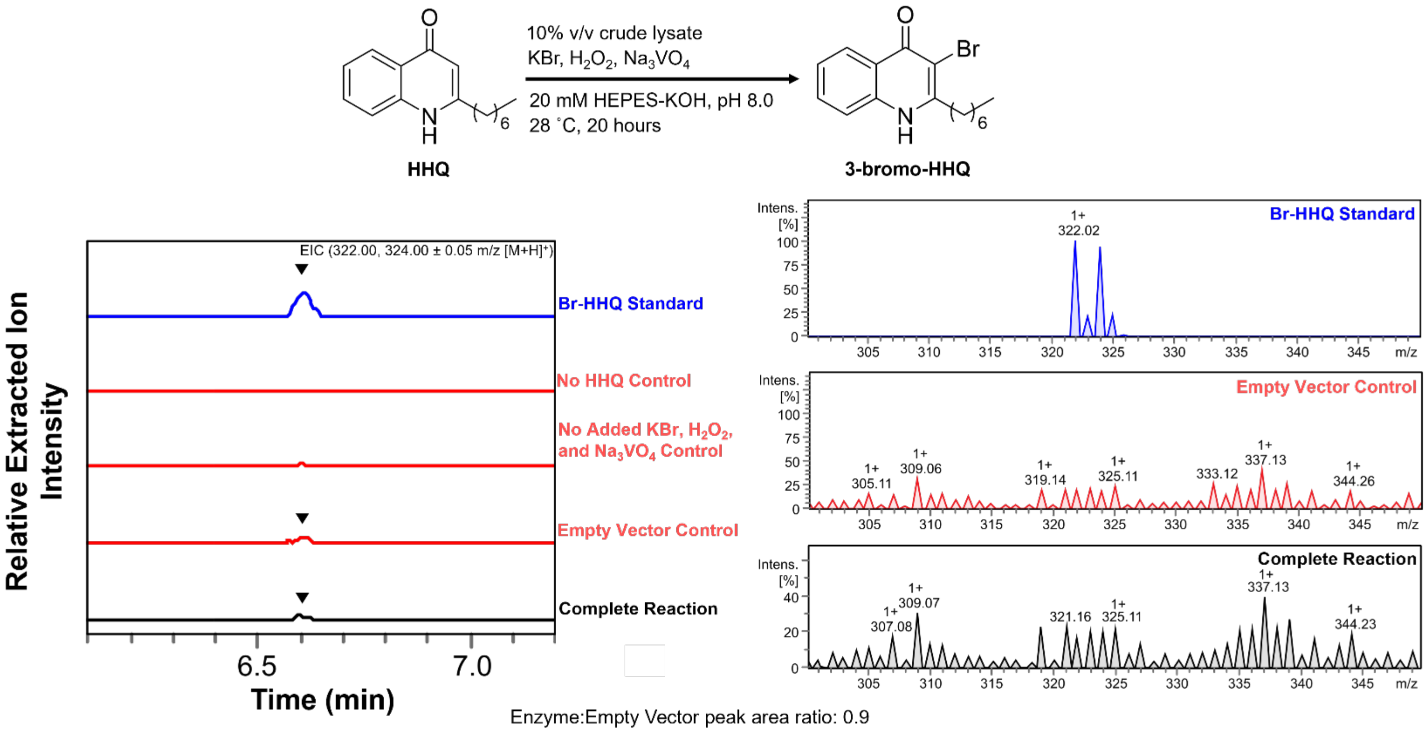


Extracted ion chromatogram (EIC) traces of HHQ bromination activity by the crude lysate of heterologously expressed SA03-VHPO, sourced from *Pseudoalteromonas* HM-SA03. All other components are as SI Fig. 3.

#### Figure S13: raVHPO crude lysate HHQ bromination activity

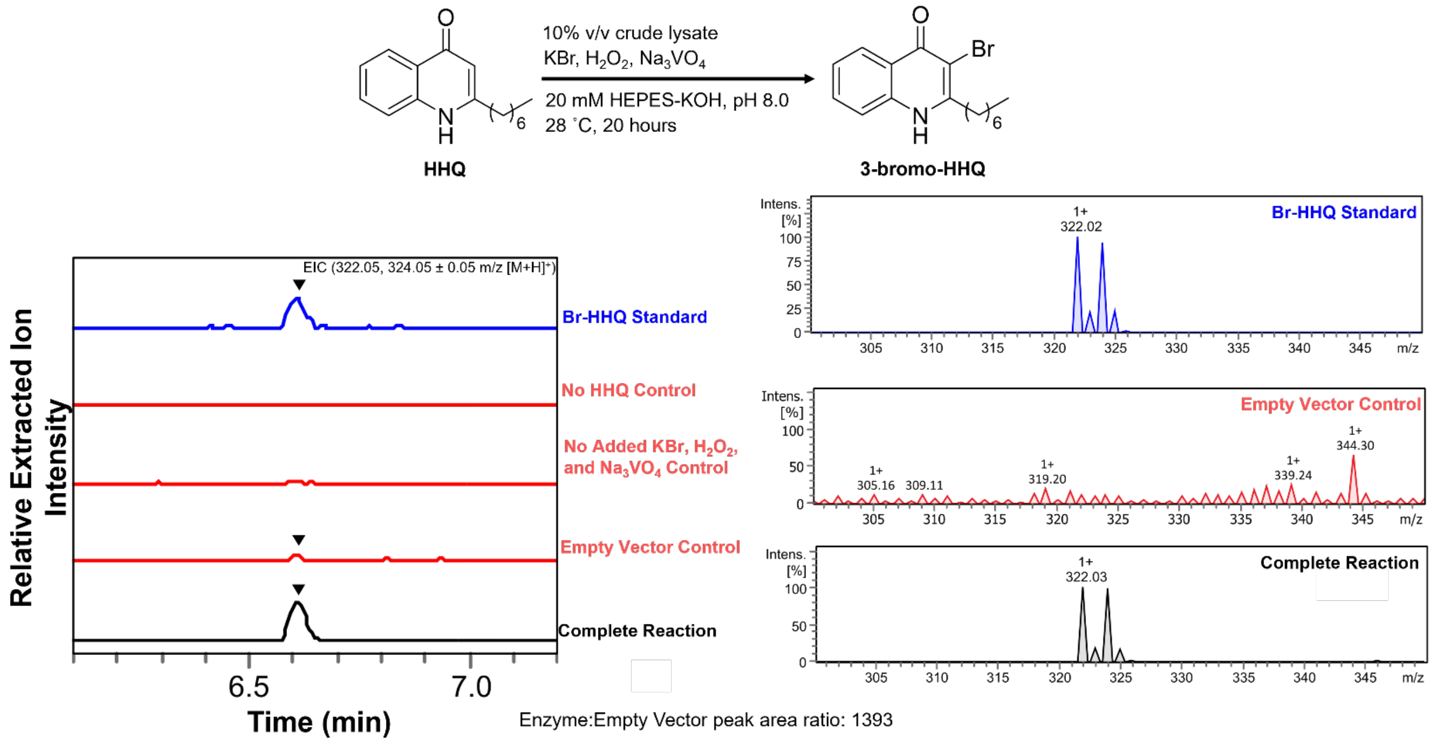


Extracted ion chromatogram (EIC) traces of HHQ bromination activity by the crude lysate of heterologously expressed raVHPO, sourced from *Rubrivivax albus*. All other components are as SI Fig. 3.

**Figure S14: thVHPO crude lysate HHQ bromination activity**

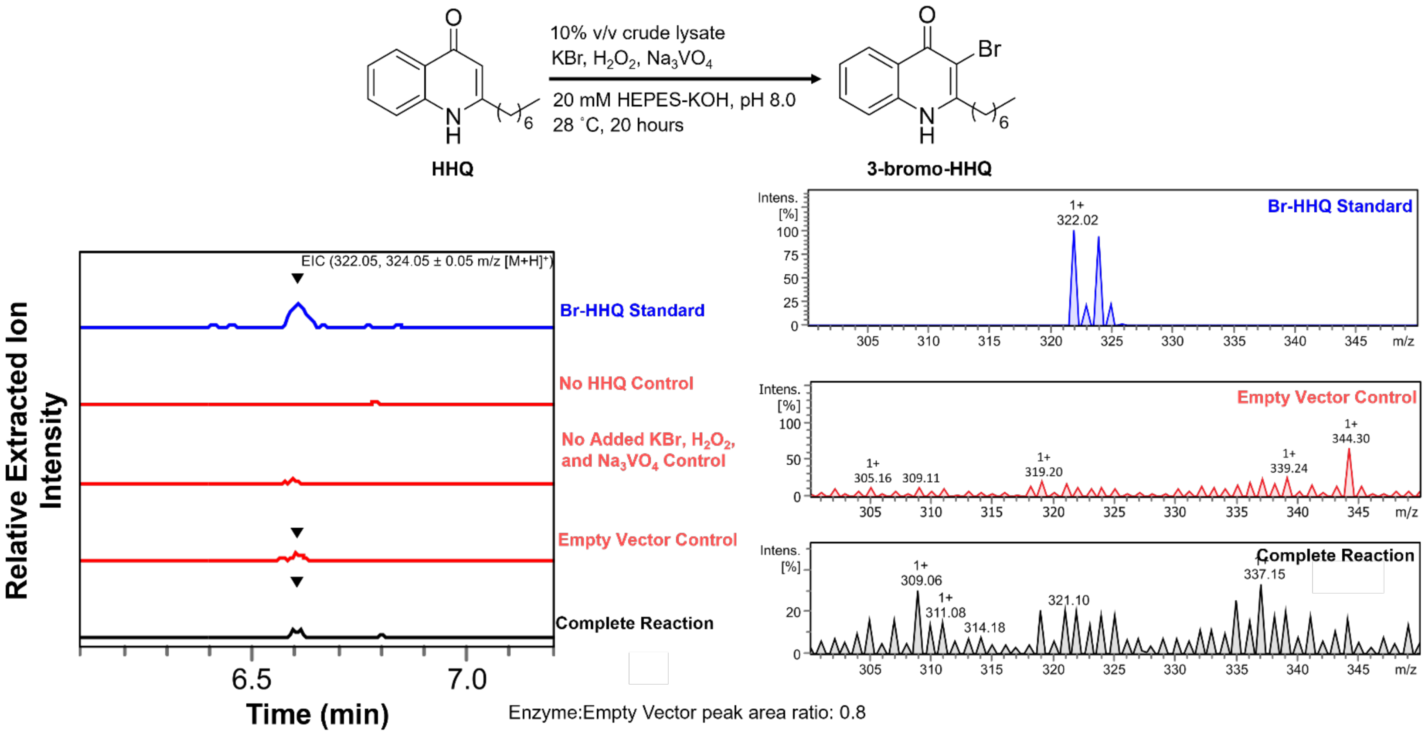


Extracted ion chromatogram (EIC) traces of HHQ bromination activity by the crude lysate of heterologously expressed thVHPO, sourced from *Tamaricihabitans halophyticus*. All other components are as SI Fig. 3.

**Figure S15: trVHPO crude lysate HHQ bromination activity**

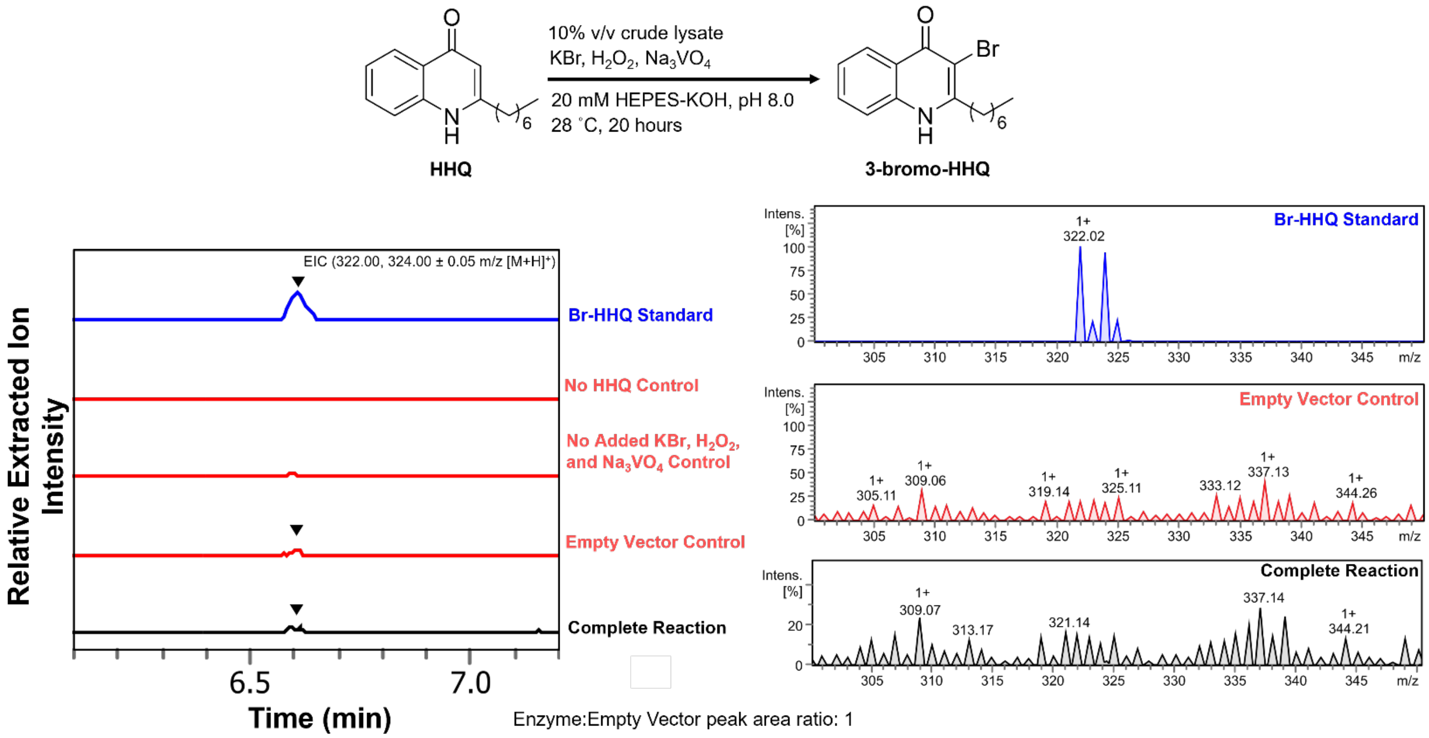


Extracted ion chromatogram (EIC) traces of HHQ bromination activity by the crude lysate of heterologously expressed trVHPO, sourced from *Thiocapsa roseopersicina*. All other components are as SI Fig. 3.

**Figure S16: vbVHPO crude lysate HHQ bromination activity**

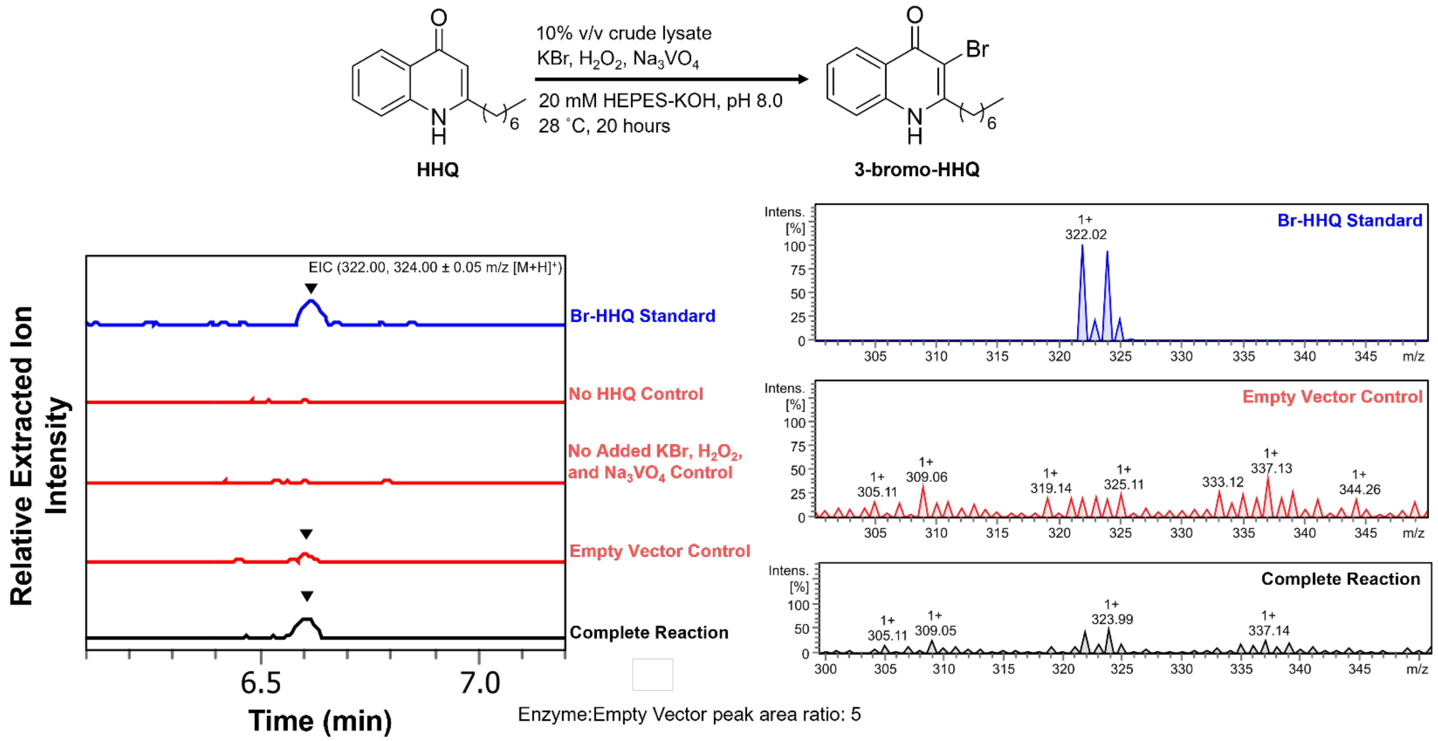


Extracted ion chromatogram (EIC) traces of HHQ bromination activity by the crude lysate of heterologously expressed vbVHPO, sourced from *Vibrio breoganii* 1C10. All other components are as SI Fig. 3.

**Figure S17: vnVHPO crude lysate HHQ bromination activity**

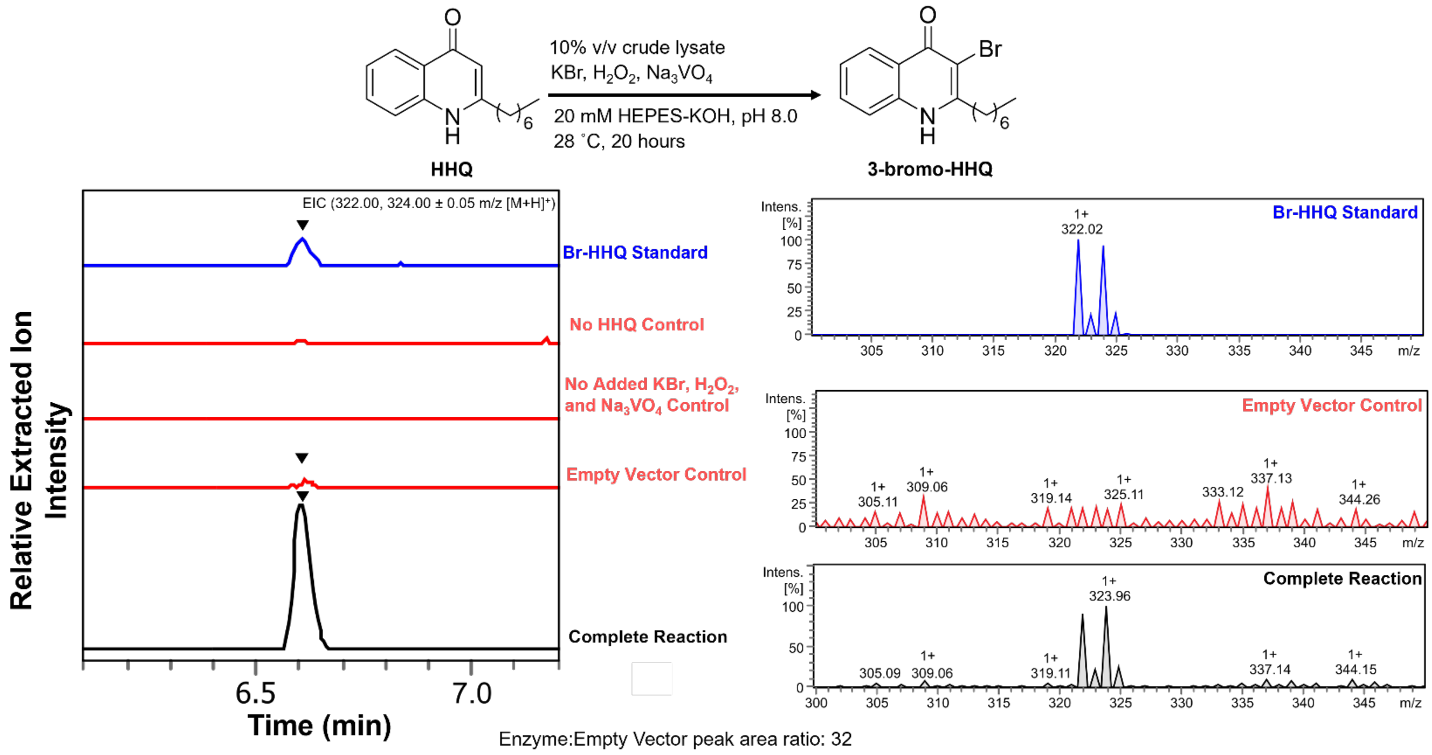


Extracted ion chromatogram (EIC) traces of HHQ bromination activity by the crude lysate of heterologously expressed vnVHPO, sourced from *Vibrio natriegens* CCUG 16374. All other components are as SI Fig. 3.

**Figure S18: Environmental distribution of confirmed AQ-brominating VHPOs**

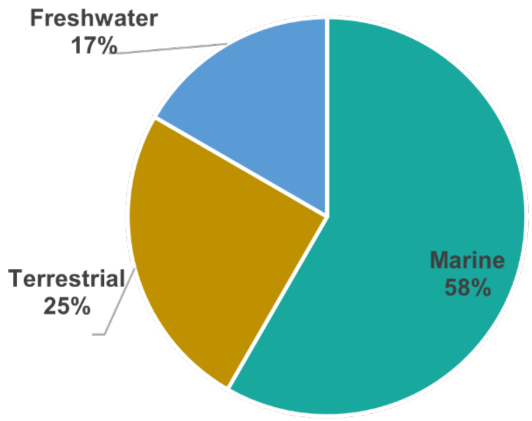


Environmental distribution of source species containing confirmed active AQ-VHPOs. The proportion of marine organisms are shown in teal, terrestrial organisms are in brown, and freshwater organisms are in light blue.

**Figure S19: Sequence Alignment of AQ-VHPOs**

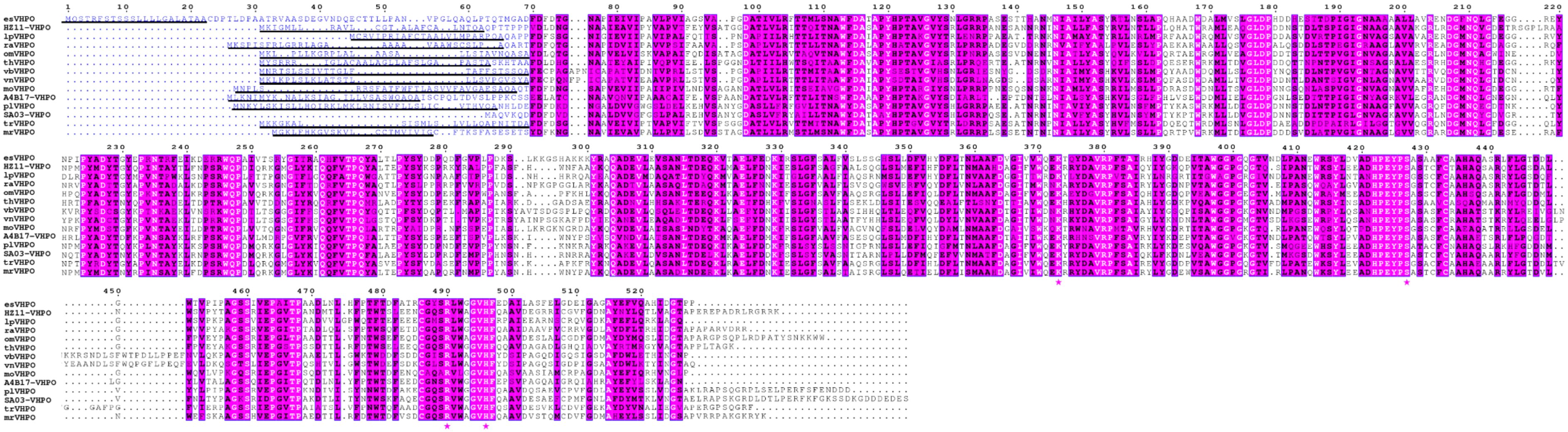

Sequence alignment of VHPOs that were tested for AQ bromination activity in crude lysate. Initial alignment was prepared with ClustalOmega[13] and was prepared in ESPript[14]. Residues blocked in purple and in white text are strictly conserved, residues blocked in purple and in bold black text are functionally conserved and at least 70% identical. The variable N-terminal region is highlighted in blue text and the SignalP 6.0[26] predicted signal peptide sequence is underlined.

**Figure S20: Conservation of secretion signals across all AQ-VHPOs**

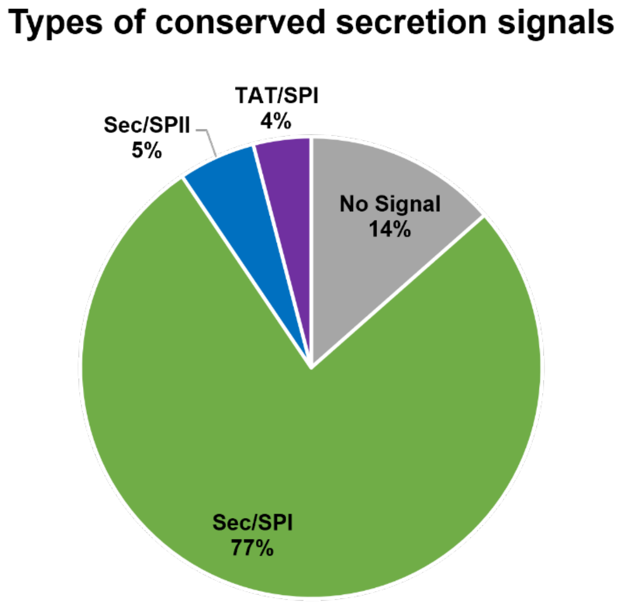


Visual representation of the conservation of secretion signals across putative AQ-VHPOs. All signal types and predictions were identified using SignalP 6.0.[26] Attribution of each signal and predicted signal cleavage sites to specific putative AQ-VHPOs are given in Table S6.

**Figure S21: Phylogenetic distinction between the presence and absence of signal peptides in characterized VHPOs**

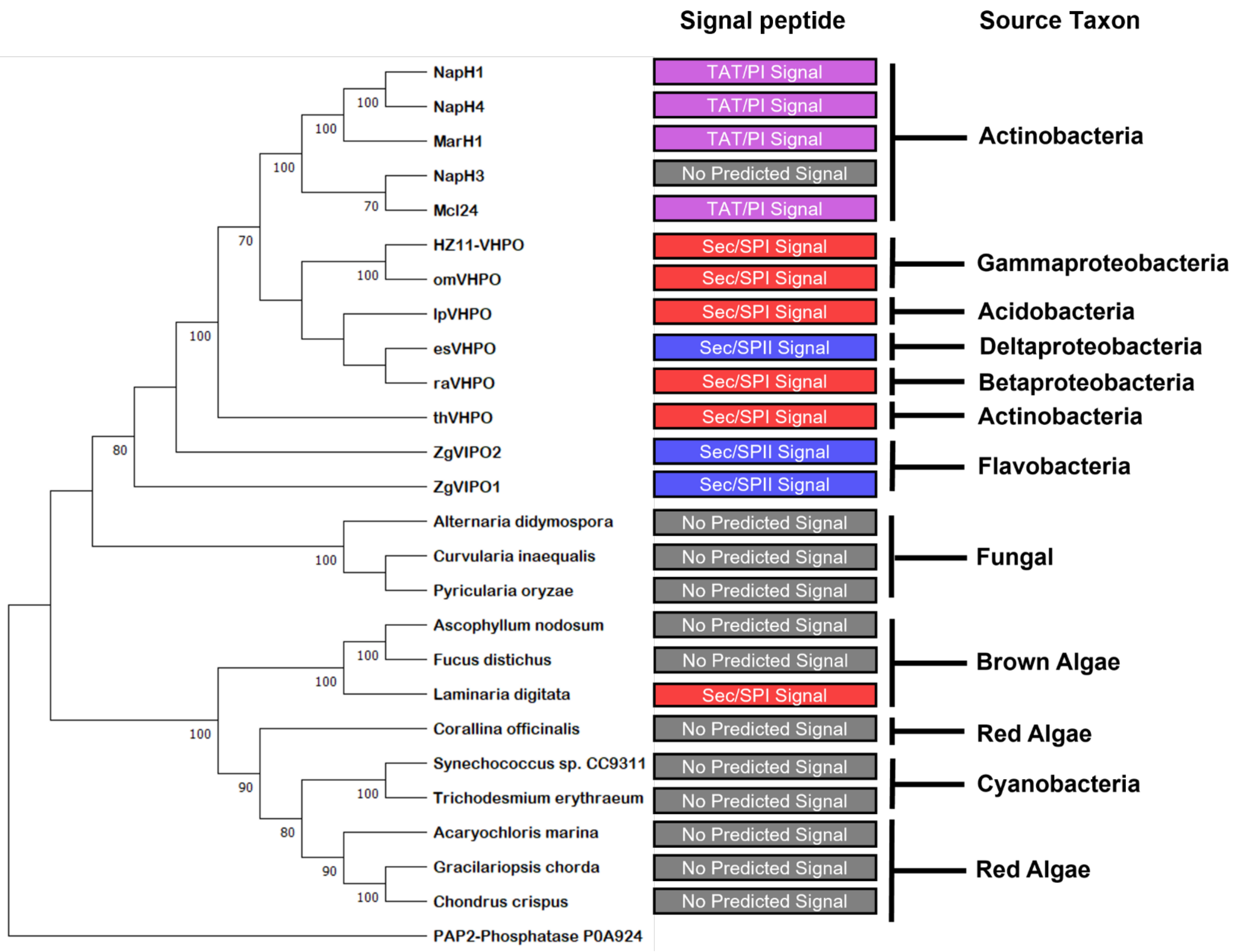


Representative phylogenetic tree for characterized VHPOs from across all kingdoms and chemistry types. All eukaryotic, flavobacterial, and cyanobacterial VHPOs listed here have been characterized as non-selective. Signal peptides were predicted with SignalP 6.0[26] using the eukaryotic, Gram-negative, and Gram-positive settings as appropriate.[26] A PAP2-Phosphatase from *E. coli* K12 was added to the tree as the root. Accession numbers for each gene product are listed in Table S7.

**Figure S22: SDS-PAGE of purified AQ-VHPOs**

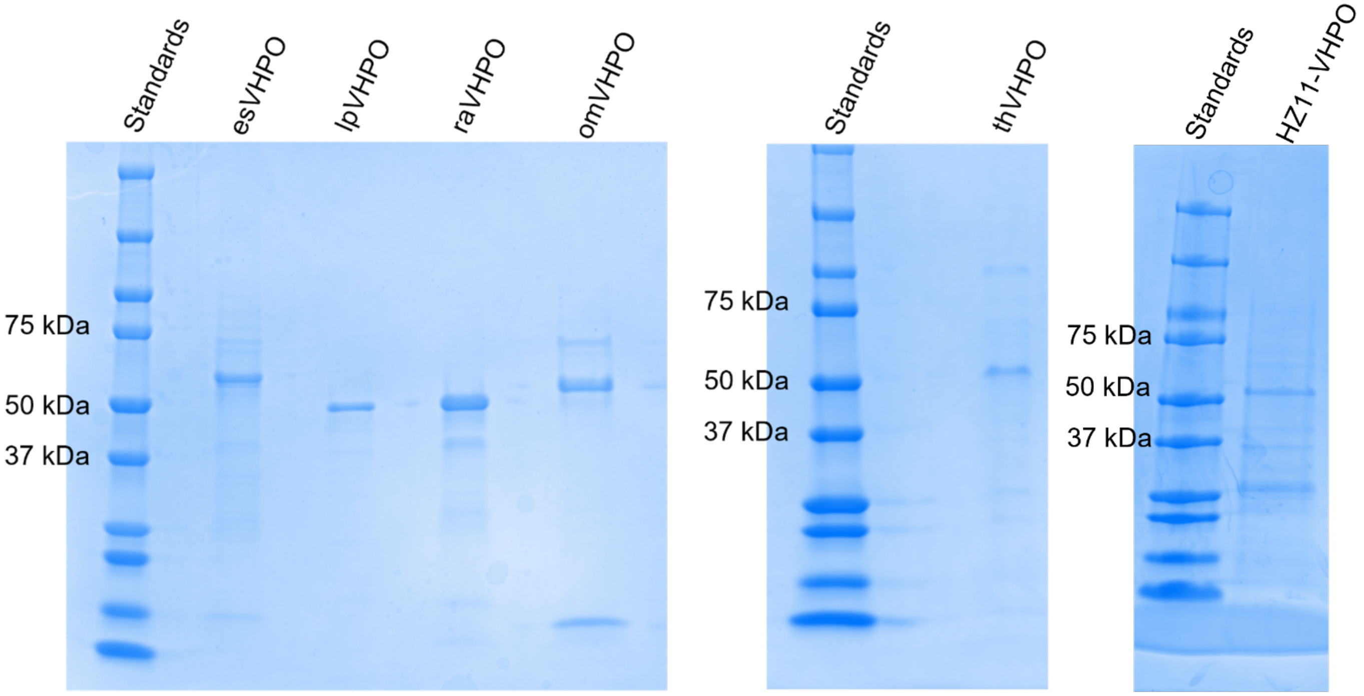


SDS-PAGE of homogenously purified AQ-VHPOs. Both gels were stained with Coomassie blue and Promega Precision Plus Protein Dual Color Standards was used as reference for both. Expected Molecular weights for each VHPO with and without its N-terminal signal sequence follows: esVHPO (57 kDa, 55 kDa), lpVHPO (53 kDa, 50.5 kDa), raVHPO (54 kDa, 51 kDa), omVHPO (55 kDa, 52 kDa), thVHPO (55 kDa, 52 KDa), and HZ11-VHPO (55.6 kDa, 53 kDa).

**Figure S23: MCD assay of AQ-VHPOs demonstrates absence of diffusible hypobromous acid production**

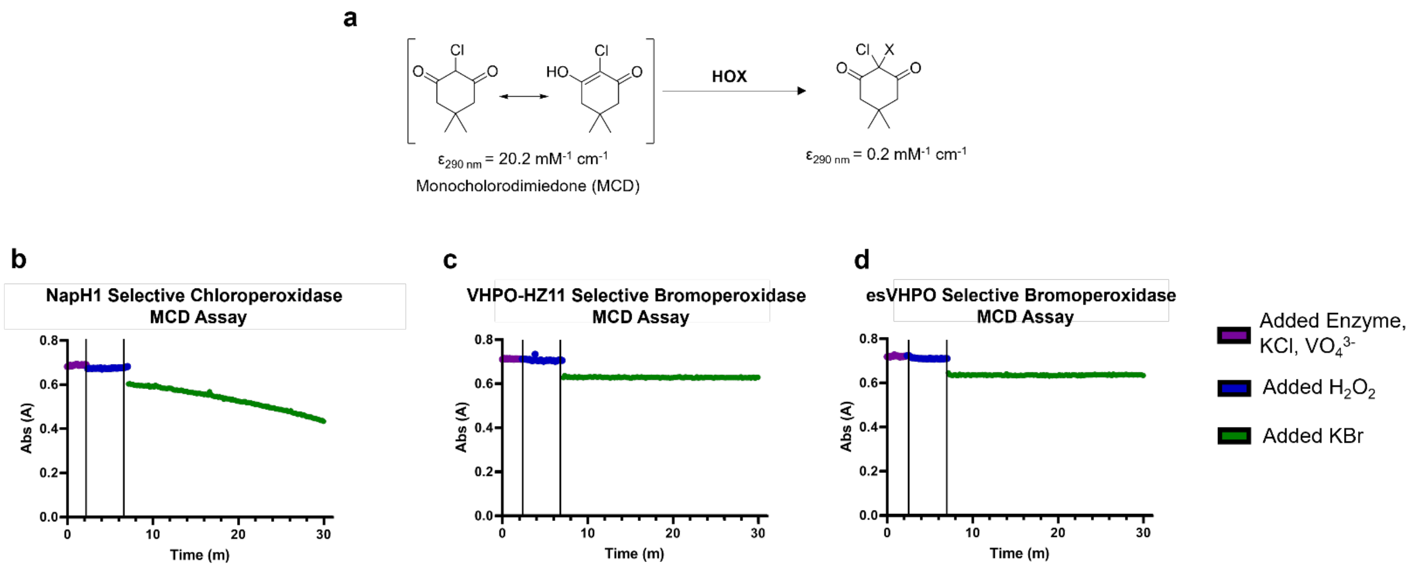


(a) Generalized reaction scheme for the monochlorodimedone (MCD) assay. Reactions were observed at 290 nm and the decrease of absorbance over time represents the release of hypohalous acid from the enzyme. (b) MCD assay of the previously characterized selective chloroperoxidase, NapH1, at pH 8.0. This was used as a positive control due to its ability to release hypobromous acid non-selectively. (c) MCD assay of the previously characterized HZ11-VHPO at pH 8.0. (d) MCD assay of esVHPO, characterized in this study, at pH 8.0. Each reaction was split into three time-dependent conditions and the change in data point color represents the point where the next substrate was added into the reaction.

**Figure S24: Co-substrate and co-factor dependency of purified AQ-VHPOs**

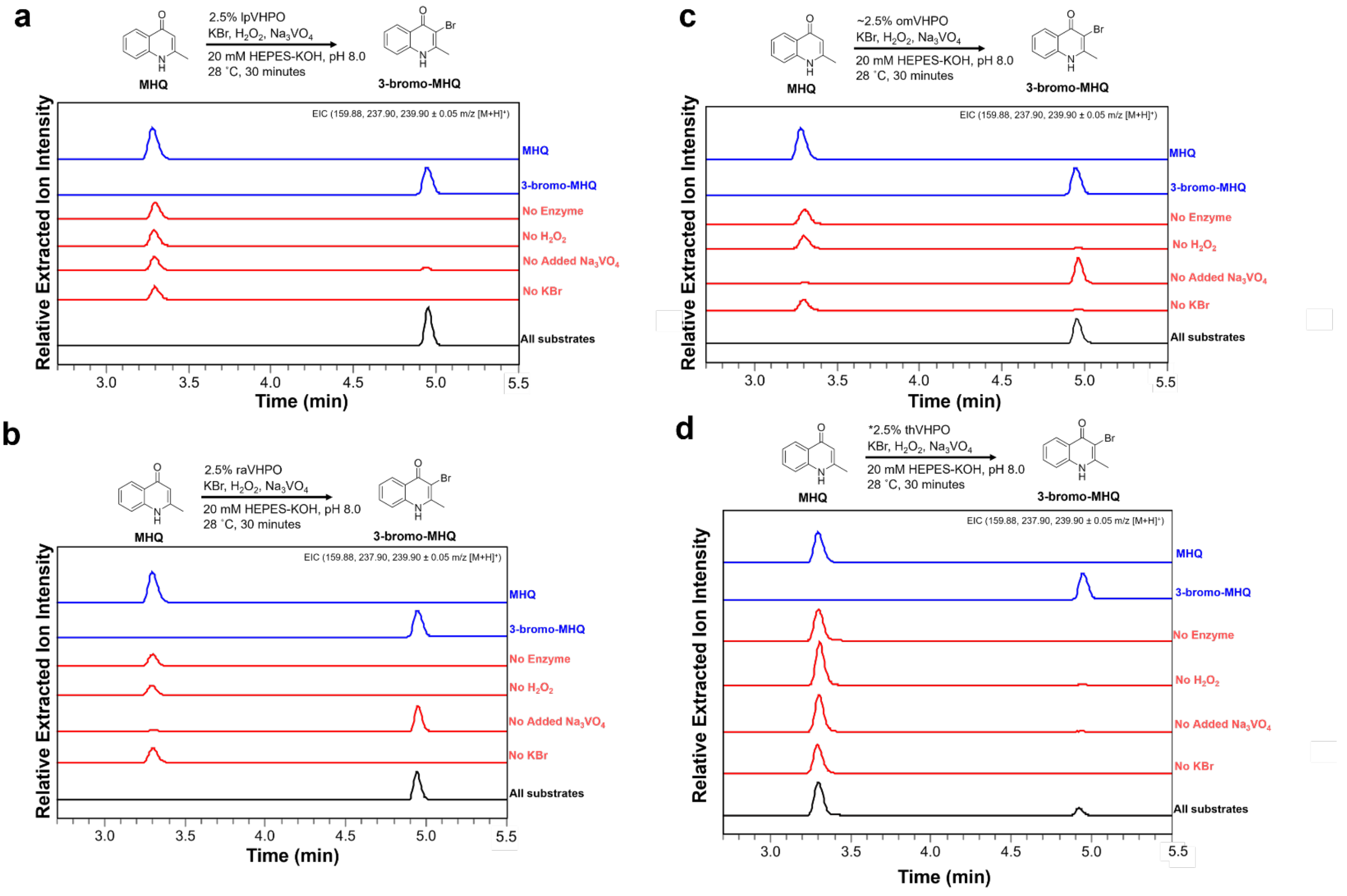


Co-substrate dependencies of lpVHPO (a), raVHPO (b), omVHPO (c), and thVHPO (d). Negative control traces follow the generalized reaction conditions, with the omission of its listed substrate from the reaction. Extracted ions are given in the upper right corner of each figure. The yield of thVHPO was low enough that concentration had to be roughly calculated and thus the precise molar ratio is likely to be less than the 2.5% based on the output concentration.

**Figure S25: IpVHPO control of MHQ bromination**

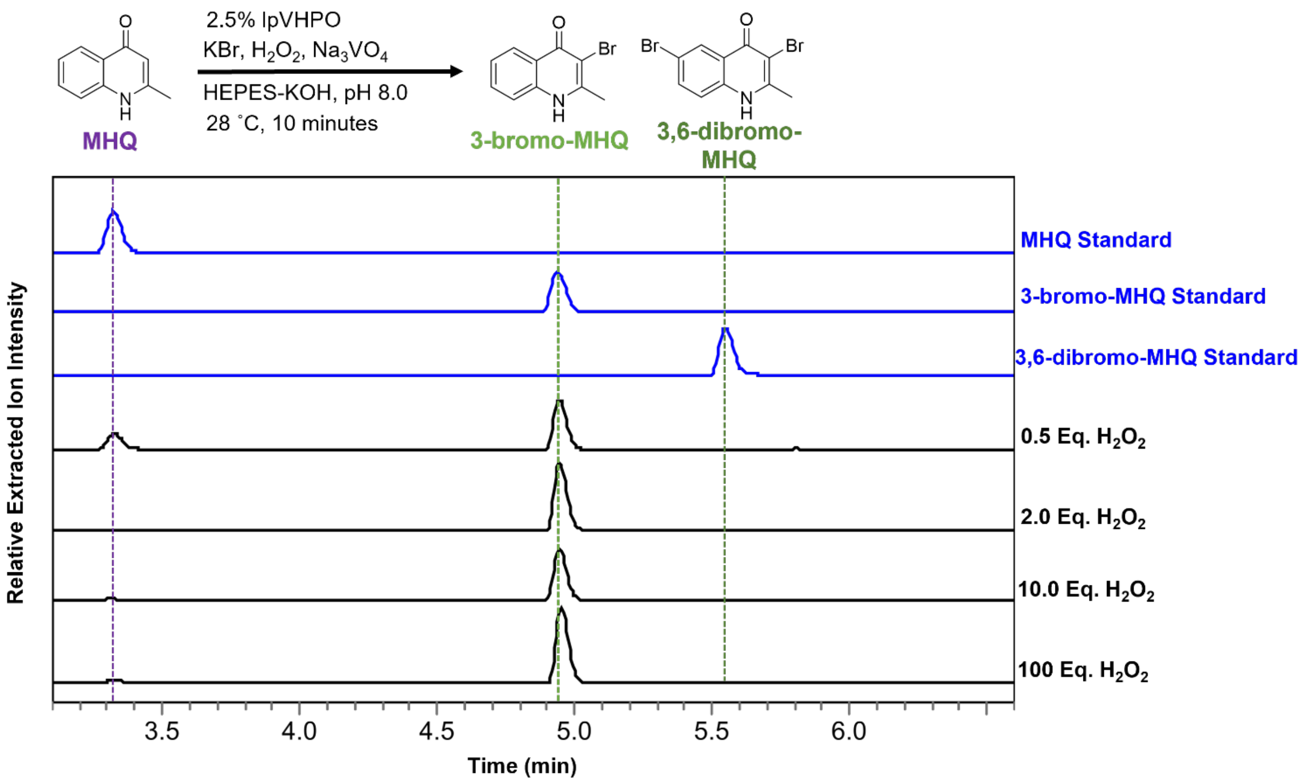


EIC traces of IpVHPO catalyzed MHQ bromination in the presence of increasing stoichiometric equivalents (compared to MHQ) of $H_2O_2$. Reaction conditions, substrate, and observed brominated products are given above the traces. Synthetic standard traces (blue) and experimental traces (black) are indicated. The following ions representing their species were extracted for all chromatogram traces: MHQ (160.07 ± 0.1 m/z $[M+H]^+$), Br-MHQ (237.98, 239.98 ± 0.1 m/z $[M+H]^+$), $Br_2$-MHQ (315.89, 317.89, 319.89 ± 0.1 m/z $[M+H]^+$), and $Br_3$-MHQ (393.80, 395.80, 397.80, 399.79 ± 0.1 m/z $[M+H]^+$). Each trace was acquired in 50:50 water:methanol.

**Figure S26: IpVHPO halide oxidation profile**

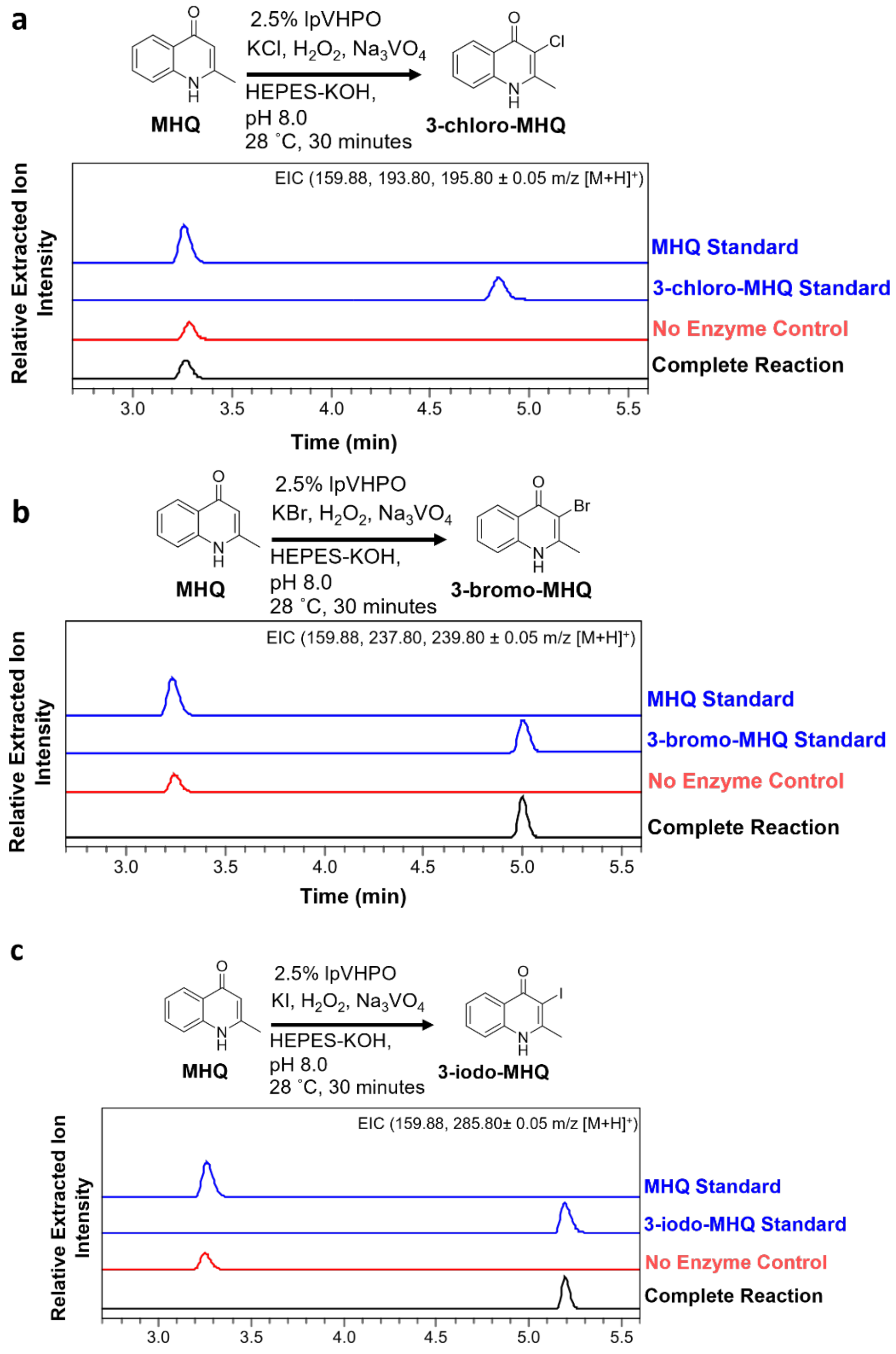


The ability for IpVHPO to oxidize each halide is shown. Extracted ion chromatogram (EIC) traces for each condition are shown with the extracted ions listed in the upper right of each trace. Generalized reaction conditions, AQ substrate, and products are given above EIC traces. Synthetic standard traces (blue), negative control traces (red), and experimental traces (black) are indicated. (a) Reaction of IpVHPO with potassium chloride as the halide source. (b) Reaction of IpVHPO with potassium bromide as the halide source. (c) Reaction of IpVHPO with potassium iodide as the halide source.

**Figure S27: raVHPO halide oxidation profile**

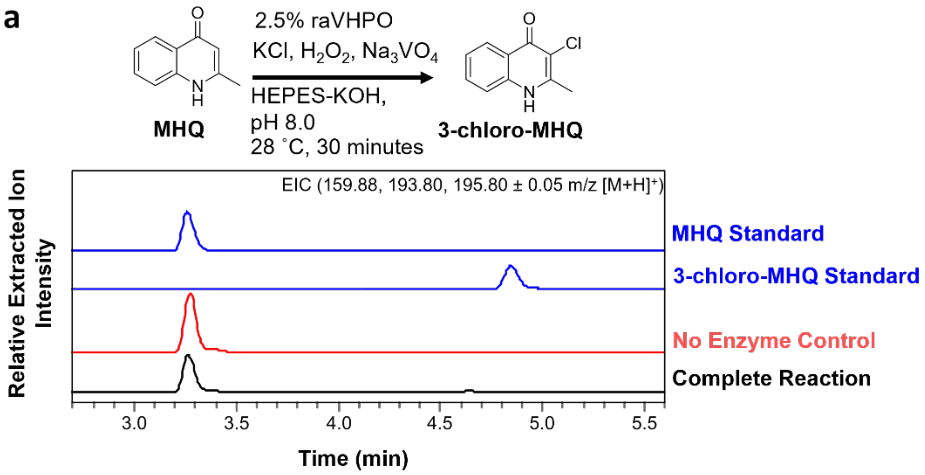


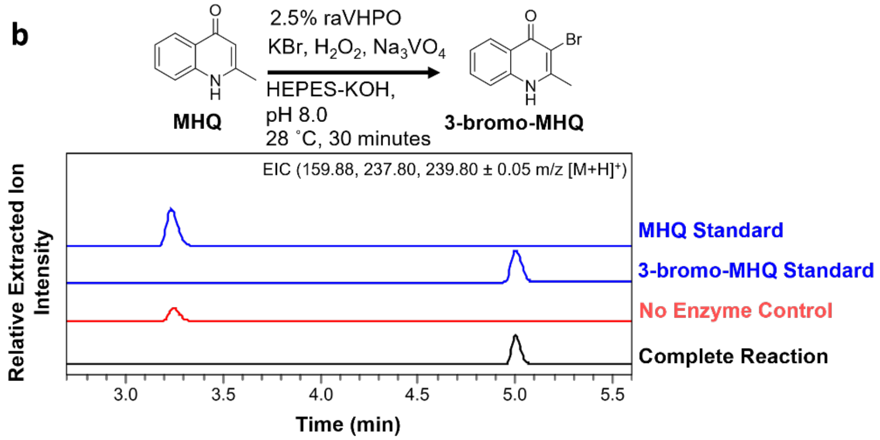


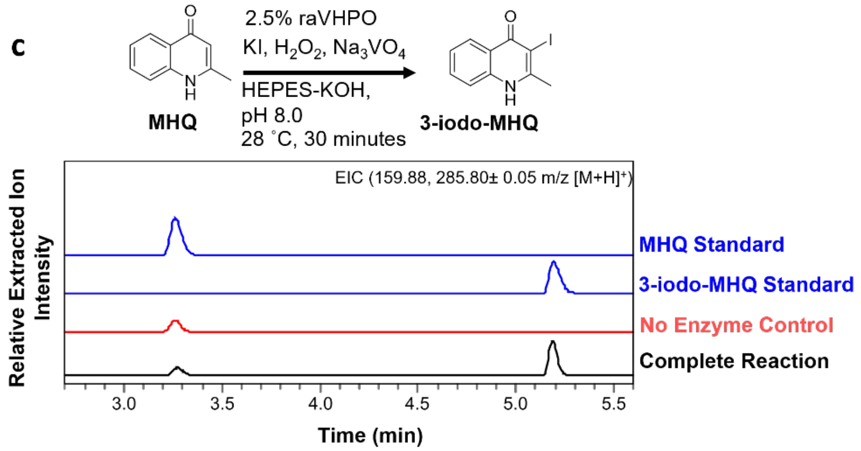


The ability for raVHPO to oxidize each halide is shown. All other components are as Figure S24. (a) Reaction of raVHPO with potassium chloride as the halide source. (b) Reaction of raVHPO with potassium bromide as the halide source. (c) Reaction of raVHPO with potassium iodide as the halide source.

**Figure S28: omVHPO halide oxidation profile**

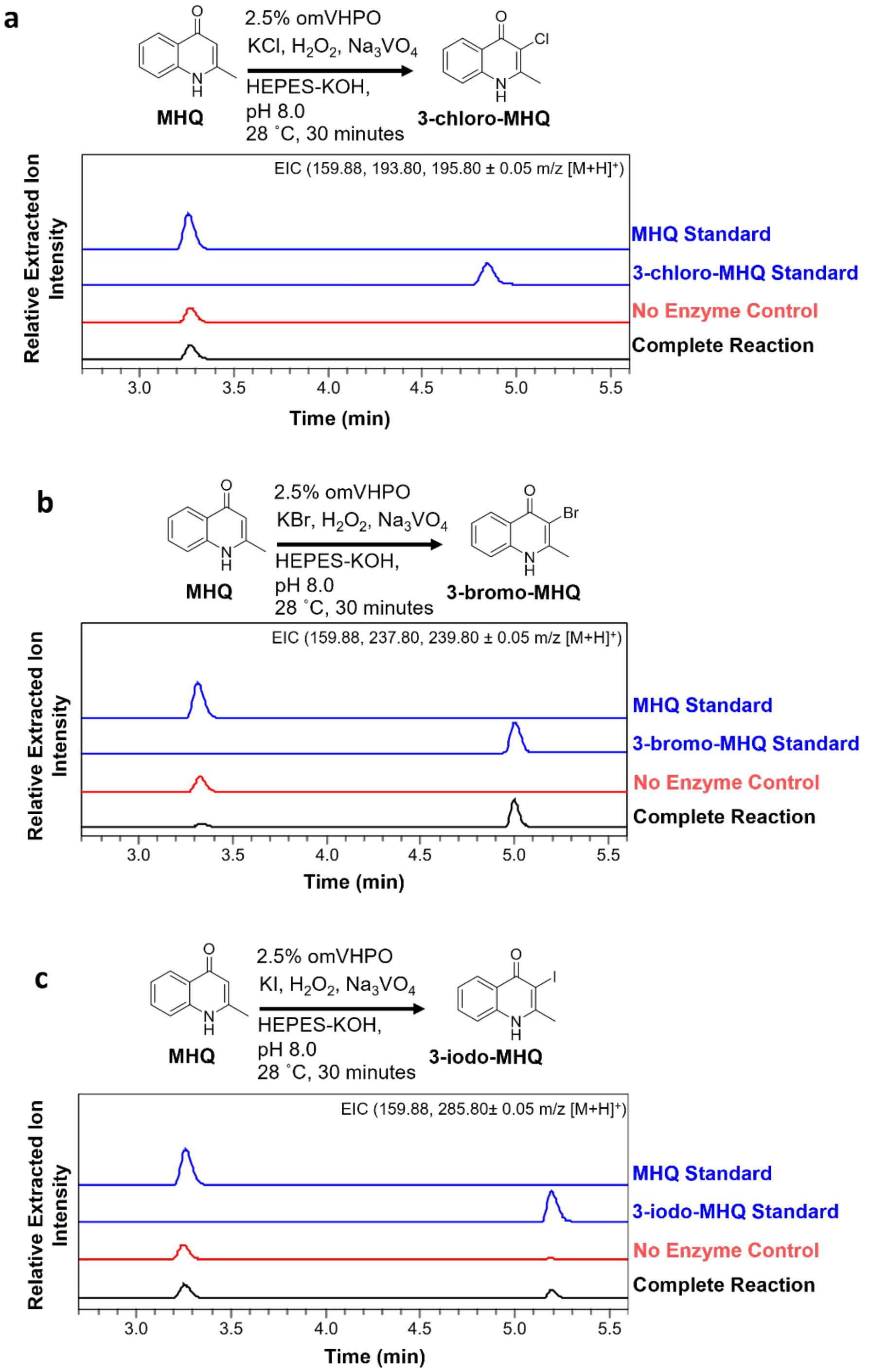


The ability for omVHPO to oxidize each halide is shown. All other components are as Figure S24. (a) Reaction of omVHPO with potassium chloride as the halide source. (b) Reaction of omVHPO with potassium bromide as the halide source. (c) Reaction of omVHPO with potassium iodide as the halide source.

**Figure S29: Response of *V. natriegens* CCUG 16374 to exogenous HHQ and Br-HHQ**

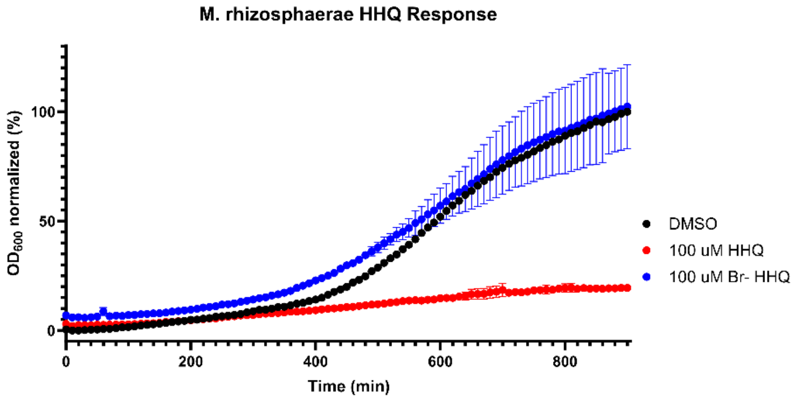


Response of *V. natriegens* to two concentrations of synthetic HHQ. Error bars represent the mean ± s.d. (n=3). Each [HHQ] growth curve is normalized to the lowest and highest points of the DMSO growth curve. Growth curve of *V. natriegens* is the presence of DMSO (black), 100 μM HHQ (red), and 100 μM Br-HHQ (blue).

**Figure S30: Response of *M. rhizosphaerae* to exogenous HHQ and Br-HHQ**

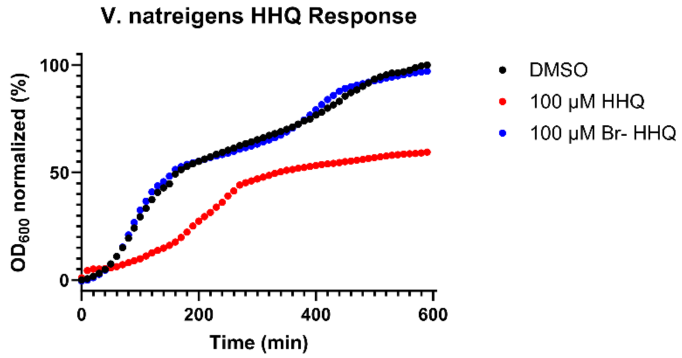


Response of *M. rhizosphaerae* to two concentrations of synthetic HHQ. Error bars represent the mean ± s.d. (n=3). Each [HHQ] growth curve is normalized to the lowest and highest points of the DMSO growth curve. Growth curve *M. rhizosphaerae* is the presence of DMSO (black), 100 μM HHQ (red), and 100 μM Br-HHQ (blue).

#### Figure S31: Response of *O. marinus* to exogenous MHQ and Br-MHQ

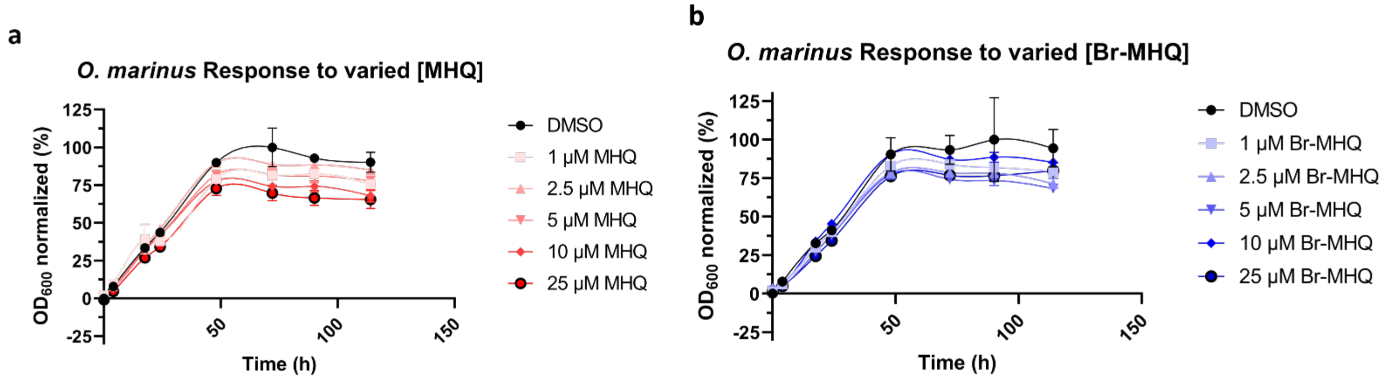


Response of *O. marinus* to the increasing concentration of synthetic MHQ and Br-MHQ. Error bars represent the mean ± s.d. (n=3). Lines do not represent a line of best fit and are added for visual clarity only. Each [MHQ] growth curve is normalized to the lowest and highest points of the DMSO growth curve. (a) Growth curve of *O. marinus* is the presence of DMSO and a gradient of MHQ concentrations. (b) Growth curve of *O. marinus* is the presence of DMSO and a gradient of Br-MHQ concentration.

**Figure S32: *In vivo* biotransformation of exogenous HHQ by O. marinus**

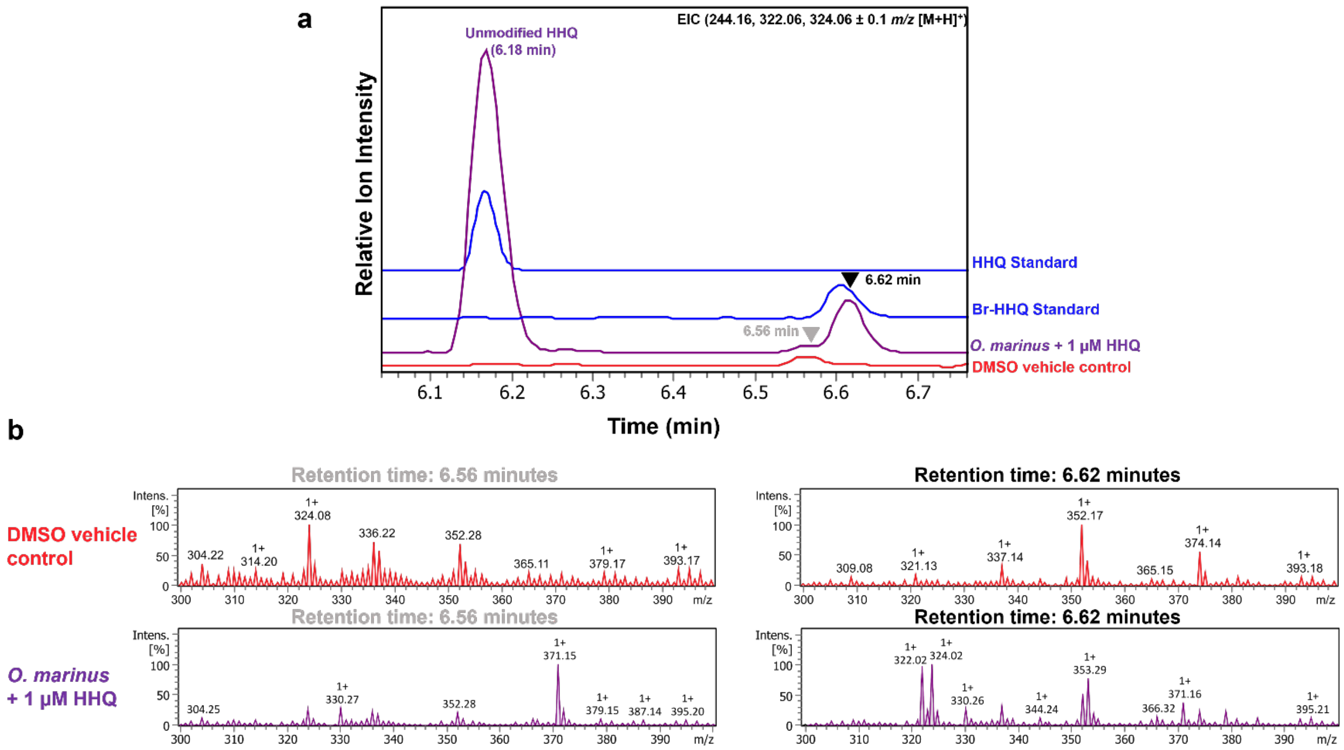


(a) EIC traces of *O. marinus* culture extracts treated with DMSO (red) and 1 µM HHQ (purple). DMSO and HHQ were added at the time of inoculation and cultures were extracted when stationary phase had been reached. 500 nM HHQ and 250 nM Br-HHQ standards (blue) are included for reference. Ions for HHQ and Br-HHQ were extracted and are given in the upper right corner. The HHQ mass ion peak at 6.18 minutes is shown for reference. The retention time for Br-HHQ is marked with a black arrow, while the retention time for a background 324.08 *m/z* is marked with a grey arrow. (b) Mass spectra for extracted ions at 6.56 minutes and 6.62 min from panel **a.** The characteristic isotopic distribution of brominated compounds is seen for the mass ion peak in *O. marinus* culture supplemented with HHQ is observed at 6.62 minutes which aligns with the synthetic standard.

**Figure S33: *O. marinus* omVHPO qPCR**

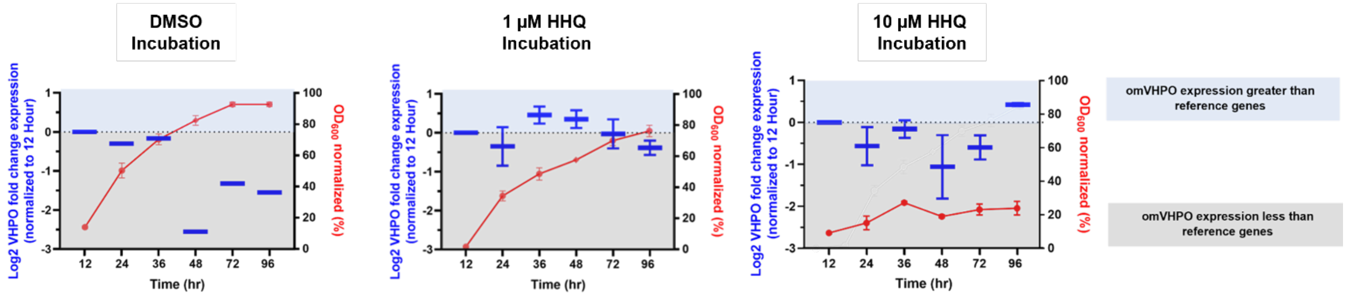


Box plots (blue) show the fold change of VHPO transcript expression in the presence of DMSO (left), 10 µM HHQ (middle) and 10 µM HHQ (right) compared to the initial transcription of the omVHPO gene at 12 hours of growth. The $OD_{600}$ of the culture at each time point is superimposed on the plot (red). Error bars represent the mean ± s.d. (n=3). VHPO expression was referenced to 16s rRNA and gyrA. The center box represents the mean change in fold expression and the error bars represent the s.d. (n=3 for 1 µM HHQ, n=2 for 10 µM HHQ).

**Figure S34**: ***O. marinus* growth curves for all HHQ concentrations in each media condition**

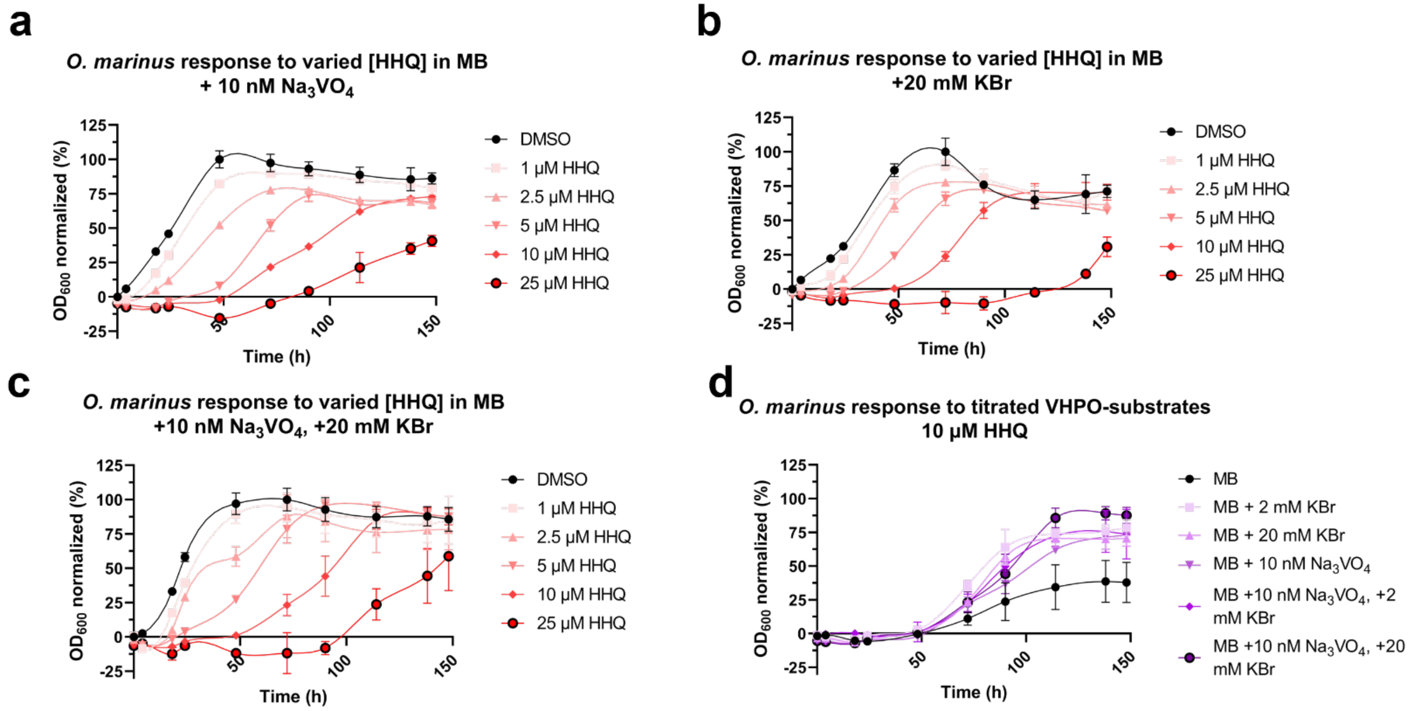


(a) Response of *O. marinus* in MMBP media to increasing concentrations of synthetic HHQ. Error bars represent the mean ± s.d. (n=3). Each [HHQ] growth curve is normalized to the lowest and highest points of the DMSO growth curve. In the following panels (b-c) each culture media was modified with differing amounts of VHPO co-substrate and tested against increasing concentrations of HHQ. Each growth curve was normalized against the DMSO growth curve of its media modification. (d), Full response of *O. marinus* to each media condition tested in the presence of 10 μM HHQ, as was partially depicted in Figure 5c.

**Figure S35: Optical images of *O. marinus* and *P. aeruginosa* PA14 co-cultures used for MALDI-MSI**

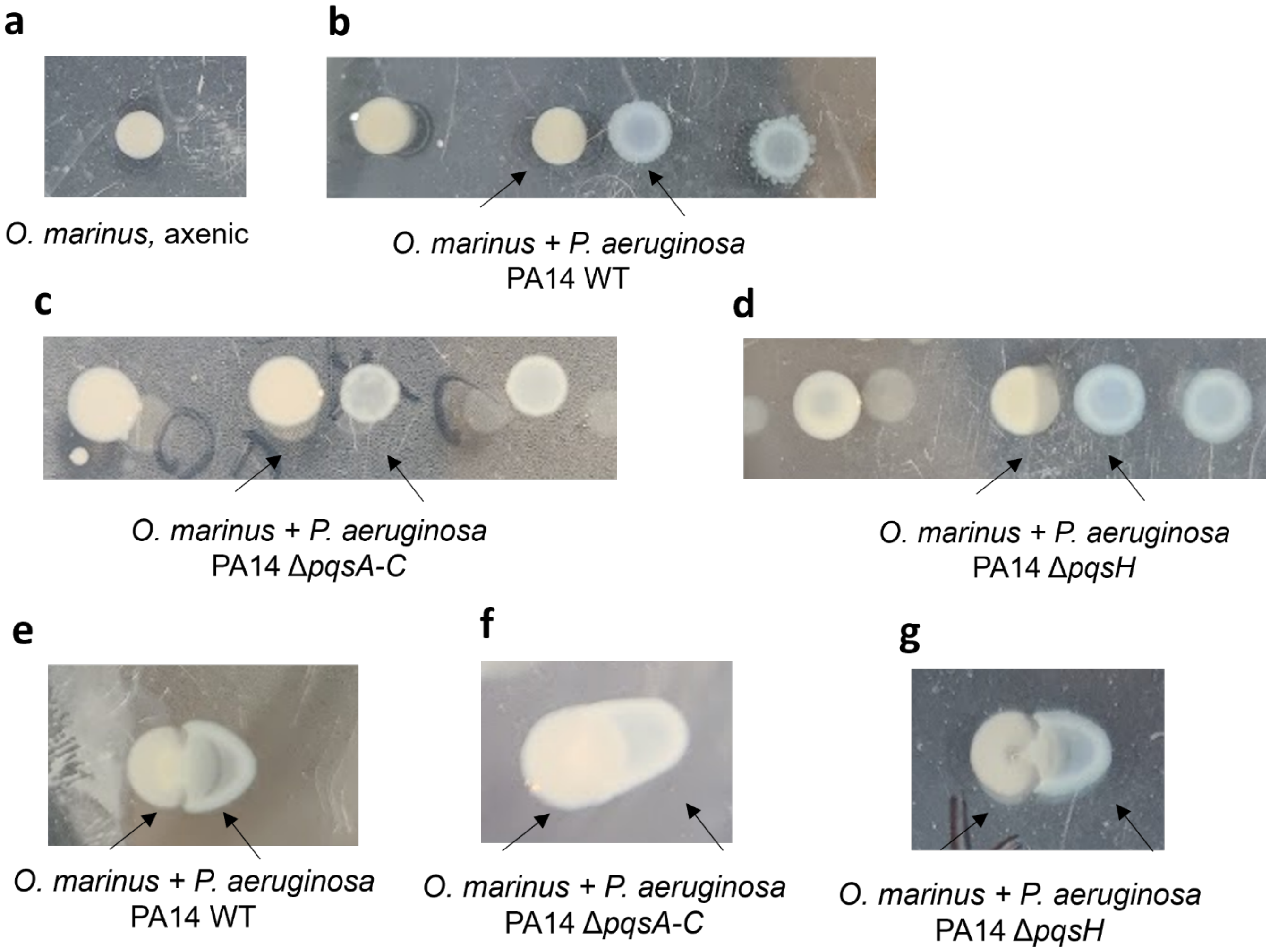


Optical images of co-cultures used for Figure 5 and Figure S36 prior to their excision from the plate. For all images, *O. marinus* colonies are on the left and *P. aeruginosa* PA14 colonies are on the right. Black arrows indicate where each colony sequence begins. (a) axenic *O. marinus* was spotted in absence of any *P. aeruginosa* PA14. Colonies of *P. aeruginosa* PA14 WT (b) *P. aeruginosa* PA14 ΔpqsA-C (c), and *P. aeruginosa* PA14 ΔpqsH (d) colonies were inoculated near, but not touching, previously established *O. marinus* colonies. Colonies of *P. aeruginosa* PA14 WT (e), *P. aeruginosa* PA14 ΔpqsA-C (f), and *P. aeruginosa* PA14 ΔpqsH (g) colonies were inoculated directly onto previously established *O. marinus* colonies.

**Figure S36: Additional experiments for *O. marinus* MSI**

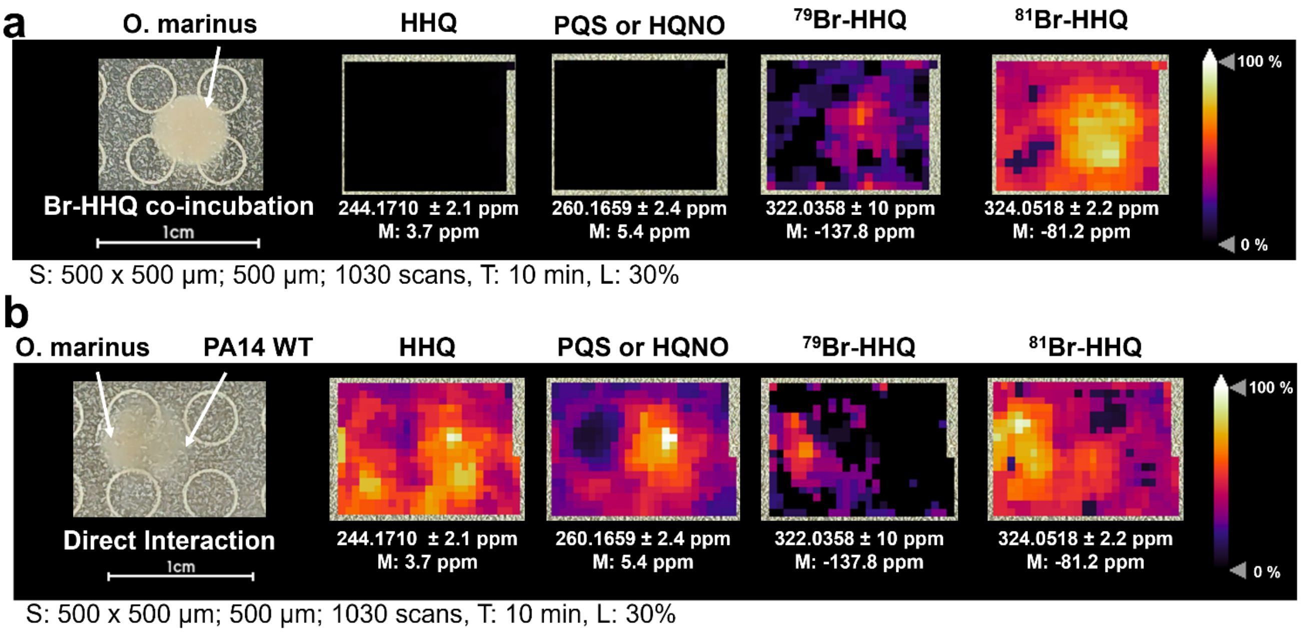


(a) The MALDI-QqTOF MSI visualization of axenic *O. marinus* supplemented with synthetic Br-HHQ. Ion images for HHQ, PQS/HQNO, and Br-HHQ were identified and visualized. (b) The MALDI-QqTOF MSI visualization of *O. marinus* co-cultured with wild type *P. aeruginosa* PA14, but with *P. aeruginosa* inoculated immediately next to *O. marinus*. For all panels, signal intensity is displayed as a heatmap and shows the spatial distribution of the ion image of interest. 1 cm scale bars are provided for each MSI collection. Spot raster; size; scan number (S), mass interval confidence (M), acquisition time (T), and laser power (L) are given in the figure. The annotation (A) for each experiment was targeted. the resolving power (R) of the instrument used for all MALDI-MSI data collection was $60000_{FWMH}$ at 1200 *m/z*.

**Figure S37: Analysis of contribution to $^{79}$Br-HHQ MALDI mass ion intensity by competing background peaks**

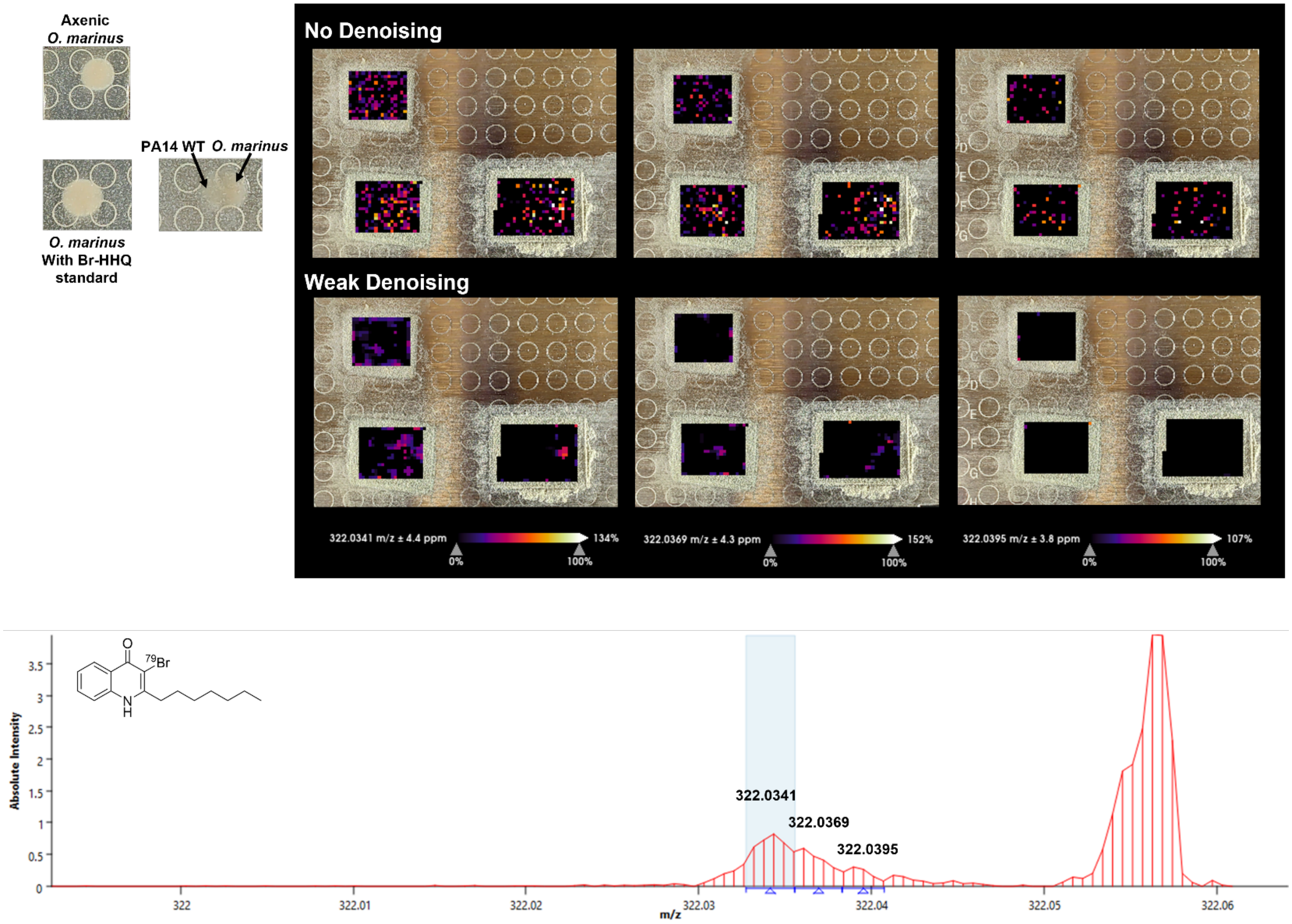


Deconvolution of the mass ion peak corresponding to $^{79}$Br-HHQ on the MALDI plate containing axenic *O. marinus*, axenic *O. marinus* incubated with 50 µM synthetically prepared Br-HHQ, and a co-culture of *O. marinus* and *P. aeruginosa* PA14 WT. The MALDI mass ion peaks for each of the three centered masses are shown both with no denoising algorithm applied to visualize low intensity ions and with a weak denoising algorithm to correspond with other MALDI-MSI images prepared in this work.

**Figure S38: Specific MS1 Spectra chosen for MALDI imaging of Figure 5d and Figure S35ab**

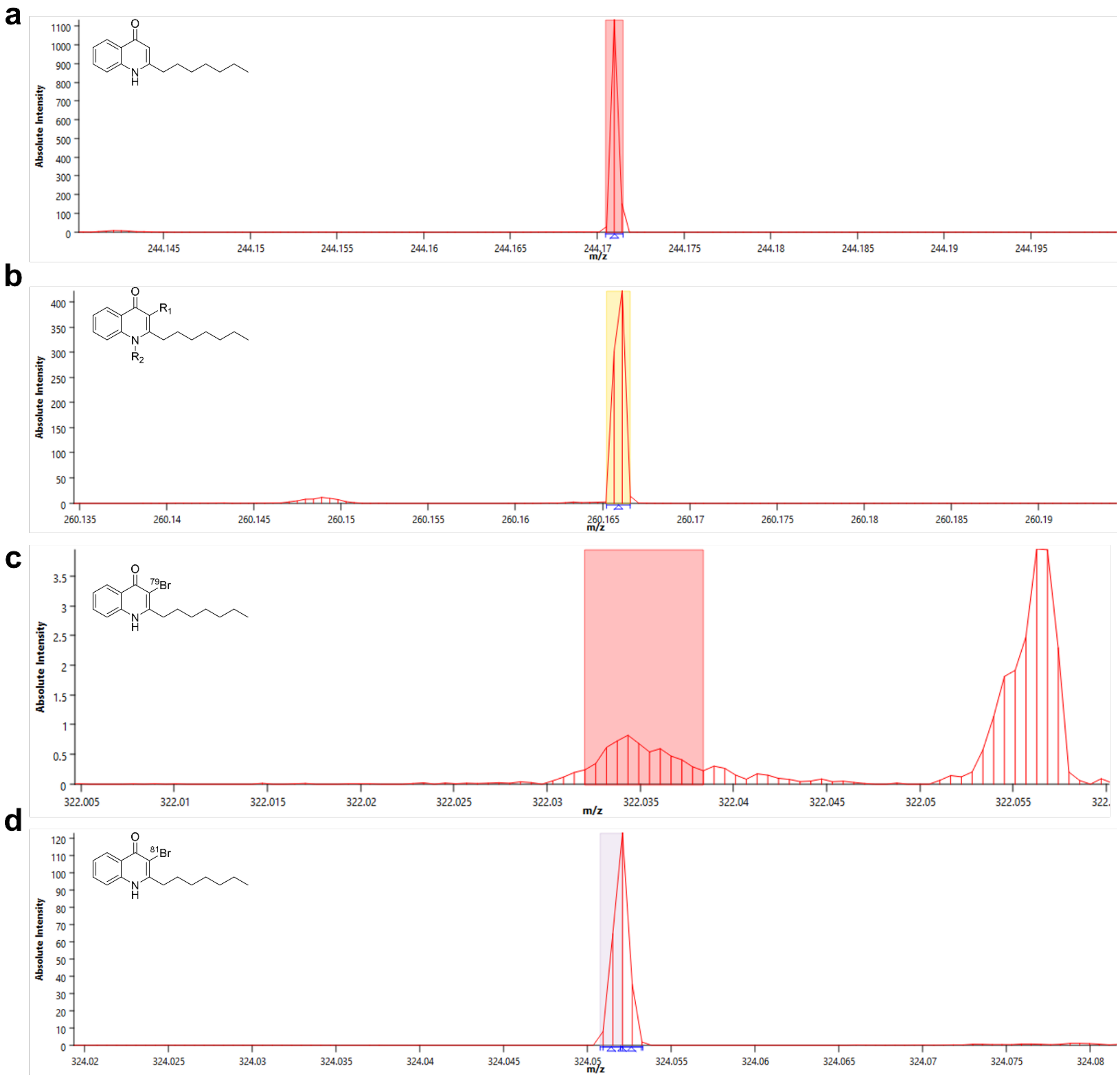


MS1 spectra for each analyzed mass ion on the MALDI plate containing the agar cut-outs of axenic *O. marinus*, axenic *O. marinus* incubated with 50 µM synthetically prepared Br-HHQ, and a co-culture of *O. marinus* and *P. aeruginosa* PA14 WT that had been inoculated for direct interaction. The colored boxes correspond to the region of the peak that was selected for mass ion visualization.

**Figure S39: Specific MS1 Spectra chosen for MALDI imaging of Figure 5e**

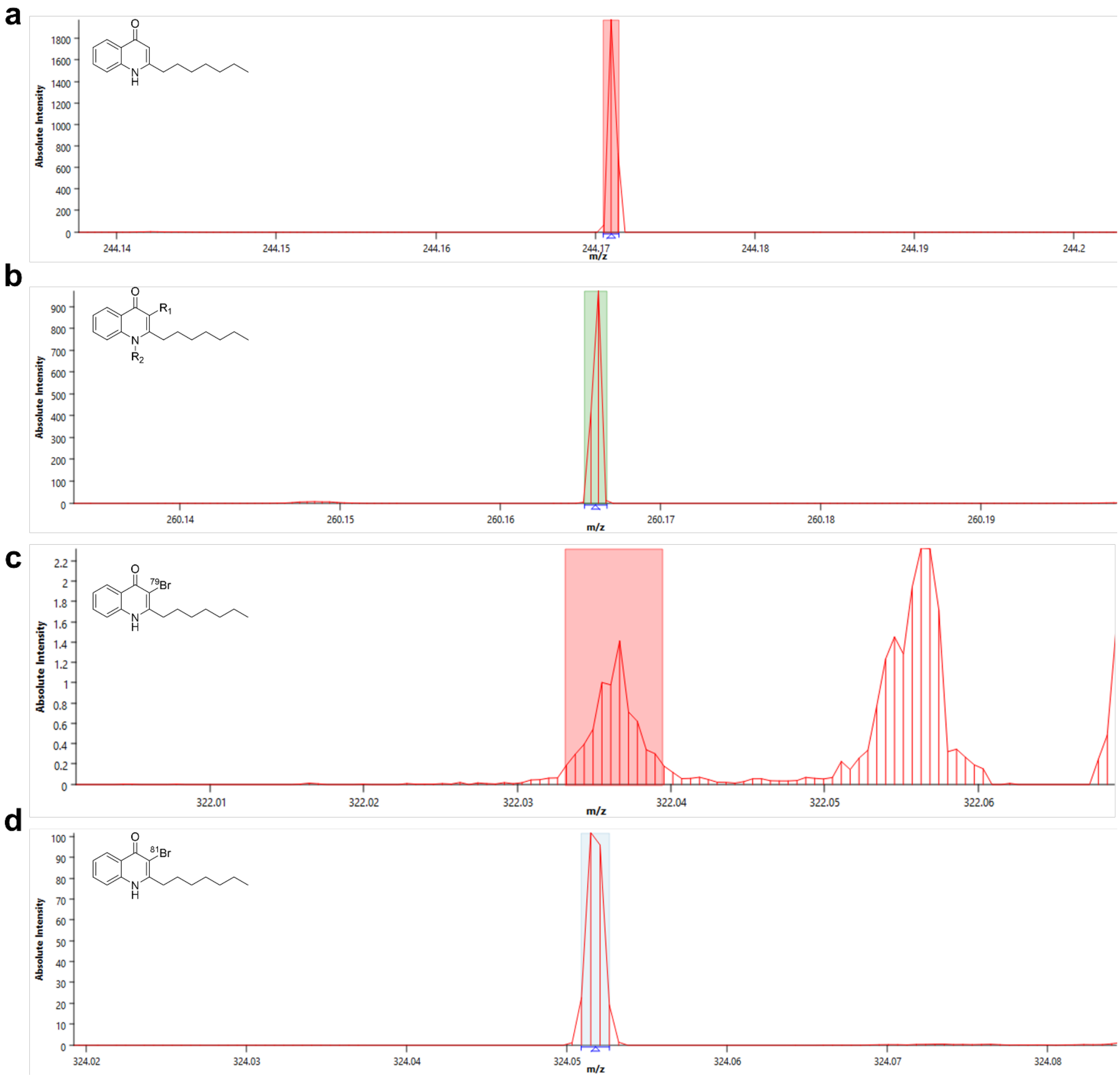


MS1 spectra for each analyzed mass ion on the MALDI plate containing the agar cut-out of the co-culture of *O. marinus* and *P. aeruginosa* PA14 WT that had been inoculated for interaction through agar diffusion. The colored boxes correspond to the region of the peak that was selected for mass ion visualization.

**Figure S40: Specific MS1 Spectra chosen for MALDI imaging of Figure 5fg**

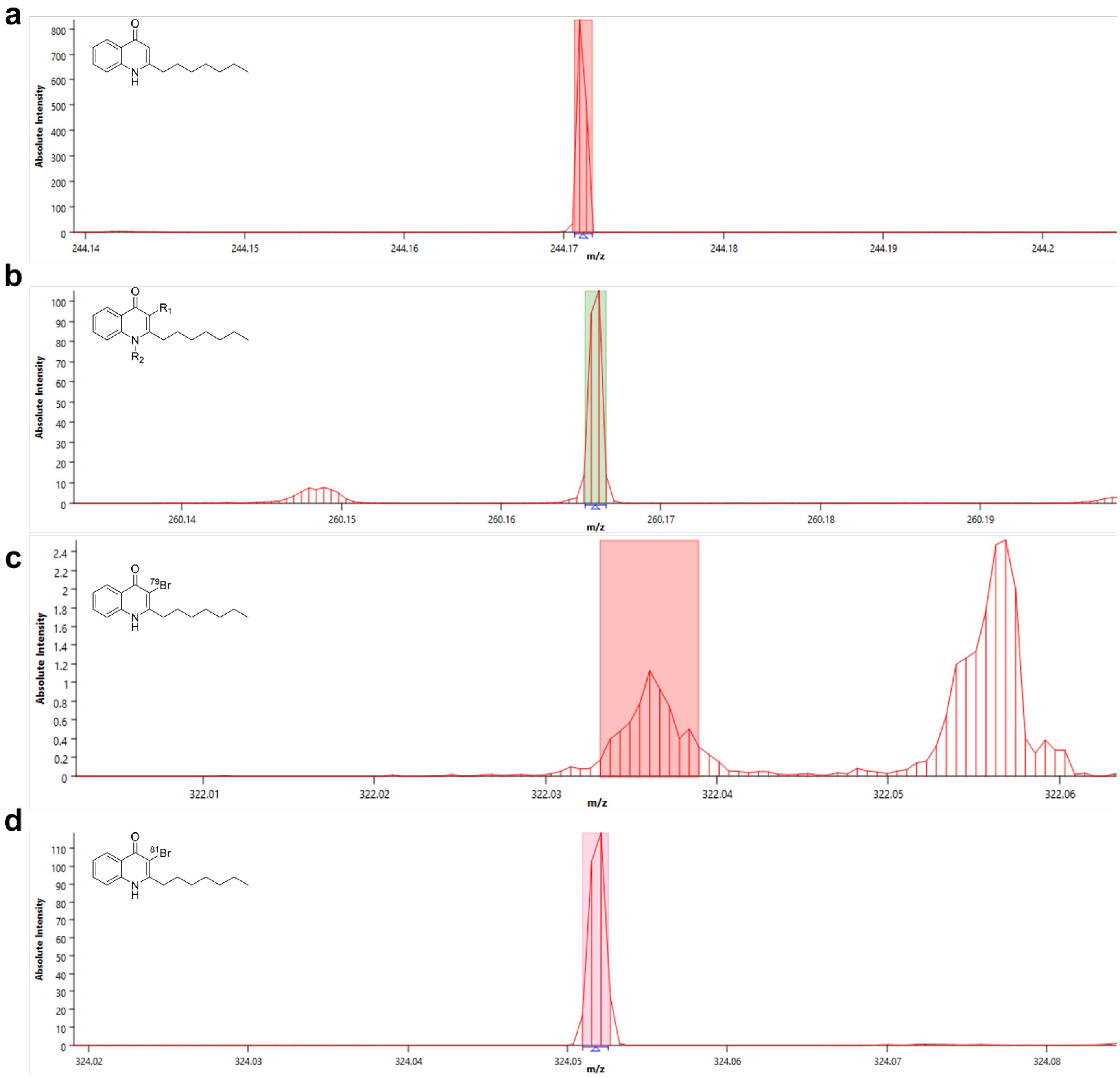


MS1 spectra for each analyzed mass ion on the MALDI plate containing the agar cut-out of the co-culture of *O. marinus* and *P. aeruginosa* PA14 ΔpqsA-C and the co-culture of *O. marinus* and *P. aeruginosa* PA14 ΔpqsH that had been inoculated for direct interaction. The colored boxes correspond to the region of the peak that was selected for mass ion visualization.

**Figure S41: Optical images of *V. natriegens* and *P. aeruginosa* PA14 co-cultures used for MALDI-MSI**

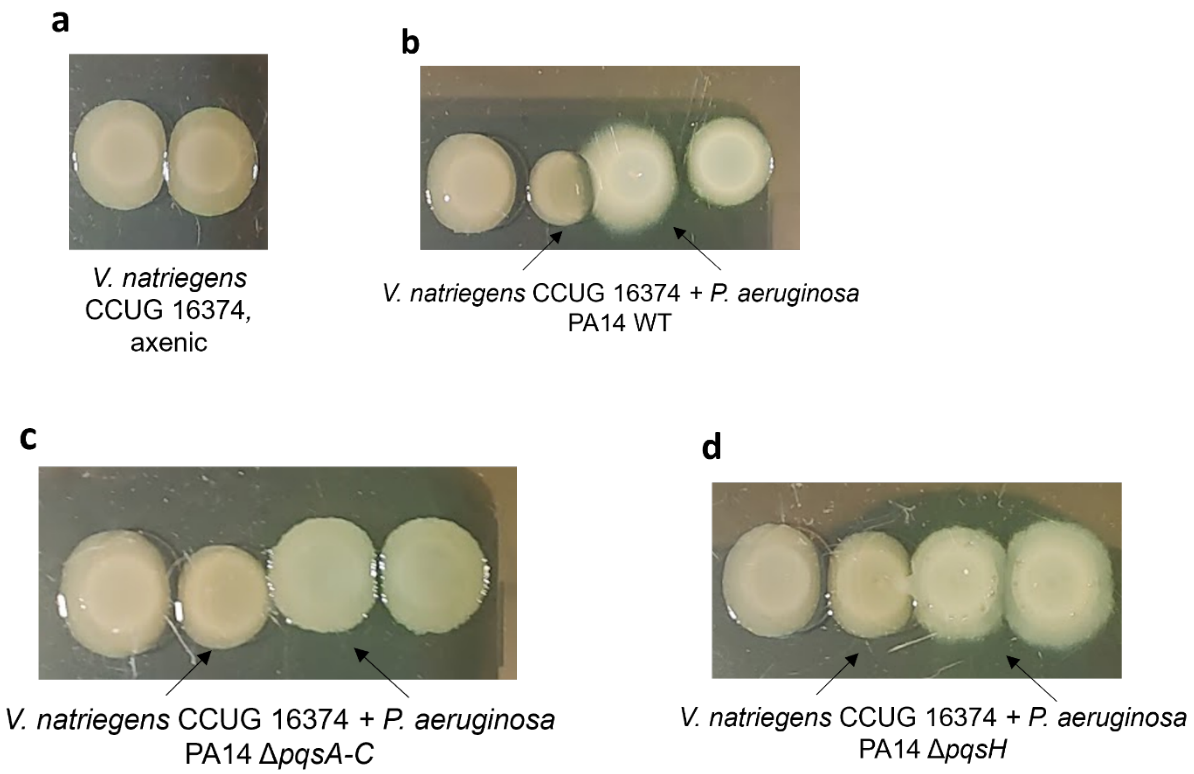


Optical images of co-cultures used for Figure S42 prior to their excision from the plate. For all images, *V. natriegens* colonies are on the left and *P. aeruginosa* PA14 colonies are on the right. Black arrows indicate where each colony sequence begins. (a) axenic *V. natriegens* was spotted in absence of any *P. aeruginosa* PA14. Colonies of *P. aeruginosa* PA14 WT (b) *P. aeruginosa* PA14 ΔpqsA-C (c) and *P. aeruginosa* PA14 ΔpqsH (d) colonies were inoculated directly next the inoculation for *V. natriegens*.

**Figure S42: Communal interaction between *V. natriegens* and *P. aeruginosa* PA14**

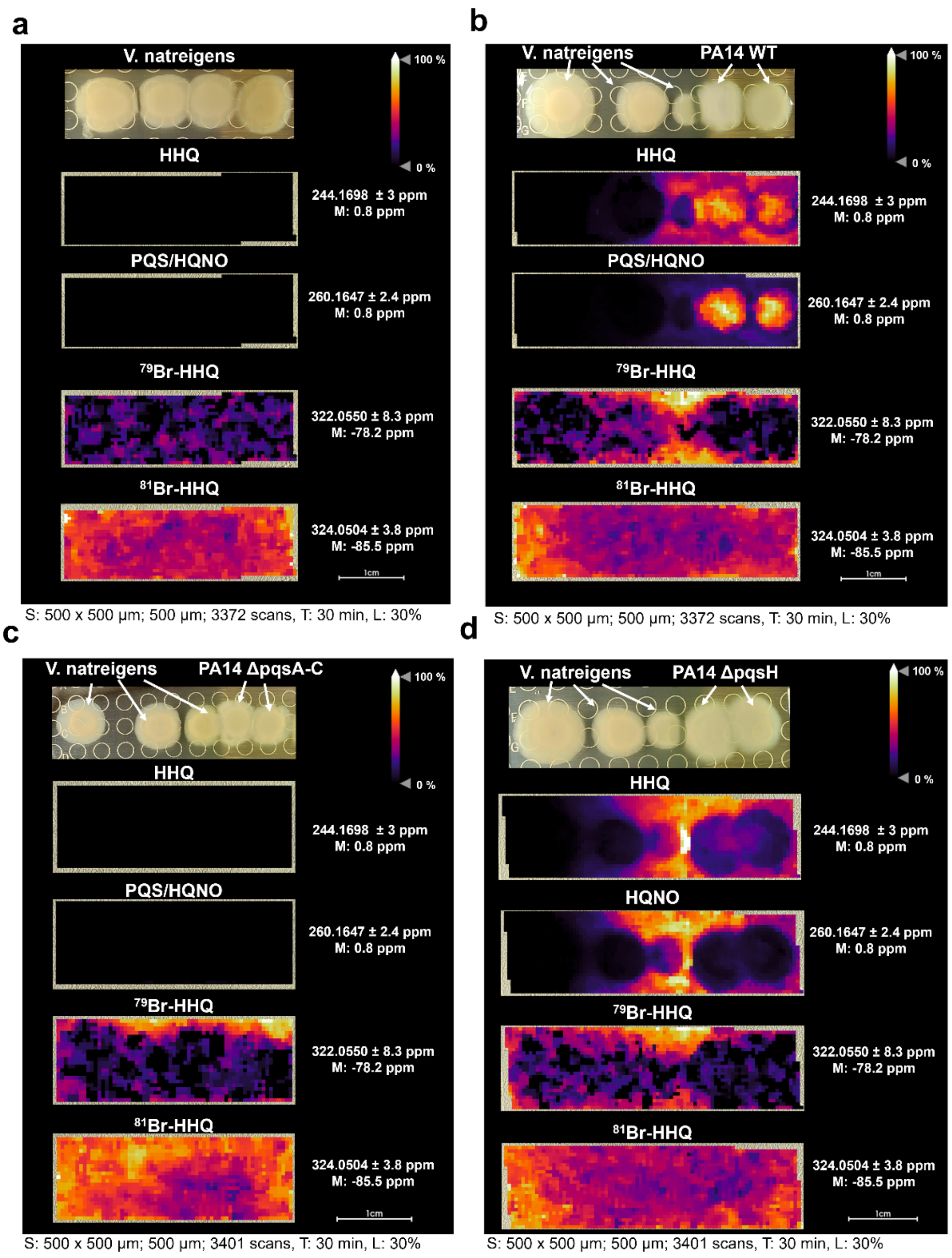


(a) The MALDI-QqTOF MSI visualization of axenic *V. natreigens*. Ion images of HHQ, PQS/HQNO, and Br-HHQ were identified and visualized. (b) The MALDI-QqTOF MSI visualization of *V. natreigens* co-cultured with wild type *P. aeruginosa* PA14. (c) The MALDI-QqTOF MSI visualization of *V. natreigens* co-cultured with the AQ knockout strain *P. aeruginosa* PA14 ΔpqsA-C. (d) The MALDI-QqTOF MSI visualization of *V. natreigens* co-cultured with the AQ knockout strain *P. aeruginosa* PA14 ΔpqsH.For all panels, signal intensity is displayed as a heatmap and shows the spatial distribution of the ion image of interest. 1 cm scale bars are provided for each MSI collection. Spot raster; size; scan number (S), mass interval confidence (M), acquisition time (T), and laser power (L) are given in the figure. The annotation (A) for each experiment was targeted. the resolving power (R) of the instrument used for all MALDI-MSI data collection was $60000_{FWMH}$ at 1200 *m/z*.

**Figure S43: Specific MS1 Spectra chosen for MALDI imaging of Figure S42 (Panels a and b)**

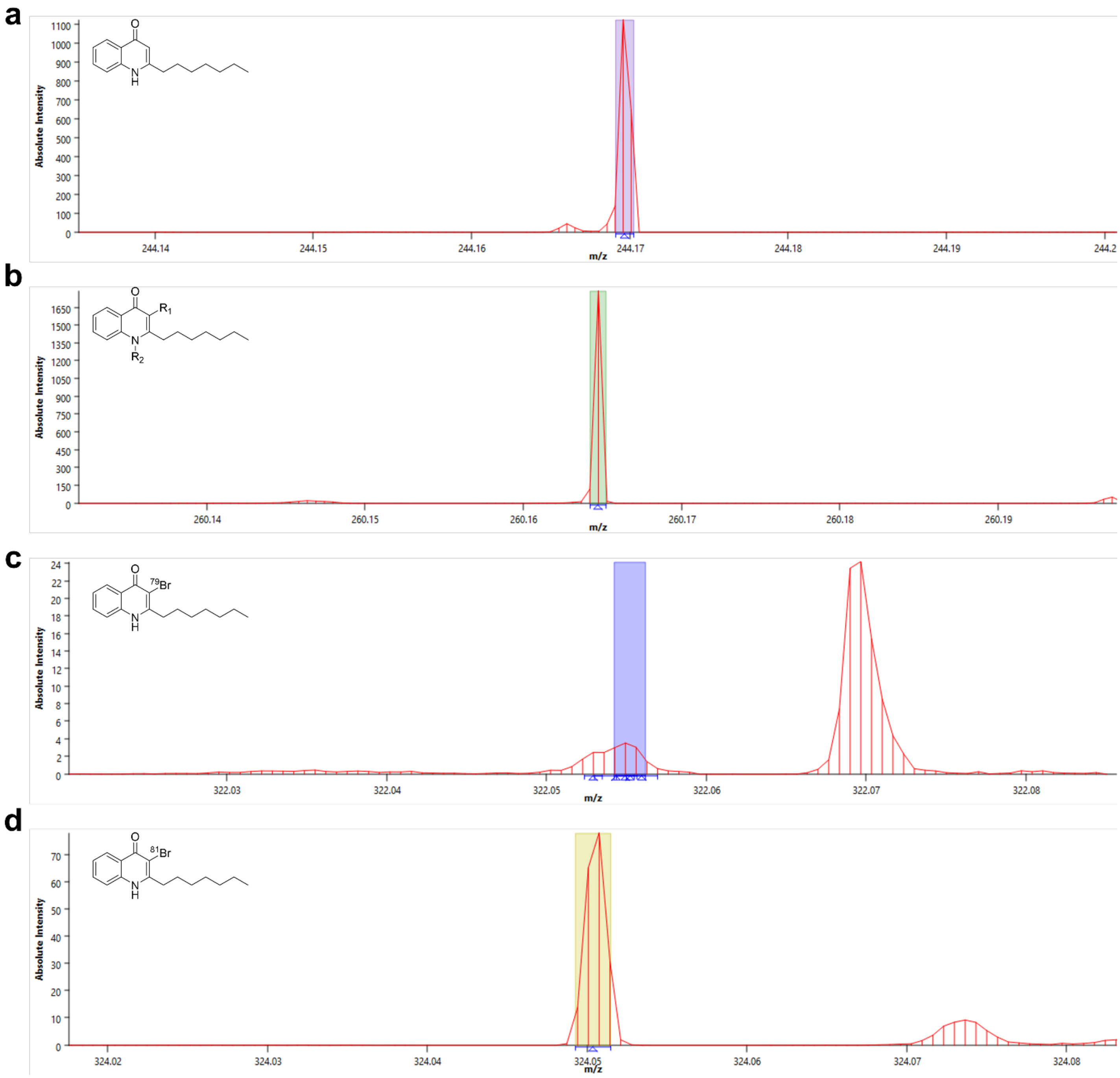


MS1 spectra for each analyzed mass ion on the MALDI plate containing the agar cut-outs of axenic *V. natreigens*, and a co-culture of *V. natreigens* and *P. aeruginosa* PA14 WT. The colored boxes correspond to the region of the peak that was selected for mass ion visualization.

**Figure S44: Specific MS1 Spectra chosen for MALDI imaging of Figure S42 (Panels c and d)**

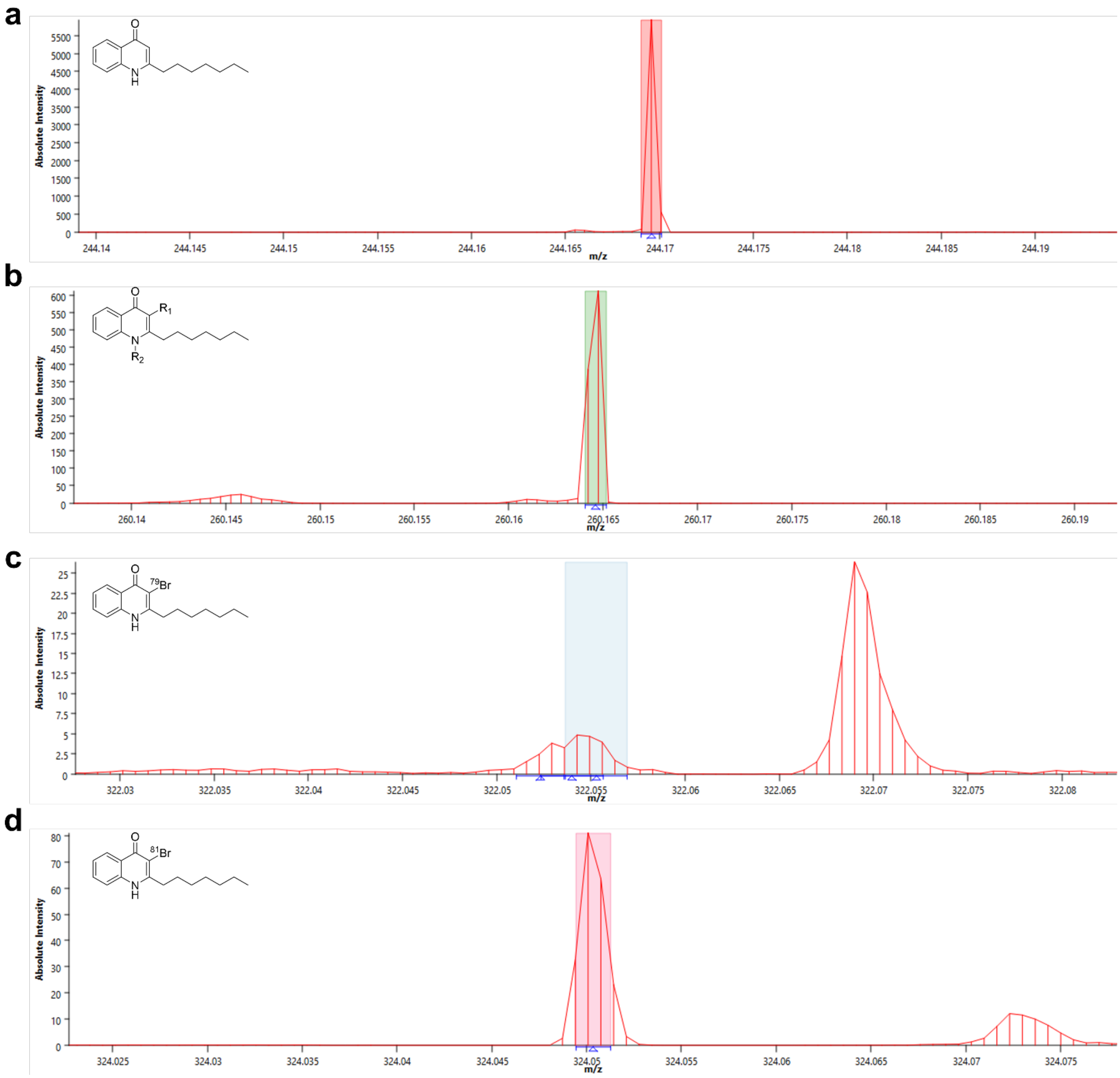


MS1 spectra for each analyzed mass ion on the MALDI plate containing the agar cut-outs of a co-culture of *V. natreigens* and *P. aeruginosa* PA14 ΔpqsA-C and a co-culture of *V. natreigens* and *P. aeruginosa* PA14 ΔpqsH. The colored boxes correspond to the region of the peak that was selected for mass ion visualization.

**Figure S45: Optical images of *M. rhizosphaerae* and *P. aeruginosa* PA14 co-cultures used for MALDI-MSI**

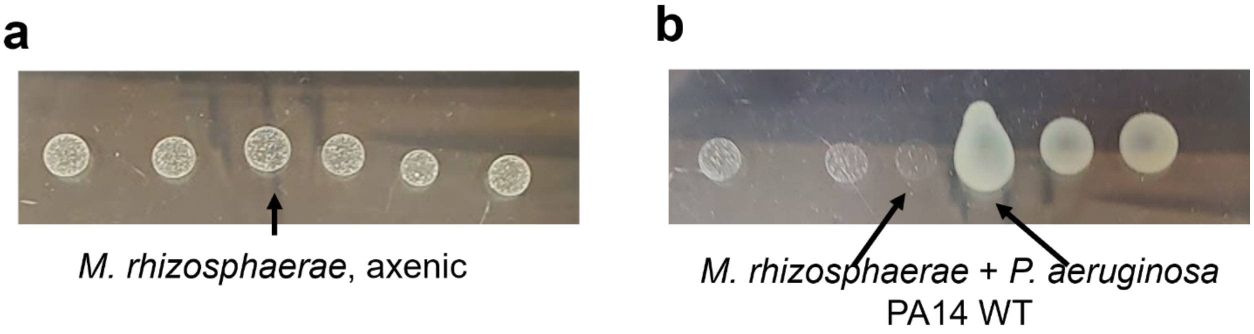


Optical images of co-cultures used for Figure S46 prior to their excision from the plate. For all images, *M. rhizosphaerae* colonies are on the left and *P. aeruginosa* PA14 colonies are on the right. Black arrows indicate where each colony sequence begins. (a) axenic *M. rhizosphaerae* was spotted in absence of any *P. aeruginosa* PA14. (b) *M. rhizosphaerae* co-spotted with colonies of *P. aeruginosa* PA14 WT

**Figure S46: Communal interaction between *M. rhizosphaerae* and *P. aeruginosa* PA14**

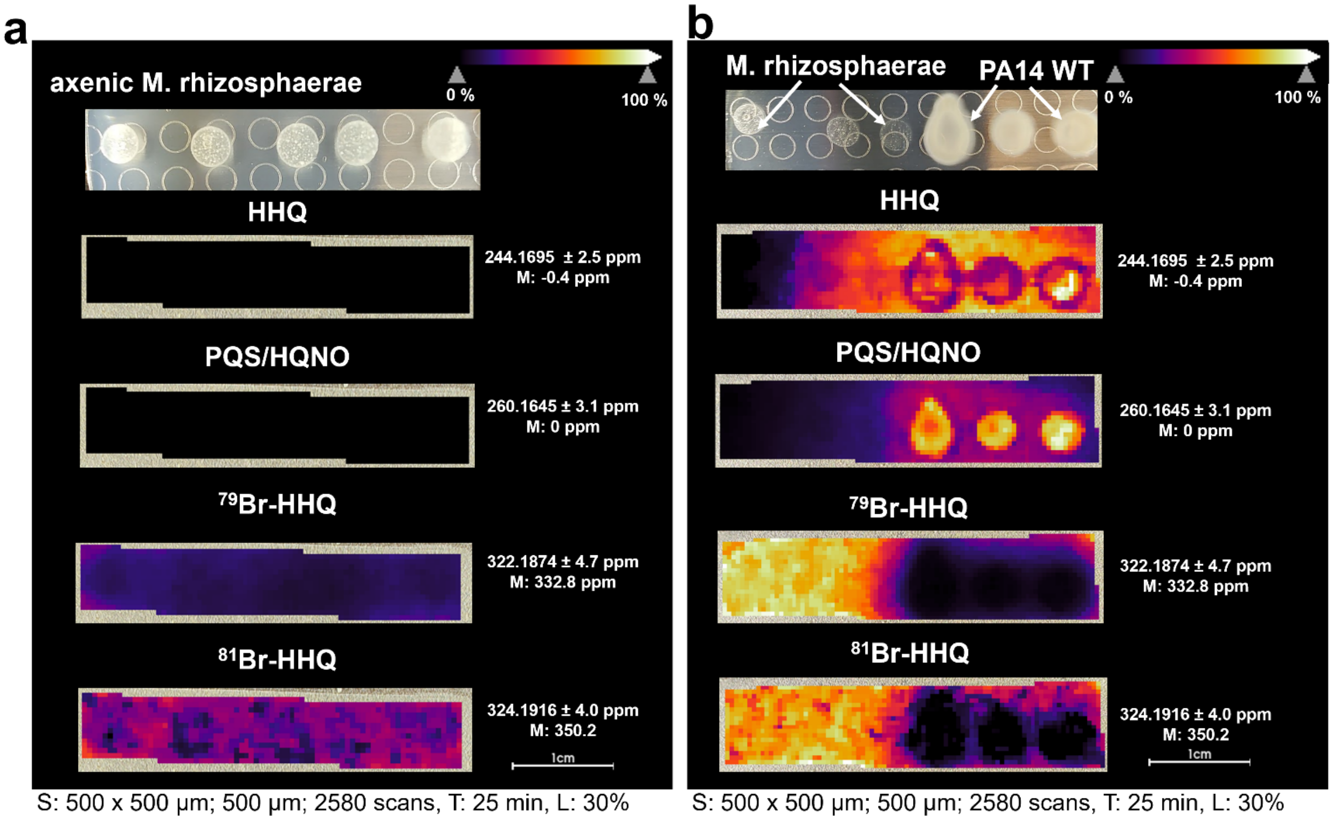


(a) The MALDI-QqTOF MSI visualization of axenic *M. rhizosphaerae*. Ion images of HHQ, PQS/HQNO, and Br-HHQ were identified and visualized. (b) The MALDI-QqTOF MSI visualization of *M. rhizosphaerae* co-cultured with wild type *P. aeruginosa* PA14. For all panels, signal intensity is displayed as a heatmap and shows the spatial distribution of the ion image of interest. 1 cm scale bars are provided for each MSI collection. Spot raster; size; scan number (S), mass interval confidence (M), acquisition time (T), and laser power (L) are given in the figure. The annotation (A) for each experiment was targeted. the resolving power (R) of the instrument used for all MALDI-MSI data collection was $60000_{FWMH}$ at 1200 *m/z*.

**Figure S47: Specific MS1 Spectra chosen for MALDI imaging of Figure S46**

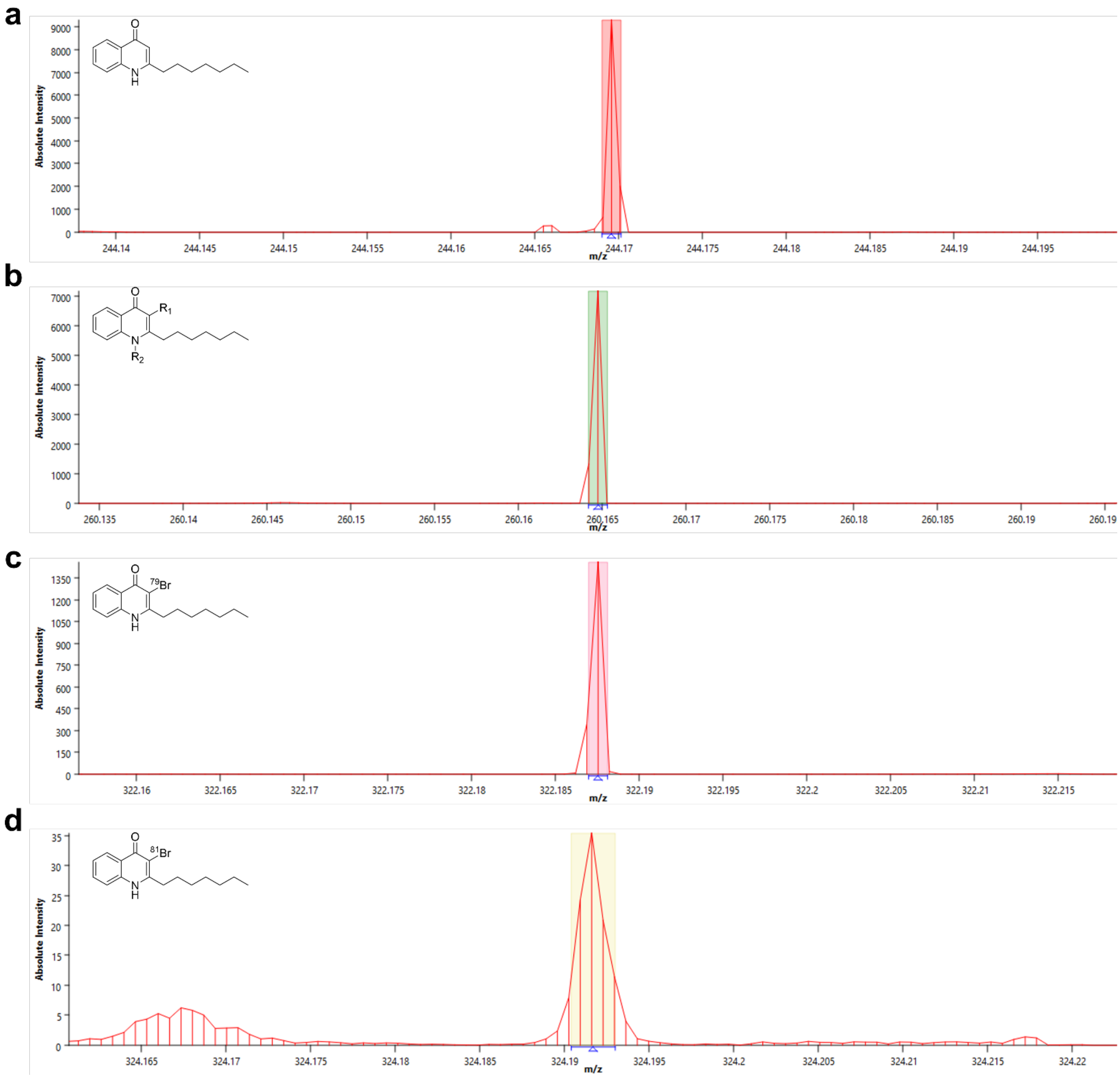


MS1 spectra for each analyzed mass ion on the MALDI plate containing the agar cut-outs of axenic *M. rhizosphaerae*, and a co-culture of *M. rhizosphaerae* and *P. aeruginosa* PA14 WT. The colored boxes correspond to the region of the peak that was selected for mass ion visualization.

**Figure S48: Optical images of *P. aeruginosa* PA14 Swarming motility**

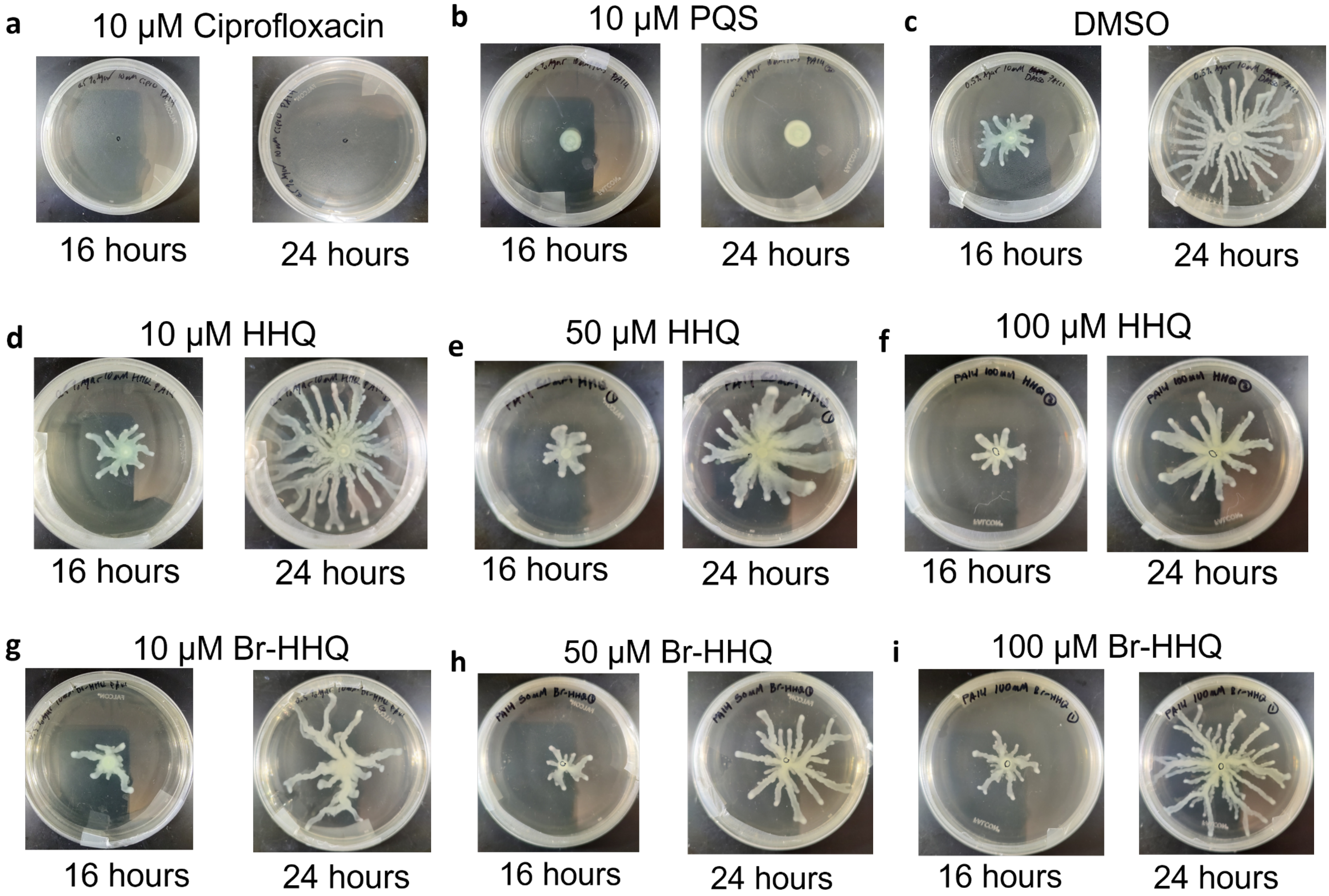

Optical images of *P. aeruginosa* PA14 swarms on 0.5% agar LB media. Images were taken at 16 hours and 24 hours of incubation at 37 °C. (a) Extent of swarming of *P. aeruginosa* PA14 in the presence of 10 μM ciprofloxacin (negative control) at 16 and 24 hours of incubation. (b) Extent of swarming of *P. aeruginosa* PA14 in the presence of 10 μM PQS (positive control) at 16 and 24 hours of incubation. (a) Extent of swarming of *P. aeruginosa* PA14 in the presence of DMSO (vehicle control) at 16 and 24 hours of incubation. Extent of swarming of *P. aeruginosa* PA14 in the presence of 10 μM HHQ (d), 50 μM HHQ (e), and 100 μM HHQ (f) at 16 and 24 hours of incubation. Extent of swarming of *P. aeruginosa* PA14 in the presence of 10 μM Br-HHQ (g), 50 μM Br-HHQ (h), and 100 μM Br-HHQ (**i**) at 16 and 24 hours of incubation.

**Figure S49: *P. aeruginosa* PA14 swarm areas**

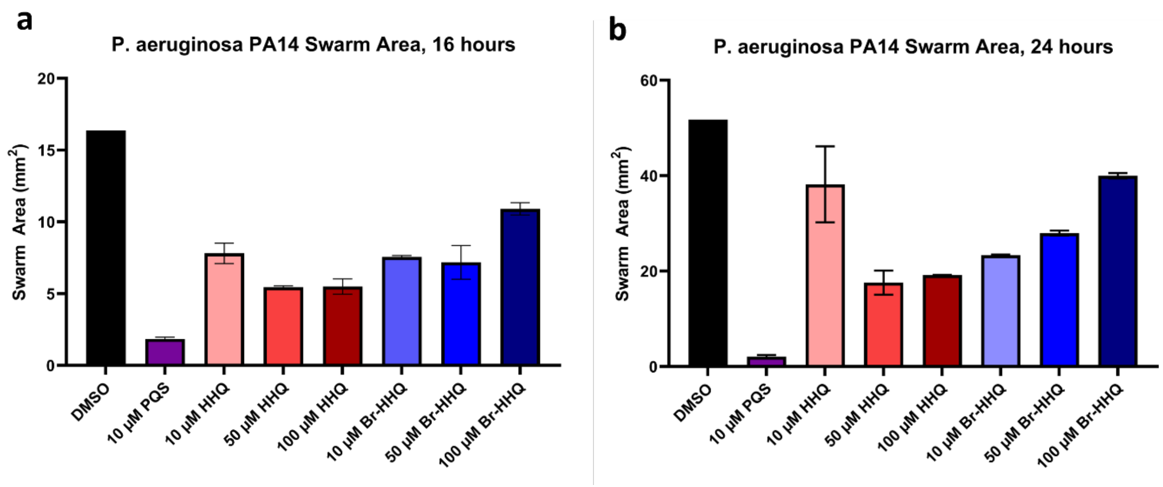


Optical images of *P. aeruginosa* PA14 swarms from Figure S48 were converted into areas ($mm^2$) using ImageJ. Swarm extent was measured following the perimeters of the swarm where it clearly transitioned from bacteria to unoccupied agar. Each swarm condition was performed in duplicate and the error bars represent the mean ± s.d. (n=2). (a) Bar chart of swarm areas for each condition at 16 hours. (b) Bar chart of swarm areas for each condition at 24 hours.

#### Chemical Synthesis

**2-heptyl-4(1H)-quinolone (HHQ)**

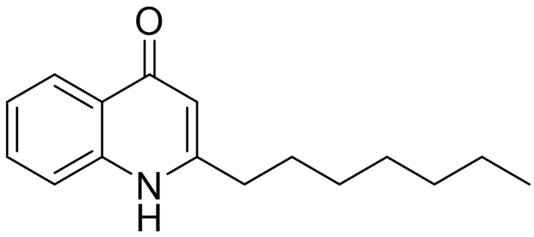

**HHQ** was prepared according to literature.[27] A reaction of ethyl 3-oxodecanoate (6.8 g, 34 mmol, prepared according to literature[28]), aniline (3.3 mL, 35.7 mmol), p-toluene sulfonic acid (0.12 g, 0.70 mmol) and hexanes (150 mL) yielded HHQ as a cream-colored solid (0.23 g, 0.95 mmol).

$^1$H NMR (499 MHz, 90% $CDCl_3$/10% $CD_3OD$) δ 8.21 (d, *J* = 8.2 Hz, 1H), 7.54 – 7.49 (m, 1H), 7.41 (d, *J* = 8.3 Hz, 1H), 7.28 (m,1H), 2.87 – 2.81 (m, 2H), 1.67 (p, *J* = 7.7 Hz, 2H), 1.36 – 1.17 (m, 8H), 0.81 – 0.76 (m, 3H). HRMS (MALDI-qQToF) Calculated exact mass for $C_{16}H_{22}NO$ 244.1701 (100%), found 244.1702 (100%) (M+H)

**2-heptyl-3-bromo-4(1H)-quinolone (Br-HHQ)**

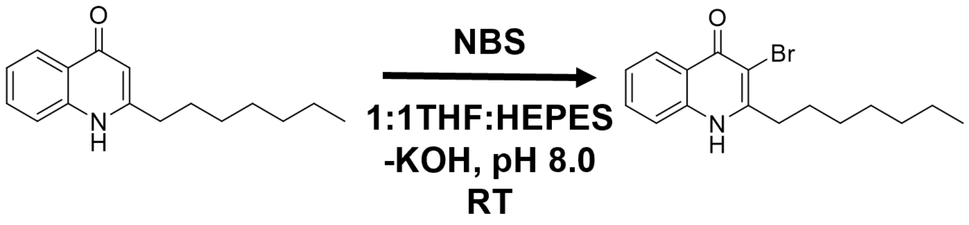

Method was adapted from previously established protocol[27] with additional adaptations inspired by aqueous enzymatic conditions.[19] Recrystallized *N*-bromosuccinimide (0.040 g, 0.226 mmol, 1.1 equiv) was dissolved in a THF (20 mL) and a 50 mM HEPES (pH 8.0, 20% glycerol) solution (20 mL). HHQ (0.050 g, 0.205 mmol, 1.0 equiv) was added to the solution and sonicated to completely dissolve. Reaction was stirred at 28 ˚C for 16 hours. Reaction progress was monitored with LC-MS analysis. The reaction was stopped by removing the solvent *in vacuo*. The solid residue was re-dissolved in 50 mL water and was washed with DCM (3 x 50 mL). The organic layer was then washed with water (2 x 150 mL) and brine (1 x 150 mL), dried over magnesium sulfate, and the solvent removed *in vacuo* resulting in an off-white powder (0.031 g, 0.096 mmol, 47% yield). No additional purification was required.

$^1$H NMR (499 MHz, 90% $CDCl_3$/10%$CD_3OD$) δ 8.21 (d, *J* = 8.2 Hz, 1H), 7.54 – 7.49 (m, 1H), 7.41 (d, *J* = 8.3 Hz, 1H), 7.28 (m,1H), 2.87 – 2.81 (m, 2H), 1.67 (p, *J* = 7.7 Hz, 2H), 1.36 – 1.17 (m, 8H), 0.81 – 0.76 (m, 3H). HRMS (MALDI-qQTOF; positive mode; 50:50 CHCA:DHB matrix) Calculated exact mass for $C_{16}H_{21}BrNO$ 322.0807 [$(M+H)^+$]. Observed 322.0805 [$(M+H)^+$].

**2-methyl-3-chloro-4(1H)-quinolone (Cl-MHQ)**

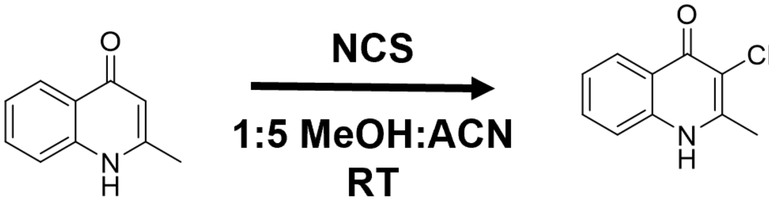

Method was adapted from previously established protocol.[29] A solution of MHQ (0.050 g, 0.31 mmol, 1.0 equiv) in acetonitrile (10 mL) was cooled to 0 ºC. Recrystallized *N*-chlorosuccinimide (0.105 g, 0.78 mmol, 2.5 equiv) was added all at once. Methanol (2 mL) was added to reaction to maintain solubility of MHQ and NCS. Reaction was warmed to room temperature and stirred for 24 hours. The reaction was concentrated *in vacuo* and resuspended in dichloromethane (25 mL) and water (25 mL). The layers were separated, and the aqueous layer was further washed with DCM (3 x 50 mL). The pooled organic layer was then washed with brine, dried over magnesium sulfate, and concentrated *in vacuo*. Reaction product was purified with semi-preparative RP-HPLC (Phenomenex Luna C8 (2), 10.0 x 250 mm) at a flow rate of 3.0 mL/min with a 10-100% solvent B gradient (solvent A: 0.1% formic acid; solvent B: acetonitrile + 0.1 % formic acid) over 30 minutes. One major peak was collected at 10.6 minutes, corresponding to a singly-chlorinated species. Solvents were removed *in vacuo* and compound was collected as a white powder (0.011 g, 0.057 mmol, 18% yield).

$^1$H NMR (499 MHz, 90% $CDCl_3$/10%$CD_3OD$) δ 7.75 (dd, *J* = 8.3, 1.5 Hz, 1H), 7.16 (ddd, *J* = 8.6, 6.9, 1.5 Hz, 1H), 7.03 (d, *J* = 8.4 Hz, 1H), 6.89 (t, *J* = 7.6 Hz, 1H), 2.11 (s, 3H). HRMS (MALDI-qQTOF; positive mode; 50:50 CHCA:DHB matrix) Calculated exact mass for $C_{10}H_9ClNO$ [$(M+H)^+$] 194.0373. Observed 194.0374 [$(M+H)^+$].

**2-methyl-3-bromo-4(1H)-quinolone (Br-MHQ) and 2-methyl-3,6-dibromo-4(1H)-quinolone ($Br_2$-MHQ)**

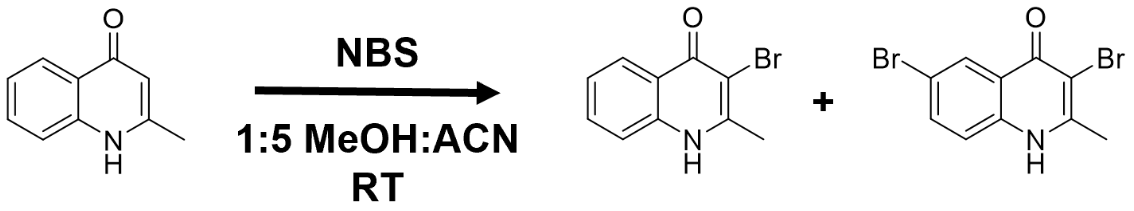


Method was adapted from previously established protocol.[29] A solution of MHQ (0.052 g, 0.33 mmol, 1.0 equiv) in acetonitrile (10 mL) was cooled to 0 ºC. Recrystallized *N*-bromosuccinimide (0.064 g, 0.36 mmol, 1.1 equiv) was added all at once. Methanol (2 mL) was added to reaction to maintain solubility of MHQ and NBS. Reaction was warmed to room temperature and stirred for 9 hours. The reaction was concentrated *in vacuo* and resuspended in dichloromethane (25 mL) and water (25 mL). The layers were separated, and the aqueous layer was further washed with DCM (3 x 50 mL). The pooled organic layer was then washed with brine, dried over magnesium sulfate, and concentrated *in vacuo*. The crude reaction product was purified through a short silica flash column using a 9:1 chloroform: methanol elution system. Reaction product was further purified with semi-preparative RP-HPLC (Phenomenex Luna C8 (2), 10.0 x 250 mm) at a flow rate of 3.0 mL/min with a 10-100% solvent B gradient (solvent A: 0.1% formic acid; solvent B: acetonitrile + 0.1 % formic acid) over 30 minutes. Two major peaks eluted at 11.4 minutes and 14.6 minutes, corresponding to a singly-brominated species and a doubly-brominated species, respectively. Solvents were removed *in vacuo* and singly-brominated species (0.009 g, 0.038 mmol, 12% yield) and doubly-brominated species (0.007 g, 0.034 mmol, 11% yield) were collected as white powders.

Brominated Species

$^1H$ NMR (499 MHz, 90% $CDCl_3$/10%$CD_3OD$) δ 8.09 (d, *J* = 8.2 Hz, 1H), 7.44 (ddd, *J* = 8.4, 6.9, 1.5 Hz, 1H), 7.32 (d, *J* = 8.4 Hz, 1H), 7.23 – 7.16 (m, 1H), 2.46 (s, 3H). HRMS (MALDI-qQTOF; positive mode; 50:50 CHCA:DHB matrix) Calculated exact mass for $C_{10}H_9BrNO$ [$(M+H)^+$] 237.9862. Observed 237.9867 [$(M+H)^+$].

Di-brominated species

$^1H$ NMR (499 MHz, 90% $CDCl_3$/10%$CD_3OD$) δ 8.09 (s, *J* = 8.27 Hz, 1H), 7.55 (d, *J* = 8.4 Hz, 1H), 7.25 (d, 1H), 2.49 (s, 3H). HRMS (MALDI-qQTOF; positive mode; 50:50 CHCA:DHB matrix) Calculated exact mass for $C_{10}H_8Br_2NO$ [$(M+H)^+$] 315.8973. Observed 315.8970 [$(M+H)^+$].

**2-methyl-3-iodo-4(1H)-quinolone (I-MHQ)**

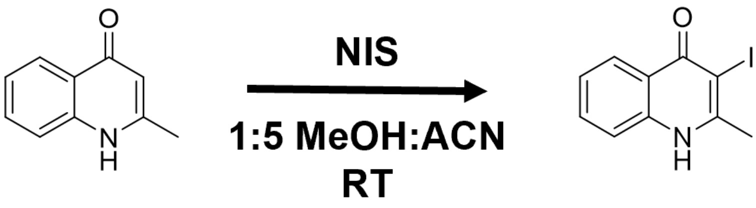


Method was adapted from previously established protocol.[29] A solution of MHQ (0.050 g, 0.31 mmol, 1.0 equiv) in acetonitrile (10 mL) was cooled to 0 ºC. Recrystallized *N*-iodosuccinimide (0.084 g, 0.37 mmol, 1.2 equiv) was added all at once. Methanol (2 mL) was added to reaction to maintain solubility of MHQ and NIS. The reaction was warmed to room temperature and stirred for 20 hours. The reaction was concentrated *in vacuo* and resuspended in dichloromethane (25 mL) and water (25 mL). The layers were separated, and the aqueous layer was further washed with DCM (3 x 50 mL). The pooled organic layer was then washed with brine, dried over magnesium sulfate, and concentrated *in vacuo*. Reaction product was purified with semi-preparative RP-HPLC (Phenomenex Luna C8 (2), 10.0 x 250 mm) at a flow rate of 3.0 mL/min with a 10-100% solvent B gradient (solvent A: 0.1% formic acid; solvent B: acetonitrile + 0.1 % formic acid) over 30 minutes. One major peak was collected at 13.2 minutes, corresponding to a singly-iodinated species. Solvents were removed *in vacuo* and compound was collected as a white powder (0.022 g, 0.111 mmol, 36% yield).

$^1$H NMR (499 MHz, 90% $CDCl_3$/10%$CD_3OD$) δ 8.27 (d, *J* = 8.4 Hz, 1H), 7.86 (d, *J* = 8.6 Hz, 1H), 7.69 (t, *J* = 7.6 Hz, 1H), 7.46 (t, *J* = 7.0 Hz, 1H), 2.85 (s, 3H). HRMS (MALDI-qQTOF; positive mode; 50:50 CHCA:DHB matrix) Calculated exact mass for $C_{10}H_9INO$ [$(M+H)^+$] 285.9729. Observed 285.9722 [$(M+H)^+$].

#### Supplementary Tables

##### Table S1 – All putative alkyl quinolone modifying VHPOs and their source organisms

| Accession Number | Source organism | NCBI Taxonomy ID | Organism Class | Organism Order | 16S rRNA accession | Organism geographic location | Organism Isolation source |
|---|---|---|---|---|---|---|---|
| MBM3759267.1 | Acidobacteria bacterium | 1978231 | Unclassified | Unclassified | Metagenomic | Lake Tanganyika, Tanzania | lake metagenome |
| MCC6989002.1 | Acidobacteria bacterium | 1978231 | Unclassified | Unclassified | Metagenomic | Sjolunda, Sweden | Waterwater metagenome |
| WP_086933921.1 | Agarilytica rhodophyticola | 1737490 | Gammaproteobacteria | Cellvibrionales | NR_158125.1 | Hainan, China | *Gracilaria blodgettii* associated |
| WP_246970656.1 | Alcanivorax sp. S6407 | 2926424 | Gammaproteobacteria | Oceanospirillales | NZ_JALKBH010000020.1 (Locus tag: MWU49_RS17655) | Weihai, China | Marine sediment |
| WP_162906947.1 | Allorhizocola rhizosphaerae | 1872709 | Actinomycetia | Micromonosporales | KX128909.1 | Xinjiang, China | Calligonum mongolicum rhizosphere |
| MCE2471135.1 | Anaerolineae bacterium | 2052143 | Anaerolineae | Unclassified | Metagenomic | Achziv, Israel | Petrosia ficiformis metagenome |
| MCE2473632.1 | Anaerolineae bacterium | 2052143 | Anaerolineae | Unclassified | Metagenomic | Achziv, Israel | Petrosia ficiformis metagenome |
| WP_130969110.1 | Aquabacterium lacunae | 2528630 | Betaproteobacteria | Burkholderiales | NR_174244.1 | Chiayi, Taiwan | Pond water |
| MBM4778199.1 | Archangiaceae bacterium | 2684409 | Deltaproteobacteria | Myxococcales | Metagenomic | Fuzhou, China | Activated sludge metagenome |
| MBL8938882.1 | Archangium sp. | 1872627 | Deltaproteobacteria | Myxococcales | Metagenomic | Nanjing, China | Bioreactor sludge metagenome |
| MCR9160379.1 | bacterium | 1869227 | Unclassified | Unclassified | Metagenomic | N/A | Alexandrium tamarense phycosphere metagenome |

| | | | | | | | |
|---|---|---|---|---|---|---|---|
| MBE0658962.1 | Bryobacteraceae bacterium | 2212468 | Acidobacteriia | Byrobacterales | Metagenomic | Bothnian Sea | Marine sediment |
| MBX2809551.1 | Cellvibrionaceae bacterium | 2026723 | Gammaproteobacteria | Cellvibrionales | Metagenomic | Santa Pola, Spain | Neptune Grass metagenome |
| MXV92527.1 | Chloroflexi bacterium | 2026724 | Unclassified | Unclassified | Metagenomic | Great Barrier Reef | Ircinia ramosa sponge metagenome |
| MXX84763.1 | Chloroflexi bacterium | 2026724 | Unclassified | Unclassified | Metagenomic | Great Barrier Reef | Ircinia ramosa sponge metagenome |
| MYD09778.1 | Chloroflexi bacterium | 2026724 | Unclassified | Unclassified | Metagenomic | Great Barrier Reef | Ircinia ramosa sponge metagenome |
| MYD09780.1 | Chloroflexi bacterium | 2026724 | Unclassified | Unclassified | Metagenomic | Great Barrier Reef | Ircinia ramosa sponge metagenome |
| MYD09944.1 | Chloroflexi bacterium | 2026724 | Unclassified | Unclassified | Metagenomic | Great Barrier Reef | Ircinia ramosa sponge metagenome |
| MYD38911.1 | Chloroflexi bacterium | 2026724 | Unclassified | Unclassified | Metagenomic | Great Barrier Reef | Ircinia ramosa sponge metagenome |
| MBA2454044.1 | Chloroflexia bacterium | 2448782 | Chloroflexia | Unclassified | Metagenomic | Mackay Glacier, Antartica | Soil metagenome |
| MBA2454144.1 | Chloroflexia bacterium | 2448782 | Chloroflexia | Unclassified | Metagenomic | Mackay Glacier, Antartica | Soil metagenome |
| MBA2758935.1 | Chloroflexia bacterium | 2448782 | Chloroflexia | Unclassified | Metagenomic | Mackay Glacier, Antartica | Soil metagenome |
| WP_052553269.1 | Enhygromyxa salina | 215803 | Deltaproteobacteria | Myxococcales | NR_024807.1 | Hokkaido, Japan | Wet black mud |

| | | | | | | | |
|---|---|---|---|---|---|---|---|
| MBC8211257.1 | Gammaproteobacteria bacterium | 1913989 | Gammaproteobacteria | Unclassified | Metagenomic | Black Sea western gyre | suspended particulate matter metagenome |
| MBL4851085.1 | Gammaproteobacteria bacterium | 1913989 | Gammaproteobacteria | Unclassified | Metagenomic | North Pond, Atlantic ocean | marine sediment metagenome |
| MBR9909232.1 | Gammaproteobacteria bacterium | 1913989 | Gammaproteobacteria | Unclassified | Metagenomic | North Pond, Atlantic ocean | marine sediment metagenome |
| MBU2115646.1 | Gammaproteobacteria bacterium | 1913989 | Gammaproteobacteria | Unclassified | Metagenomic | Sweden | Groundwater metagenome |
| MCB1723870.1 | Gammaproteobacteria bacterium | 1913989 | Gammaproteobacteria | Unclassified | Metagenomic | Hong Kong, China | Activated sludge metagenome |
| MCB1800213.1 | Gammaproteobacteria bacterium | 1913989 | Gammaproteobacteria | Unclassified | Metagenomic | Hong Kong, China | Activated sludge metagenome |
| NCC26771.1 | Gammaproteobacteria bacterium | 1913989 | Gammaproteobacteria | Unclassified | Metagenomic | Santa Fe, Argentina | Waterwater metagenome |
| WP_157898942.1 | Luteitalea pratensis | 1855912 | Vicinamibacteria | Vicinamibacterales | NR_156918.1 | Thuringia, Germany | Grassland soil |
| WP_236985789.1 | Marinagarivorans sp. GE09 | 2721545 | Gammaproteobacteria | Cellvibrionales | AP023086.1 (Locus tag: MARGE09_R01) | Kagoshima, Japan | Marine sediment |
| WP_137437665.1 | Marinobacter sp. PJ-38 | 2576384 | Gammaproteobacteria | Pseudomonadales | NZ_SZYH01000004.1 (Locus tag: FDP08_RS20220) | Panjin, China | Red beach Tidal Flat |
| WP_188862589.1 | Marinobacterium nitratireducens | 518897 | Gammaproteobacteria | Oceanospirillales | NR_044528.1 | East China Sea | Marine sediment |
| WP_139559104.1 | Methylotetracoccus oryzae | 1919059 | Gammaproteobacteria | Methylococcales | NZ_SRSH01000014.1 (Locus tag: BWR02_RS14795) | Cixi, China | Paddy Field Soil |
| WP_231757960.1 | Microbulbifer elongatus | 86173 | Gammaproteobacteria | Cellvibrionales | NR_025246.1 | South Korea | Seawater |
| WP_183457526.1 | Microbulbifer rhizosphaerae | 1562603 | Gammaproteobacteria | Cellvibrionales | NR_148866.1 | Lebrija marshes, Spain | *Arthrocnemum macrostachyum* rhizosphere |

| | | | | | | | |
|---|---|---|---|---|---|---|---|
| WP_108732878.1 | Microbulbifer sp. A4B17 | 359370 | Gammaproteobacteria | Cellvibrionales | AB243106.1 | Palau | Ascidian associated |
| WP_156035230.1 | Microbulbifer sp. HZ11 | 1453501 | Gammaproteobacteria | Cellvibrionales | NZ_JELR01000004.1 (Locus tag: AU11_RS15845) | East China Sea | Seawater |
| WP_152453784.1 | Microbulbifer sp. THAF38 | 2587856 | Gammaproteobacteria | Cellvibrionales | CP045369.1 (Locus tag: FIU95_02215) | Giessen, Germany | Marine aquarium containing coral |
| WP_040629207.1 | Microbulbifer variabilis | 266805 | Gammaproteobacteria | Cellvibrionales | NR_041021.1 | Palau | Algae/Coral associated |
| TPV96993.1 | Myxococcales bacterium FL481 | 1979545 | Deltaproteobacteria | Myxococcales | Metagenomic | Layton, Florida | Spongia barbara metagenome |
| MBE2251792.1 | Myxococcus sp. | 1929279 | Deltaproteobacteria | Myxococcales | Metagenomic | USA | Waterwater metagenome |
| MAE34203.1 | Oceanospirillaceae bacterium | 1899355 | Gammaproteobacteria | Oceanospirillales | Metagenomic | North Pacific Ocean | marine water sample metagenome |
| WP_076514445.1 | Oleibacter marinus | 484498 | Gammaproteobacteria | Oceanospirillales | NR_112787.1 | Pari Island, Indonesia | Seawater |
| TNC83410.1 | Oleibacter sp. | 2024845 | Gammaproteobacteria | Oceanospirillales | Metagenomic | Mariana Trench | hadal water metagenome |
| KZZ11471.1 | Oleibacter sp. HI0075 | 1822250 | Gammaproteobacteria | Oceanospirillales | LWFP01001472.1 (Locus tag: A3746_34715) | Hawaii, USA | Seawater |
| MBM3996742.1 | Planctomycetes bacterium | 2026780 | Unclassified | Unclassified | Metagenomic | Lake Tanganyika, Tanzania | lake metagenome |
| WP_171626078.1 | Pseudoalteromonas caenipelagi | 2726988 | Gammaproteobacteria | Alteromonadales | MT348162.1 | Jebu Island, South Korea | Tidal Flat sediment |
| WP_063362279.1 | Pseudoalteromonas luteoviolacea | 43657 | Gammaproteobacteria | Alteromonadales | NR_026221.1 | San Diego, California | Seawater |
| KZN74784.1 | Pseudoalteromonas luteoviolacea H33-S | 1365252 | Gammaproteobacteria | Alteromonadales | NZ_AUXY01000116.1 (Locus tag: N477_RS22360) | Tasman Sea | Seawater |
| WP_192540197.1 | Pseudoalteromonas prydzensis | 182141 | Gammaproteobacteria | Alteromonadales | NR_044803.1 | Antartica | Sea Ice |

| | | | | | | | |
|---|---|---|---|---|---|---|---|
| WP_141240123.1 | Pseudoalteromonas sp. HM-SA03 | 2029678 | Gammaproteobacteria | Alteromonadales | NZ_NSDG01000197.1 (Locus tag: CKO50_23305) | Moreton Bay, Australia | Blue-ringed octopus salivary gland |
| WP_166165103.1 | Pseudomaricurvus alcaniphilus | 1166482 | Gammaproteobacteria | Cellvibrionales | NR_137412.1 | South Korea | Tidal Flat sediment |
| MCG8311860.1 | Pseudomonadales bacterium | 1891229 | Gammaproteobacteria | Pseudomonadales | Metagenomic | Abo Shoosha reef, Saudi Arabia | Coral reef metagenome |
| WP_128198684.1 | Rubrivivax albus | 2499835 | Betaproteobacteria | Burkholderiales | OP597510.1 | Hunei, Taiwan | Fish pond |
| WP_230564822.1 | Saccharothrix Luteola | 2893018 | Actinomycetia | Pseudonocardiales | OL378195.1 | Chifeng, China | Soil |
| NUS64381.1 | Saccharothrix sp. | 1873460 | Actinomycetia | Pseudonocardiales | Metagenomic | Hawaii, USA | Soil metagenome |
| MCH1931814.1 | Shewanella shenzhenensis | 2914042 | Gammaproteobacteria | Alteromonadales | MZ477861.1 | Shenzen, China | Mangrove sediment |
| WP_237443824.1 | Sinobacterium norvegicum | 1641715 | Gammaproteobacteria | Cellvibrionales | NR_144601.1 | Valencia, Spain | Pecten maximus gonads |
| TVR74215.1 | Sphaerobacteraceae bacterium | 2099683 | Thermomicrobia | Sphaerobacterales | Metagenomic | Cariboo Plateau, Canada | Alkaline lake metagenome |
| WP_132875434.1 | Tamaricihabitans halophyticus | 1262583 | Actinomycetia | Pseudonocardiales | NR_145901.1 | Jiangsu, China | Tamarix chinesis rhizosphere |
| WP_093031253.1 | Thiocapsa roseopersicina | 1058 | Gammaproteobacteria | Chromatiales | NR_041729.1 | Global | waste washwater pond |
| WP_200249935.1 | Thiococcus pfennigii | 1057 | Gammaproteobacteria | Chromatiales | NR_036977.1 | Massachusetts, USA | Salt marsh |
| WP_207188716.1 | Thiocystis minor | 61597 | Gammaproteobacteria | Chromatiales | NR_036976.1 | Reyershausen, Germany | Lake |
| MCF7989346.1 | Thiohalocapsa sp. | 2497641 | Gammaproteobacteria | Chromatiales | Metagenomic | Laguna del Tobar | lake water metagenome |
| WP_017029669.1 | Vibrio breoganii 1C10 | 1205908 | Gammaproteobacteria | Vibrionales | NZ_AKXW02000127.1 (Locus tag: B003_RS13070) | Massachusetts, USA | Brown seaweed |

| | | | | | | | |
|---|---|---|---|---|---|---|---|
| OED83209.1 | Vibrio breoganii ZF-55 | 1188253 | Gammaproteobacteria | Vibrionales | NZ_AJYL02000008.1 (Locus tag: A1QE_RS01130) | Massachusetts, USA | Filtered Seawater |
| WP_155759790.1 | Vibrio natriegens CCUG16374 | 1219067 | Gammaproteobacteria | Vibrionales | NR_026124.1 | Hawaii, USA | Enriched Seawater |
| WP_118119916.1 | Vibrio sp. dhg | 2163016 | Gammaproteobacteria | Vibrionales | NZ_CP028944.1 (Locus tag: DBX26_RS20645) | Korea | Seaweed sludge |
| WP_014234363.1 | Vibrio sp. EJY3 | 111637 | Gammaproteobacteria | Vibrionales | NC_016613.1 (Locus tag: VEJY3_RS00150) | Incheon, South Korea | Decomposed grapsid crab |
| WP_237314952.1 | Vibrio sp. J1-1 | 2912251 | Gammaproteobacteria | Vibrionales | NZ_JAKKCM010000051.1 (Locus tag: L3V31_RS21990) | China | Marine sediment |
| WP_248006878.1 | Vibrio sp. ZSDE26 | 2847292 | Gammaproteobacteria | Vibrionales | NZ_JAJHVV010000030.1 (Locus tag: KP803_RS21720) | Qingdao, China | seawater |
| MBR9787475.1 | Vibrionaceae bacterium | 1889773 | Gammaproteobacteria | Vibrionales | Metagenomic | East China Sea | marine sediment metagenome |
| MBR9875570.1 | Vibrionaceae bacterium | 1889773 | Gammaproteobacteria | Vibrionales | Metagenomic | East China Sea | marine sediment metagenome |

**Table S2 – Biosynthetic potential of alkyl quinolone VHPO containing strains**

| Accession Number | Source organism | NCBI Taxonomy ID | Genomic Assembly availability | Presence of PQS BGC | Number predicted secondary BGCs in organisms | Putative BGC context of VHPO |
|---|---|---|---|---|---|---|
| WP_086933921.1 | Agarilytica rhodophyticola | 1737490 | ASM215722v2 | No | 13 | No Cluster |
| WP_246970656.1 | Alcanivorax sp. S6407 | 2926424 | ASM2301611v1 | No | 2 | No Cluster |
| WP_162906947.1 | Allorhizocola rhizosphaerae | 1872709 | ASM342677v1 | No | 34 | No Cluster |
| WP_130969110.1 | Aquabacterium lacunae | 2528630 | ASM431086v1 | No | 3 | No Cluster |
| WP_052553269.1 | Enhygromyxa salina | 215803 | ASM299463v1 | No | 43 | Nystatin A1 Cluster (27% id) |
| WP_157898942.1 | Luteitalea pratensis | 1855912 | ASM161886v1 | No | 4 | No Cluster |
| WP_236985789.1 | Marinagarivorans sp. GE09 | 2721545 | ASM2165555v1 | No | 11 | No Cluster |
| WP_137437665.1 | Marinobacter sp. PJ-38 | 2576384 | ASM2473361v1 | No | 3 | No Cluster |
| WP_188862589.1 | Marinobacterium nitratireducens | 518897 | ASM1464537v1 | No | 15 | No Cluster |
| WP_139559104.1 | Methylotetracoccus oryzae | 1919059 | ASM617598v1 | No | 7 | No Cluster |
| WP_231757960.1 | Microbulbifer elongatus | 86173 | ASM2255410v1 | No | 5 | No Cluster |
| WP_183457526.1 | Microbulbifer rhizosphaerae | 1562603 | ASM1419172v1 | No | 8 | pentabromopseudilin cluster (20% id) |
| WP_108732878.1 | Microbulbifer sp. A4B17 | 359370 | ASM307627v1 | No | 6 | No cluster |
| WP_156035230.1 | Microbulbifer sp. HZ11 | 1453501 | No Accession provided | Yes (PqsBC) No PqsL/H | 5 | No cluster |

| | | | | | | |
|---|---|---|---|---|---|---|
| WP_152453784.1 | Microbulbifer sp. THAF38 | 2587856 | ASM936353v1 | No | 5 | No cluster |
| WP_040629207.1 | Microbulbifer variabilis | 266805 | ASM38056v1 | No | 5 | No cluster |
| TPV96993.1 | Myxococcales bacterium FL481 | 1979545 | ASM650810v1 | No | 11 | No Cluster |
| WP_076514445.1 | Oleibacter marinus | 484498 | IMG-taxon 2681813515 | No | 5 | No Cluster |
| KZZ11471.1 | Oleibacter sp. HI0075 | 1822250 | ASM163500v1 | No | 4 | No cluster |
| WP_171626078.1 | Pseudoalteromonas caenipelagi | 2726988 | ASM1314085v1 | No | 11 | No cluster |
| WP_063362279.1 | Pseudoalteromonas luteoviolacea | 43657 | ASM670414v1 | No | 25 | No cluster |
| KZN74784.1 | Pseudoalteromonas luteoviolacea H33-S | 1365252 | H33-S_1.0 | No | 19 | No cluster |
| WP_192540197.1 | Pseudoalteromonas prydzensis | 182141 | ASM1492535v1 | No | 6 | No cluster |
| WP_141240123.1 | Pseudoalteromonas sp. HM-SA03 | 2029678 | ASM228934v1 | No | 11 | aranazole cluster (1% id) |
| WP_166165103.1 | Pseudomaricurvus alcaniphilus | 1166482 | ASM1144039v1 | No | 4 | No Cluster |
| WP_128198684.1 | Rubrivivax albus | 2499835 | ASM401651v1 | No | 10 | No Cluster |
| WP_230564822.1 | Saccharothrix Luteola | 2893018 | ASM2085956v1 | No | 43 | No Cluster |
| MCH1931814.1 | Shewanella shenzhenensis | 2914042 | ASM2232155v1 | No | 3 | No Cluster |
| WP_237443824.1 | Sinobacterium norvegicum | 1641715 | Sinobacterium_norvegicum_CECT_8267_Spades_Prokka | No | 5 | No Cluster |
| WP_132875434.1 | Tamaricihabitans halophyticus | 1262583 | ASM434087v1 | No | 11 | No cluster |
| WP_093031253.1 | Thiocapsa roseopersicina | 1058 | IMG-taxon 2684622843 annotated assembly | No | 9 | Unassigned betalactone |
| WP_200249935.1 | Thiococcus pfennigii | 1057 | ASM1658383v1 | No | 7 | No Cluster |
| WP_207188716.1 | Thiocystis minor | 61597 | ASM1665346v1 | No | 9 | No Cluster |

| | | | | | | |
|---|---|---|---|---|---|---|
| WP_017029669.1 | Vibrio breoganii 1C10 | 1205908 | ASM28088v2 | No | 2 | No Cluster |
| OED83209.1 | Vibrio breoganii ZF-55 | 1188253 | ASM28697v2 | No | 3 | No Cluster |
| WP_155759790.1 | Vibrio natriegens CCUG16374 | 1219067 | ASM168008v1 | No | 7 | Nearby Alkylpyrone cluster |
| WP_118119916.1 | Vibrio sp. dhg | 2163016 | ASM344377v1 | No | 6 | Nearby Alkylpyrone cluster |
| WP_014234363.1 | Vibrio sp. EJY3 | 111637 | ASM24138v1 | No | 6 | Nearby Alkylpyrone cluster |
| WP_237314952.1 | Vibrio sp. J1-1 | 2912251 | ASM2172839v1 | No | 5 | Nearby Alkylpyrone cluster |
| WP_248006878.1 | Vibrio sp. ZSDE26 | 2847292 | ASM2314569v1 | No | 5 | No Cluster |

**Table S3 – *E. coli* codon optimized DNA sequences for selected alkyl quinolone VHPOs**

| Accession no. | Organism | Abbreviation | Sequence |
|---|---|---|---|
| WP_052553269.1 | *Enhygromyxa salina* | esVHPO | ATGCAGTCTACGCGCTTTAGCACCTCATCGTCATTATTACTTCTTGGGGCGCTGGCTACA GCCGCGTGCGATCCGACGCTTGATCCTGCCGCGACGCGCGTAGCAGCCTCAGACGAG GGAGTTAACGACCAGGAGTGCACTACCCTGTTACCCGCAAACGTGCCGGGTCTGCAG GCGCAGCTGCCGACACAGACCATGGGTGCCGACTTCGACTTTGACACCGGCAACGCT CCGATCGAGATCGTTATCCCTGCCGTATTGCCGGTCATCGCTGGCAGCGTGGCACCGG GCGACGCAACGATCGTACTTCGCTTCACCACGATGCTGAGCAACGCGTGGTTTGACGC GACCGCGCCCTACCACCCCACTGCCGTGGGTGTGTACTCCAATTTAGGCCGCCGCCCT GCCAGCGAGAGCACCACCCACGCGAACATGAACATCGCGATCCTCTACGCCAGCTACC GTACGCTGAACAGCCTGGCTCCACAGCACGCGGCGGACTGGGACGCACTGATGGTGA GTCTGGGTCTTGATCCTCACGACGACCACGAGTCAACCACCGATCCTATCGGAATCGG AAACGCTGCGGCTGCGGCTCTGTTGGCTGTTCGCGAGAATGACGGCTTCAACCAGTTA GGTTTCGAAGGCGGCCGCGAGTACAACCCCATCCCATACGCGGACTACACGGGCTAC GAACCTCGCAACACACGCTTCGAGATCAAGGACGAGCGCCGCTGGCAACCGGCCATT GTGACGAGCCGCTACGGCATCACCCGCGCCCAACACTTCGTAACGCCGCAGTATGCCC TTACTCTGCCGTACAGTTACGACGACCCTCAGGACTTTGGCGTGCCTTTACCGGACAA GTCATTAAAGAAGGGTAGTCACGCCAAGAAGAAATACCGTGCTCAGGCGGACGAGGTG CTTGAGGTCTCTGCGAACCTGACGGACGAGCAAAAGGTGACTGCTGAGCTGTTTGAG GACAAGATCCGCAGTCTGGGTTTCTCAGCATTATTTGTGAGTCTGTCAAGCGGTCACAG CTTACTGGACTTCGTACACTACGACTTCTTGACCAACTTGGCCGCGTTCGACGTTGGCA TCGTTGTCTGGCAGGAGAAAACGCAGTACGACGCCGTGCGTCCTTTCACCGCGATCCG CCATATCTATGGCGACGACGAGATCACAGCGTGGGGCGGACCCGGCCAGGGCACCGT GAACGACCTGCCAGCGAACGAATGGCGCTCATACCTCGACGTAGCTGACCACCCAGA GTACCCGTCGGCATCGGCAGCGTTTTGCGCAGCGCATGCCCAGGCAAGTCGCCTTTTC CTTGGCACCGACGATTTAGGCTGGACGGTTCCGATCCCGGCCGGTAGCTCGATCGTCG AGCCCGCCATCACTCCGGCAGCCGACCTGAACCTGCACTTTCCGACCTTCACCGACTT CGCCACTCGCTGCGGATACAGCCGTCTTTGGGGCGGTGTACACTTCGAAGACGCCATC TTAGCTAGTTTCGAACTCGGCGACGAGATCGGGGCCGGCGCCTACGAGTTCGTGCAA GCGCACATCGACGGCACTCCACCCTGA |
| WP_157898942.1 | *Luteitalea pratensis* | lpVHPO | ATGTGCAGAGTTATCCCGCGTATTGCCTTTTGTACCGCCGCCCTGGTTTTAATGCCAGC ACGTCCCCAAGCCCAAGCGCCGCCTTTTGATTTTAGCTCAGGTAATATTGGGATTGAAG TAATAATTCCGGCGGTTATCCCAGCTCTGTTTCAAACAACAAGCCCTAATGATGCCCCAA TAATTTTACGCCATACAACCCTGATTACAAACGCCTGGTTTGATGCTATTGCGCCGTATCA CCCTACAGCAAAAGGTGTGTATACACGCTTGGAAAACCGTCCACCAGCTGAAGCGACG ACTCGCAATAAGAATATAGCAATGGCTTATGCCACCTATCGACTTTTGAATAGATTAATGC CACGCTTTGCCGCTGATTGGCGCCAAATGTTAGTTTCAGTGGGTCTGGACCCCGATGA CGCCTCGGTGGACGTACGTACCCCCATAGGGATTGGCAATGTGGCAGGTTCCGCAGTT GCGGTCGCTCGTGAACGCGATGGGATGAATCAACTGGGCGATGAAGGTGGACGTGTT |

| | | | |
|---|---|---|---|
| | | | TATGACTTACGTCCATACGCTGATTATACAGGTTATATGCCAGTGAATACTCCATGGAAATTGTCCAACCCCTCTAGATGGCAACCTCTTTTCACTACGCCAGGAAATGGAACATTCCTTGGCCAACAATTTGCGACGCCCCAGTGGGGAGCCACAACTCCATATTCATATGGGAACCCCGCTGCATTTGGTACACCGCCACCAATAGATTCTAATCACCATCGTCGTCAAGCATACGAAGCTCAGGCAGATGAAGTAATGGATGCACAAGCCACCCTTACTGACTATCAGAAAATGGTTGCTGAACTGTTTGATAATAAAATAACCGGTCTTGGGTTTGCCGCACTGTTTATAGCTCAAAGCCGTAATATGTCCCTGGATGAATTTGTCCATTATGACTTTCTGACAAATGTTGCAGCTTTTGATGGTGGCATTACCACCTGGCGTGATAAATACTTATATGATGCGGTTCGCCCGGTTACGGCTATCCGTTATTTGAACAGAGGCCGCACCATTACGGGTTGGGCTGGTCCGGGTAGAGGTATTGTTAATGATCTCCCTGCTAATGAATGGCGGAGCTATCTCAATACTGCTAATCATCCAGAATACCCAAGTGGGTCGAGCTGCTTTTGTGCTGCTCATGCACAAGCTAGTCGCCGTTACTTAGGGAGTGACCAATTCGGATGGAGCGTTCCTAAACCAGCGGGTTCTAGCGTCATAGAACCGGGTGTAACTCCAGCGGCGGATGTGGTTCTCGGCCCCTGGCAAACTTTTACAGAATTTGAAGAAGAGTGTGGAATGTCCCGACTGTGGGGTGGTGTCCACTTTCGGCCGGCTATTGAAGAAGCCCGCAATTCCTGCCGCCAAGTTGGTGATAAAGCATTTGAATTTCTTCAACGTAAACTCGCAGGTCAATGA |
| WP_139559104.1 | *Methylotetracoccus oryzae* | moVHPO | ATGAATCCGCTCTCTCGTCGTTCTTTCGCGACATTCTGGTTCACGTTAGCCTCGGTTGTATTCGCTGTAGGGGCTGAATCCGCTCAAGCACAAACATTTGACTTTGATAACGGATCAGCACCGGTCGAAGTTATTATTCCGGCCATAATACCAATTGTACTGAATGATGTAAGTGCTGGCGCCAATGACGCCACCCTCGTGTTGCGCATCACGTCGGAAATTGCGGTTGGTTGGTTTGATGCTATCGCGCCATACCATCCGACAGCACGAGGCATTTACAGTAATTTGCCTCGCCGTCCACCGAATGAATCTCAGAGTAATTCGAACAAGAACATTGCCATGTTATACGCTAGCCGTAGAATACTTATGTCATTACTGCCTGATAGAGCATCGGATTGGGATAATCTTCTCACGTCCGTCGGCCTGGACCCTAATAATGGAAGTCAAAACCTTGCGAGCCCCGTAGGCATCGGCAATGTAGCGGGTAACGCTGTGGTCGCGTTTAGAGAGCATGATGGTATGAATCAGTTGGGAGATGAGGGCGGCCGAACATTCAACAGATTCCCATATATGGACTCAACGGGTTTTAAACCTGTTAATACGGCTTATGAAATACTGGACCCAACACGCTGGCAACCCTTAGTTGTGACACAAGGGAATGGAATTTTCCGAGTCCAACAGTATGTGACGCCTCAACTTGCACGGACGCGTCCGTATGCTATAGATCCCCGTAACTTTTCCTCACCAGAACCTATTGCTAGCCTTCTTAAGCGCGGCAAGAACGGACGTGATGCGTATAAACAACAGGTAGATGAGGTACTGGCCATCAGTGCCAGCCTGAATGACTATACGAAAGCCCAAGCCGAATTCTTTGATAATAAGTTTCGTTCAATCGGCTTTGTAGCCCTCTTTGTAGCAGGGGTAAATCAGTTTTCGCTGGATGAATTGGTACAATATGATGCAATGCTGAATATGGCTGCCTTCGATGGGGCTATTGTCAGCTGGAAACAGAAAACCAGATGGAATGCCGTAAGACCCATGACCGCCGTTAGACACGTGTATGGGGACCGTACAATCACCGCATGGGGCGGACCAGGGAAGGGTACACAACGACTCCCGGCCAATGAGTGGCAGTCTTATTTACAAACCCCGGATCATCCCGAATACCCGAGCGGCTCTAGTTGTTTCTGTGCAGCTGAAGCACAGGCGACACGCCGTCTGCTGGGGTCGGATGAGTTGGGATGGCAGGTGCTGGTTCCAAAGGGTCAGAGTCGTATTGAACCTGGGATAACTCCATCCCAAGATACGACTTTAGTCTTTAATACCTGGACTGATTTTGAAAATCAATGCGCACAAGCCCGAGTTTTAGGTGGCGTACATTTTCAAGCTTCCGTGGCAGCGAGCATAGCTATGTGC |

| | | | |
|---|---|---|---|
| | | | CGGCCCGCAGGGGATGCCGCATATGAATTCATACAACGTCACGTTAATGGTCTCCCTTAA |
| WP_183457526.1 | *Microbulbifer rhizosphereae* | mrVHPO | ATGGGGAAACTGTTCCATAAGGGAGTTAGTAAAGTCCTGTGCTGTACGATGGTAATAGTCATCGGGTGTTTTACTAAGTCTTTCGCATCCGAGTCGGAAACATCGTACGATTTCAAGAACGGAAACGCAGTTATTGAGGTAGCTGTGCCTGCCCTGCTCCCCGTAATCTTGAGTGACGTGTCCTCTACAGCTGGAGACGCGACACTTATATTGCGTATGTCAACTCTTATGAGCAACGCATGGTTTGACGCGTCAGCCCCCTACCACCCAACCGCTGTAGGCGTATACAGTAGACTGGGTCGGCGTCCGCTGAGTGAGTCCGAGACTAACACTAACATTAACATCGCCATCTTATACGCGAGCTACAGAGTTCTTTCATCGTTACTTCCGCAACGTACCCCAGTGTGGAGAAAGATGCTGACAGACATCGGACTGGACCCAGACGACGATTCAGTCGATTTGGCTACGCCAGTCGGTATCGGAAACGCTGCAGGATTAGGGGTAGTGCGAGGTCGGGCGCGTGACGGAATGAATCAACTCGGGGATGAATCTGAAAGAGCTTTTAACCCCATGCCGTATATGGACTACACTAATTACCGTCCTATTAACAGTGCGTATCGCTTATTTGATCCAAGCCGTTGGCAGCCGGACTTGCAACGTAAAGGTATGGGATTATATAAGATTCAACAATTCGTTACTCCGCAATACGCACTCACGGAACCGTACTCATATCAAGCTCCGCAACGGTTTAACATGCCCCCACCATACGCAAGTAACCATTGGAACTACCCTGCATACAAACAACAAGCAGACGAGGTCTTAGCCGCGTCAGCCGATTTAAATGATGAACGAAAACTGAAGGCTGAACTGTTCGATAACAAGATCGAGTCGCTCGGTTTTAGCGCCCTCTCAACAGCAATTAGCCGCGGCCTGAGTTTACAAGAGACTATTGAATTAGACTTCTTAATTTCGATGGCAGCGCATGATGCTGGGATAGTGATTTGGCAAGAGAAAAGACGTTACGATGCAGTTCGCCCGTTTTCCGCGATCCGATATTTATACGGTGACGAGTGGGTCTCTGCTTGGGGTGGTCCAGGTCGTGGAACCATCCGCTTACCAGCGAATCAATGGAAAAGTTACTTAGAAGAGGCCGACCACCCAGAATACCCTTCTGCCTCTACCTGCTTTTGCGCGGCCCACGCTCAAGCCGCCCGTCGCTATCTGGGTACCGACGTTCTCAACTGGGAGTTCAGTAAGGCTGCAGGCTCATCACACGTAGAACCTGGAATTACACCCGCAGAGGATACGATTCTGCGGTTCGATACTTGGACAGACTTCGTTTCTGACTGTGGGCAATCTCGCGTCTGGGCCGGCGTTCACTTCCAAGCGGCCGTTGATGTTTCAACCCAGATGTGCGATGTGTTCGGCGACATGGCACATGAGTATTTAAGCTCACTCATAGACGGTTCCGCTCCAGTTCGGCGCCCCGCGAAAGGAAAGAGATATAAGTGA |

| | | | |
|---|---|---|---|
| WP_108732878.1 | *Microbulbifer* sp. A4B17 | A4B17-VHPO | ATGCTGAAGAATATCATGTATAAGAACGCGTTAAAATCCATTGCCGGCATCCTCTTAGTG<br>AGTGCAAGCTGGCAGGCACAGGCGATCAGTTGCCCCCAGCTGACCGACGTTTCGCTT<br>CCGCCGAAATGTAGCTCCGAAGAATTAGCAACGTGCAACGCAGCCGTCCAGAACGTCA<br>TTCCCGCCGCCGCCTGTGCCATCTTTGAGGTTAGCCCAGCAGCAAATGATGCGACCTT<br>AGTACTTCGTTTCACCACCCTCATCACTAATAGTTGGTTTGATGCAGCTGCACCTTATCA<br>CCCAACGGCCAAGGGTGTTTATAGCGACATCGCCCGCTTAACTGAACCTACGGACAATA<br>CCGAGTTAAACATTGCGCTTAGCTACGCTTCACATAAGGTACTGAGCGGCCTCTTCCCG<br>CATCTGGTATCAGAATGGGACGATATGTTGATCGAGTTAGGCCTGGATCCGGGTAATTTG<br>TCCGAGGATACCACCACCCCGATTGGGATCGGAAACTATGCTGGTCGTAAGGTACTTGA<br>GGGGCGTGCTAATGACGGTATGAATCAACTTGGTAATGAGGGCAACCAGCTGTATCACC<br>GTCTGCCGTATGCGGATTACACCGACTTTAAGCCGGCTAATTCAGCCTACAAATTACGTT<br>TTCCTAGCAAGTGGCAACCGGCAGTCCTGATGGATCGTCCGGGCGTATTTCGCGTCCA<br>ACAGTTTGTCACCCCTCAGATCGCCCTTACCACGCCATACAGCTATGAGTCGCCAGAAG<br>AATTCACTAGCCCGGTGCCTCTCAAATCAAAGATTTGGAATTACCCGTCATATGTGAGTC<br>AAGTTAACGACGTGTTAGCAACCTCGGCCAATCTGACCGACGAGCAGAAAATGAAGGC<br>CGAGCTGTTTGACATGAAGATTTTCAGCCTGGGCTTTGCCGCCGTATTTGCCAGCGAG<br>AGCCAGGGCTTGTCCCTGATGGATTTTATCCACTATGATTTCCTGACCAATATGGCGGCC<br>TTTGACACGGCGATCACCATCTGGAAAGAGAAGCGTCGCCATAATAGTGTACGTCCGTT<br>TTCGGCCGTACGTTATATTTATAAGGATGAGCCCGTTACAGCCTGGGGCGGTGTTGGTA<br>AGGGCACCGTCAACGATTTACCTGCCACGCAGTGGACCTCGTACCTGCCGGTGGCCG<br>ATCATCCGGAGTACCCGTCCGCTAGCGCCTCATTTTGTGCGGCGCACGCAGAAACCAG<br>CCGTTTATTTCTGCCGCAAGGCGACACTTTGGGCTACCTCGTGACAGCGCTCGCGGGT<br>AGCAGTCAAATTGAGCCTGGAATTACGCCGCAGACCGACTTAAACCTTTATTTCCAAC<br>CTGGACGTCATTTGAAGAAGACTGTGGCAATTCGCGTGTGTGGGGCGGAGTACACTTT<br>GAACCGAGCGTGCCGGCCGGCCAAGCAATTGGTCGTCAGATCGCCCATCGCGCATAT<br>GAATTCTATCTTTCTAAACTGGCGGGTAACTGA |
| WP_156035230.1 | *Microbulbifer* sp. HZ11 | HZ11-VHPO | ATGAAAATTGGAATGCTGCTTCGTGCTGTGCTGTGTGGAACCGCTCTTGCATTTTGTGC<br>CCTCAACACACAGGCTCAGCCGACCACCCCGCCACCCGTTGATCTCGATAATGGTAAT<br>GCAGCCATTGAGGCTGTAATCCCGTACGTCGCGCCCGTTACATTTGAGTATGTAAGTGC<br>CACAGGTGGCGACGCTACCTTAGTGCTCCGTATTACGACACAGATCACGAACGCATGG<br>TTCGACGCGTCTGCCCCCTATCACCCGACTGCGGTAGGCGTATATTCTCGCTTAGGTCG<br>CCGCCCGGCAAATGAGTCCGCAAATAATCGCAACATTAACATCGCGTTATTGTTTGCCTC<br>GTATCGTGTGCTGAATACTTTGCTGCCGCTGCAGCATGCCACATGGCGCGCGATGTTG<br>GAAGCCCAGGGTCTGGACCCGGATGACAATTCAACTGATCTGACTTCTCCAATCGGAAT<br>TGGTAACGCTGCGGGCGCCGCAGTCGCTAAAGGTCGCCTGCGCGACGGTATGAATCA<br>AGTTGGCGACGAGACCCGCAGTGGTCCTCTTCGTGCGGCTAACCCGATGCCGTATATG<br>GATTACACCGGCTACCAGCCCATCAACACTGCGTACACTCTGTTCAACCCGAGTCGTTG<br>GCAGCCAGACATCCAACGTAAAGGCATGGGCCTGTATAAAATCCAACAATTCGTCACTC<br>CGCAGTACGCCCTGACGGAGCCGTACTCATACAAGTCACCACGCAAGTATCGCGCACT<br>GCCACCTTTCGCGTCGTTCCATTGGAACTTTGCCGCATATAAACAGCAAGCGGACGAAG |

| | | | |
|---|---|---|---|
| | | | TTCTGGCAGCGTCGGCGAATCTGACCGATGAACAGAAGCTCATGGCCGAGCTGTTTGA<br>TAATAAAATTGAATCATTAGGCTTTTCTGCCGGCTTTGCGGCGTTGTCTCAAGGTTTATC<br>GCTCATGGAATTTATCCACTTTGATTTCCTGACCAATATGGCAGCACATGATGCGGGCAT<br>CTTTGTATGGCAAGAGAAACGTCGCTATGACGCTGTCCGCCCGTTCTCAGCGATTCAAT<br>ATATCTATGGAAAGCAGCCCGTGACGGCGTGGGGCGGTCCGGGACAAGGCACCCAGA<br>GTATCCCGGCAAATACGTGGAAATCTTATCTGGAAGAGGCTGATCACCCTGAATACCCG<br>TCCGCAAGCACGTGCTTCTGTACAGCCCACGCGCAATCGGCACGCCAATACTTAGGCT<br>CCGACGCGTTAAATTGGTCTGTACCGTACACGGCCGGCAGTAGTCGCATCGAGCCCGG<br>AGTTACCCCGGCAAATGATATGACCCTTAAATTTCCGACTTGGACGTCATTAGAGGAGAA<br>TTGTGGACAAAGCCGTGTATGGGCAGGTGTGCATTTCCAGGCCGCGGTTGATGAGGGT<br>CGTCGCATTTGTGGCGTGTTTGGCGACAATGCGTATAACTATTTGCAAACCTTAGTCGCT<br>GGTACCGCGCCTGAACGTGAACCAGCGGATCGTTTGCGCGGCCGCCGCAAGTGA |
| WP_076514445.<br>1 | *Oleibacter marinus* | omVHPO | ATGAAATTGCCCTTGTTGAAAGGTCGTCCGTTGGCCCTGGCCGCCTCGGCACTTCTTT<br>CTATTGCCGTTAATCAGGCGTCGGCGTATGATCTGCAAACCGGGAACGCACCAGTGGA<br>ACTGGTCATTAGCAAGGTGGCGCCGGCTATTTTCCAGGATATTAGCGCCACTGCCGGC<br>GACGCGACTCTGGTACTGCGCGTGACGGCGCAAGTAACAAATGCCTGGTACGACGCAT<br>CGGCGCCTTATCACCCTACGGCGGTAGGGGTCTATTCGCGCATCCCGAACCGCCCCGC<br>ATCCGAAAGTGTTGACAACGAGAATATTAACATCGCGACTCTGTATGCCAGCTACCAGG<br>TGCTGAAAGTGCTTTTACCGCAACGCACGGCCGAATGGCGCGGGATGTTAGTTGAGGC<br>GGGCTTAGATCCTGATGACACCTCAACAGATCTGACAACCCCGGTTGGTATCGGCAATG<br>TTGCCGGGGCAGCCGTGGCCGAAGGCCGTCTGAACGATGGCATGAATCAAGCGGGTT<br>GGATCAACGAAGATGTACATCCGCAACCTTACGCCGATTATACGGGATATGAGCCAAAG<br>AATACCGCGTATGACCTGAAGCGTCCGAGCAAATGGCAGCCGGATGTACAACGTAAAG<br>GCCTGGGCTTGTATAAAGTACAGCAATTTGTAACTCCCCAGTACGCGAATGTCGAGCCG<br>TATAGCTATGACGATCCTGAACGCTTCTCAGTGCCTTGGCCTGCGAATTCGTTCGCGCC<br>CTTCAAACACTTGTACAAGCAGCAAGCGGACGAAGTCTTAGTGGCCAGCGCGAACCTG<br>ACCGAAGAACAGAAGCTGAAAGCGGAACTGTTTGATAACAAAATCTTCAGCCTGGGCTT<br>TAGTGCGGTTTTCGCGGCTCAATCTCGTGGGCTGTCCTTACTGGAGTTCATCCAATTAG<br>ATTTTCTTACGAATATGGCGGCATTCGATGCAGGTATTTTCGTGTGGCAGGAGAAGGCG<br>GAATACGATGGCGTACGCCCTTTCAGTGCTATTCAGCATATTTATGGCGACCAACCGGTT<br>GAGGCCTGGGGCGGACCTGGAGAAGGTACGGTAATGTTGCCAGCCAACCAGTGGCGT<br>GCGTATATGGAAGAGGCGGACCACCCGGAGTACCCAAGCGCTAGCAGCTGCTTCTGTG<br>CCGCCCATAGTCAAAGCGCCCGTGCCTTTCTGGGTGACGATATGCTGGGTTTCCCCGT<br>GGAATATCCGGCAGGCAGCTCTCGTATTGAGCCGGGCCTGACGCCCGCGGCCGATAC<br>TACCCTGGTATTTAATACCTGGAGTGAGTTCGAACAAGACTGTGGACAGTCGCGTGTCT<br>GGGCTGGGGTACACTTCCAAGCGGCTGTGGATGAGTCATTAGCCCTGTGCGGTGACTT<br>CGGTGACATGGCCTATGATTACATGCAATCGCTGATTGATGGTACCGCCCCGGCCCGTG<br>GCCCATCACAGCCACTCCGTGATCCCGCGACATATAGCAATAAGAAATGGTGGTGA |

| | | | |
|---|---|---|---|
| KZN74784.1 | *Pseudoalteromonas luteovioleacea* H33-S | pIVHPO | ATGAACAAGTACTTAAGCAAAATTTCACTCTTGCACCAGATTAGAAAGCTCATGAAATTG<br>CGTAACATTAGTGTTTTCCTGCTGTCGTTAATATGCGTAACTCACGTGCAGGCTAACCAC<br>CTGGACGAATTCGATTTCGACAAGGATAACGGTGCCCTTGACGTTGTAGGCTGGGGCT<br>CTCTGGACGAGCTTAAGGAACACGTTTCAGCGAACTATGGAGACGCATCGCTCCTGTT<br>CCGTTTCGGGGTACTGATCACTAACGCATGGTACGATGCTTCTGCGCCGTACCATCCTA<br>CGGCTGTAGGAGTTTACTCTCACCTTGGTCGGAGACCTGCTGAGGAAGCCACAAATCG<br>CAACATCAACACAGCTGTGATCTACGCTTCATACAGAGTGCTGAACTCGTTCATGCCGA<br>CGTACAAGGCCAGTTGGCGGAAGATGCTTCTGGATATAGGTTTGGACCCAGATAATAATA<br>GCACAGATCTGAACACGCCAATCGGCATTGGCAACGCAGCTGGGTTCGCAGTGGTTGC<br>AGGTCGTCAATTCGATGGAATGAATCAGGAAGGAGACACCAACAAGCAATACAACCCTA<br>TGCCGTATGCAGACTACACCCAGTACAAACCTTTAAACACAGCTTACAAGTTAAAGTCAC<br>CCAGTCACTGGCAACCGGACATGCAGCGGAAGGGACTTGGCCTGTACAAGATACAACA<br>ATTCGTAACGCCCCAATTCGCACTGGCCGAGCCGTACTCCTACGACGACCCCAACGAC<br>TTCGAGGTTCCGCCACCGTATAATTCAAACTTTAAGAACAAACGAGCGTATCGCAAGCA<br>GGCAAAAGAGGTGCTGGCTGCGTCAGCAAACCTCACAGACGAACAGAAGATCAAGGC<br>GGAACTTTTCGACGACAAGTTTCGGTCCCTGAGTTACTCCTTGAGCTCTAACATCGGGC<br>CACGGAACTTATCGCTGCTGGAGTTTATACAGATAGGGTTCATGACTAACCTTGCCGCTT<br>TCGACGCTGGAATATTCGTGTGGCAGGAGAAATACCGCTTCGACGCGGTCCGTCCGTT<br>CTCCGCAATACGCAAACTTTATAAGGATGAGAGCGTACAGGCATGGGGCGGACCCGGT<br>AAAGGTACGGTGTCCATGCAGGGAGAGGAGTGGCACTCATACCTTGAGGAAGCTGATC<br>ACCCCGAGTATCCGTCTGCCACGGCGTGCTTCTGCAACGCGCACGCCCAGAGTCTGC<br>GAAAGCACTTCGGCGACGATAACATGGTCTACTATTTGCCGATACCAGCGGGGTCGTCC<br>CGCGTCGAGCCCGGGGTTACGCCGAAGAACGATATAGTTATTAGTTACAACAACTGGAC<br>AGACTTCGCCAAGGAATGTGGCCAAAGTAGAGTATGGGGCGGTGTACACTTCCAAGCA<br>GCCGTCGACCAGTCTGCCGAGGTTTGCCCCGTTTTCGGGGACCTGGCCTACGAGTAT<br>GTTAGTTCCCTGGTGGACGGCTCAGCCAAATTGCGTGCCCCGTCGCAGGGTCGGCCG<br>CTTTCAGAACTGCCCGAACGGTTTTCATTCGAGAACGACGACGACTAG |
| WP_141240123.1 | *Pseudoalteromonas* sp. HM-SA03 | SA03-VHPO | ATGGCACAGGTCAAACAGGACTTTGATTTTGACGTAGACAACGCTGCCCTGGATGTGG<br>TTGGCTTCGGCAGTCTTCCGGCGTTACGTGAGCATGTGTCAGCAAATTACGGCGACGC<br>CTCGTTAGTGTTTCGTTATGCCATTCTCCTCACGAACGCCTGGTATGATGCCACTGCAG<br>CGTACCACCCGACAGCCGTAGGCGTGTACAGCCATTTAGGGCGCCGCCCGCAGTCTG<br>AGGCGACCAACCGCAATATTAACATTGCAGTGTTCTATGCATCCTATCATGTTCTTAACTC<br>GTATATGCCGACCTACAAGCCTACGTGGCGTAAACTCCTGCTGGACGTGGGGCTTGAC<br>CCGGACAATAACTCAACCGATATTACCACCCCGATTGGCCTTGGGAATGTTGCAGGTAA<br>AGCTGTCGTTGCTGGTCGCTTGAATGATGGTATGAACCAATACGGCGACGTGGGCCGC<br>ACTTATAATCAGACTCCGTACGCAGATTATACAAATTATAAGCCTGTCAACACAGCGTATA<br>AACTGCGTAACCCCTCACGTTGGCAACCCGACATGCAGCGTAAAGGATTGGGCCTGTA<br>CAAAATTCAACAGTTTGTTACGCCCCAGTTCGCGCTGGCCGAACCCTATTCGTATGAGG<br>ATCCGCGCGACTTTGAAATGCCGCCCCCGCATAATAGCAACCATCGCAATCGCCGCGC<br>ATATCGTCAACAGGCCAAGGAAGTGCTGGACGTATCAGCAAATCTTACGGATGAGCTGA |

| | | | |
|---|---|---|---|
| | | | AGCTCAAGGCCGAACTGTTCGACGATAAGTTTTCTTCTCTGTCTTACAGCGTTGCATCC<br>AATACAACGGCCCGCAATCTGCCGCTGTTAGATTTCATGCAATTCGAGTTTGTCGTTAAT<br>ATGGCAACCTTCGATGCAGGGATTTTCGTGTGGCAGGAGAAATACCGTTATGACGCGGT<br>GCGTCCCTTCAGCGCCATTCAGAAACTGTTCAAAGATAATCTGGTCGAAGCCTGGGGT<br>GGCCCAGGTAAGGGCACCGTTACGATGAAAGGGTCGGACTGGAAGTCTTATCTCGAAG<br>AAGCGGATCATCCTGAGTACCCGAGCGCCAGCACGTGTTTCTGTTATGCGCACGCACA<br>GGCAGCCCGCCTGTATTTCGGTGACGACAACATGGTGTTTAACTTAACCTACCCCGCG<br>GGCAAGAGCCGTATCGATCCGGGTGTAACTCCGGCCAAAGATACCCTGATTACGTATAA<br>CACCTGGTCTAAATTTGCCCAGGAATGCGGCCAGTCTCGTGTGTGGGGCGGCGTGCAT<br>TTCCAAGCTGCTGTAGACGAATCAGCGGAATTCTGCCCAATGTTCGGCAACTTAGCCTT<br>TGACTACATGACAAAATTGGTAAATGGCACAGCTGAACTGCGTGCGCCGTCCAAGGGC<br>CGTGATTTGGATACCCTGCCGGAACGTTTTAAGTTCGGGAAAAGTTCGGATAAAGGCGA<br>TGATGACGAAGATGAGTCGTGA |
| WP_128198684.1 | *Rubrivivax albus* | raVHPO | ATGAAAAGTCCGATTAGCTTCCGTTTAGGTCGTCGCCTCGCAGGTGCTGCGGCCGCGG<br>CCGTCGCTGCATGGTCTTGTAGTTTACCTGCCCAAGCTAGAACATTTGATTTTCAAACG<br>GGAAATGCTCCTATTGATGTTATTATTCCTGCTGTGGTCCCTTCTATATTTGCTGCGGTCG<br>CACCTTCGGATGCTCCACTTGTATTACGAACGACAACTCTCTTAACAAACGCTTGGTTTG<br>ATGCTTTAGCTCCGTATCATGCAACCGCAGTCGGTGTTTATTCTAGACTCCCACGGCGC<br>CCTGCTGACGAGGGTGTTGATGACCGTAATCGGAATGTTGCGATATTTTATGCTGCACT<br>GCCAGTATTAGAATCGTTATATCCCGCGGAAAAGGCGCGGTGGGCAGCATTACTTCAAA<br>GCGTTGGACTTGATCCCGCACTTCAAAGCGATGACCTTTCGACCCCGGAAGGTATAGG<br>GAGAGCAGCAGGTTTGGCTGTCGTTCGTGTTCGTGAAGCAGATGGTATGAATCAACTG<br>GGCGACGCAGGTGGACGTCAATTTAACCGTGTCCCGTATGCAGATACAACAGGGTATG<br>CACCTGTAAATACTGCTGATGCTTTAAAAGATCCAAGTCGATGGCAACCTGCTGTAGTGT<br>CACGTGGGAATGGGATATTTACTGATCAACGGTTTGTAACACCACAATGGGCTCAAACAT<br>TGCCTTATAGTCTCCCACCTCGTCGTCCGTTCGTGGTTAGACCGCCTGTAGATTCAAAT<br>CCTAAAGGACCGGGCGGTCTTGCCCGATATAAAGCGCAAGCTGACACCGTTCTCGCAG<br>CTAGTGCACAACTGACAGATGCGCAGAAAATGACTGCTGAGTTGTTTAATAATAAGATTG<br>AATCACTCGGATTTGTCGCGTTATTTCTCTCTGTATCTCAAGGTTGGTCCGTTGAACGCT<br>TCGTACAATATGATTTTCTTGTAAATCTCGCAGCCTTTGACGGTGGTATTACAATGTGGAG<br>AAATAAAGCACGATATGATGCTGTCCGTCCGTTTTCAGCAATACGGTATCTCTATGGTGA<br>TAATCCTGTAACCGCGTGGGGTGGCCCGGGTCGTGGGACCGTTGAAATTCCAGCATCT<br>CAATGGCAAGCATATTTGGGAGTTGCTGATCATCCTGAGTATCCTAGCGGCTCAACGTG<br>CTTTTGTGCAGCACATGCACAAGCAAGTAGACGTTATTTAGGTACCGATACCTTAGGTTG<br>GGTAGTCCCATATGCCAGAGGTTCGTCACGTGTTGAACCAGGGATAACTCCGACAGCT<br>GATCTCCAATTGAGTTTCCCAACCTGGAGCCAATTTGAAACTGATTGCGGTCAATCCCG<br>TTTATGGGGTGGAGTTCATTTTCAAGCTGCAATAGATGCAGCTGTTCCAGTTTGTCGTAG<br>AGTTGGTGACCTCGCTTATGATTTTCTTACGCGACATATTGATGGGACCGCACCTGCAC<br>CTGCCCGAGTTGATCGCCGTTAG |

| | | | |
|---|---|---|---|
| WP_132875434.1 | *Tamaricihabitans halophyticus* | thVHPO | ATGTACTCACGAAGACGGATAGGATTGGCGTGCGCTGCGCTTGCTGGGCTGTTGGCTT<br>TTAGTTTGGGAGCTCCTGCATCGACTGCATCAAAACATACAGCTGCATTTGATTTGGATC<br>ATGGAAATGCAGCCACGGAATACGCCATACCGATTGTCCAGCCAGTGATTGAAGAATTAT<br>CTCCTGGTGGTAATGACTTAACAATTATTCTCCATTGGACATCTCAAATTACGGCTGCGT<br>GGTTCGATGCCGTAGCCCCTTACCATCCCACAGCTGTCGGAATTGCTTCACGCCTTCCC<br>CGTCAAGAACGTACAGAAGCTACTAATCGCAATGTGAATATTGCTTTACTGTATGCATCCT<br>ATCAAATTTTCTCTTCCCTTGACCCTCAAGAATCGGAGCGGTGGCGCAAAATGCTTACG<br>GACGTAGGTTTAGACCCAGACGATCAAACAACCAACCCTAATACTCCTGTAGGCATAGG<br>GAATAGCGCTGGTCGTGCCCTGGCGGAATCAAGACGTCACGATGGTATGAATCAACAT<br>GGTGATGAAGGTGGTCGTAAATACCATCGGACACCATTCGCGGATTATACGAATTACCA<br>GCCTGTCAATACTGCAGATGAACTTACGGACCCAACTCGATGGCAGCCTGCGGTTGTTA<br>CTGACGATAATGGTATTTATCGCCAACAACGATTTGTAACTCCTCAAATGCGTCTTGCAG<br>ATCCATATACTTATAGTAGCCCCGAGAAATTTCGTGCTCCGGCTCCTATTGCACGTAAAG<br>ATGGGGCCGATTCCGCAGAATATAGAGCTCAAGCAGATAATGTCCTTCATCACTCTGCG<br>AAGCTCACTGAGAGACAGAAATTGGTAGCAGAAACTTTTGATCATAAGTTTGTGTCGATT<br>GGGAATGCGTCCTTGTTTCTGTCCGAAAAGTTGGATCTCTCCATAATAGAATCCGTTCAA<br>CAAGAGGCCCTGTTCACGTTGTCAAATTATGATACGACTATTGCTGTCTGGCAAGAGAA<br>GCATAGATACGATGCAGTCCGTCCATTTTCAGCTATTGCTCATTTGTATGGTGATAAGCC<br>GGTCCAAGCCTGGGGTGGCCCGGGTAAAGGAACTGTTGCTGACATGCCTGCTAATCAA<br>TGGCGCAGTTATATAAACTCTGCGGATCATCCGGAATATCCGAGTGGCTCCGCTGCTGT<br>GTGTGCTTCTCAAGCCCAAATGGCACGTAATATGTACCAAGATGATCAACTTGGTTTTCC<br>AGTGGAATATGCAAAAGGGTCGAGTAGAATAGAACCAGGCTCGACTCCGTCCTCAGAC<br>ACTACACTTAGATTTGATACTTGGAGCGAGCTTGAGGAACAGTGTGGACAATCTCGCGT<br>CTGGGGTGGTGTTCATTTTCAGCAAGCCGTCGATGCGGGTGCCGATCTTGGCCATCAA<br>ATAGCTGACGTTGCATATCGCACCATGCGTGGATATGTCGCTGGTACGGCTCCGCCTTT<br>AACCGCAGGAAAATAA |
| WP_093031253.1 | *Thiocapsa roseopersicina* | trVHPO | ATGAAGAAGGGAAAGGCATTAAGCATCTCTATGCTTTCACTGGTACTGCTGCAAGCTCC<br>AAACATTACTGACGCGTTTGACTTTGATTCGGGCAACGCTGCTGTGGAAATAGTTATCC<br>CAACCGTTGCCCCAGTAATTTTCACAGTAGTGTCCCCGTCCGGCGGAGACGCGACTCT<br>GGTCCTGAGAGTTACTACAATGACTACCAACGCATGGTTTGATGCGACGGCCCCATATC<br>ATCCCACTGGTGTGGGAGTCTATACCCGCTTGGACCGAAGACCATCAGCAGAGAGTGC<br>AACTAATCGAAACCCCAATATTGCACTGTTGTACGCCTCTTACCACGTGTTTATGTCCTTA<br>CTCCCGGACCAAGAACAAACGTGGCGTGATATGCTCTCTAATTTAGGTCTGGATCCGGA<br>TGCAGAAGCGGATGATCCAGATCCCGCTATTCGCTTGGGCATCTTGGGTGGTACTGGC<br>GTTGTACAGGGGCGACAGAATGATGGTATGAATCAATTGGGGTTCGACGGCGGGCAAC<br>TGTATAATCCGACGCCGTACCGTGATTATACAGGTTATGCACCCGTCAATACTGCCTATG<br>AATTGAAAGACCCCAGTAGATGGCAGCCAGATCTTCAGCGTCAGGGAATGGGTCTTTAT<br>AAAGTTCAACAGTTTGTTACGCCTCAGTACGCTTCTGTTGAACCTTATTCCTTTGTGGAT<br>GCATCTGAATTTTCTGTTCCCCCACCAATAAATTCCAAGAAGGCCCCCAAGTTTCGCTAT<br>AAGAACCAAGCTGATGAAGTTCTGCAGGCTAGTGCGAATCTTACGGACGAACAGAAAC |

| | | | |
|---|---|---|---|
| | | | TGCGTGCTGAATTGTTTGATAATAAGATAGAATCACTCGGTTTTAGCGCGGTCTTCGCAG<br>CCCAATCTCGCGGTTTGTCCTTGTTGGATTTTATTCACCTGGACTTTCTGACAAATATGG<br>CGGCCTTTGATGCAGGAATTGTTGTTTGGCAGGAGAAAAGAAGATATGACGCTGTTCGC<br>CCATTTTCTGCCATAAGAGAAGTGTATGGAGATGACCCAGTAACGGCCTGGGGTGGAC<br>CTGGACAAGGTACTGTATTCGACTTGCCCGCCACGGAATGGAAATCCTACCTTGAGGTT<br>GCAGACCATCCTGAATACCCGAGTGGATCTGCGTGCTTTTGCGCCGCCCACGCTGAAG<br>CGGCACGCTTATTTCTTGGTACAGATGACCTGACAGTTGGTGGAGCTTTCCCCGGTTTC<br>GTTATTGAGCGCCCAGCTGGGAGCAGTAGAATAGAGCCAGGTGTAACCCCGGCCATTG<br>CTACTAGCCTGGTTTTCCCAAATTGGACCCAATTCGCCGCCGACTGTGGTCAATCCCGT<br>GTTTGGGCTGGTGTGCATTTTCAAAGCGCTGTGGACGAATCTGTAAAGCTGTGTGACG<br>TCTTTGGGGAAAAGGCTTACGATTACGTGAACGCCCTGATAGAGGGTGTGGCTCCAGA<br>AAGAGGCCCTTCCCAGGGTCGGTTTTAA |
| WP_017029669.<br>1 | *Vibrio breoganii*<br>1C10 | vbVHPO | ATGAATCCGCTCTCTCGTCGTTCTTTCGCGACATTCTGGTTCACGTTAGCCTCGGTTGT<br>ATTCGCTGTAGGGGCTGAATCCGCTCAAGCACAAACATTTGACTTTGATAACGGATCAG<br>CACCGGTCGAAGTTATTATTCCGGCCATAATACCAATTGTACTGAATGATGTAAGTGCTG<br>GCGCCAATGACGCCACCCTCGTGTTGCGCATCACGTCGGAAATTGCGGTTGGTTGGTT<br>TGATGCTATCGCGCCATACCATCCGACAGCACGAGGCATTTACAGTAATTTGCCTCGCC<br>GTCCACCGAATGAATCTCAGAGTAATTCGAACAAGAACATTGCCATGTTATACGCTAGCC<br>GTAGAATACTTATGTCATTACTGCCTGATAGAGCATCGGATTGGGATAATCTTCTCACGTC<br>CGTCGGCCTGGACCCTAATAATGGAAGTCAAAACCTTGCGAGCCCCGTAGGCATCGGC<br>AATGTAGCGGGTAACGCTGTGGTCGCGTTTAGAGAGCATGATGGTATGAATCAGTTGGG<br>AGATGAGGGCGGCCGAACATTCAACAGATTCCCATATATGGACTCAACGGGTTTTAAAC<br>CTGTTAATACGGCTTATGAAATACTGGACCCAACACGCTGGCAACCCTTAGTTGTGACA<br>CAAGGGAATGGAATTTTCCGAGTCCAACAGTATGTGACGCCTCAACTTGCACGGACGC<br>GTCCGTATGCTATAGATCCCCGTAACTTTTCCTCACCAGAACCTATTGCTAGCCTTCTTAA<br>GCGCGGCAAGAACGGACGTGATGCGTATAAACAACAGGTAGATGAGGTACTGGCCATC<br>AGTGCCAGCCTGAATGACTATACGAAAGCCCAAGCCGAATTCTTTGATAATAAGTTTCGT<br>TCAATCGGCTTTGTAGCCCTCTTTGTAGCAGGGGTAAATCAGTTTTCGCTGGATGAATTG<br>GTACAATATGATGCAATGCTGAATATGGCTGCCTTCGATGGGGCTATTGTCAGCTGGAAA<br>CAGAAAACCAGATGGAATGCCGTAAGACCCATGACCGCCGTTAGACACGTGTATGGGG<br>ACCGTACAATCACCGCATGGGGCGGACCAGGGAAGGGTACACAACGACTCCCGGCCA<br>ATGAGTGGCAGTCTTATTTACAAACCCCGGATCATCCCGAATACCCGAGCGGCTCTAGT<br>TGTTTCTGTGCAGCTGAAGCACAGGCGACACGCCGTCTGCTGGGGTCGGATGAGTTG<br>GGATGGCAGGTGCTGGTTCCAAAGGGTCAGAGTCGTATTGAACCTGGGATAACTCCAT<br>CCCAAGATACGACTTTAGTCTTTAATACCTGGACTGATTTTGAAAATCAATGCGCACAAG<br>CCCGAGTTTTAGGTGGCGTACATTTTCAAGCTTCCGTGGCAGCGAGCATAGCTATGTGC<br>CGGCCCGCAGGGGATGCCGCATATGAATTCATACAACGTCACGTTAATGGTCTCCCTTA<br>A |

| | | | |
|---|---|---|---|
| WP_155759790.1 | *Vibrio natriegens* CCUG 16374 | vnVHPO | ATGAAGTTAAAGCCTTTGAGCCTGAAACTGGCGACGTCTACATTACTGCTGAGTGTTCC AGGAGTCTCGTACGCATTCGAATGTCCTCAAAACTTCCCAATCTGTGCCCCTGCTACTG TAATCGAGGCTGCGGTTCCCCGCCTGTTAAGTACCGTGAGTCCGGGCGACGCCCCGC TCATTCTTCGTACCACAACCCTGATTACAGCAGCCTGGTTCGATGCCATCGCGCCGTAC TCCGACACGGCAATAGGAGTCCACAGCGATCTGGGACGTTTAAATGCCGGCGATAGCG ACTCCGCCCGTAATATTGCTATAATGTATGCGTCCCACAAGGTCTTGAAGTCATTAATGC CCCAGTTCGCGTCCGATTGGGACGCAATGCTTTTAACCGTCGGACTTGATCCCGACGA TAACACGTTGGACGCAAATACTAGTATTGGTGTCGGGAATTTAGCCGGCAATGCAGTGG TCGCTGCCCGGGTTGACGACGGAATGAATCAACTCGGTACGGATTGCAATCAGTACTAC CCGAAACCATATGCAGACTGTACGGGATACCGCCCCGTAAATACAGCTGAGAAGCTGAC TGATCCGCGCCGTTGGCAACCTGACACCTTAACTAGCGGGTCAGGGATTTTCTTTAGCC AGCAGTTCGTTACTCCCCAGCTCGGTTCCACACAGCCATTCTCTTATGATAAGCCTACTA TTACCGTTCCCAAGCCGACTCGCAGTTACGCAATTAACCCTAGCGGTAAGGCCTTCCCT GATTACATTGAACAGGCAAACGAGGTCCTCGAGGCGCAGGCCGATCTGACCGACGAG CAGAAACTGTTCTCTGAATTCTACGATAACAAGATTATCTCGCTGGGGTACTCCACTTTG GCGGCCACGTTTTACCATCATTTATCGCTTGAGCAGTTCGTACAACTGGACTATCTCGTT AATGTAGCGGCTTTTGACGCGGGTATCACGGTATGGGATAACAAACGTAAGTATGATGC GGTTCGTCCTTTCAGCGCAATAGGATACCTTTACAAGAATGATAATCTGACAGCTTGGGG CGGACCGGGCATGGGCACCGTTTCCGACCTGAAGGGATCTGAATGGCGGCCCTACCT GCAGTCTGCCAACCATCCTGAGTATCCAAGCGCGAGCGCATCCTTCTGTCGGGCCCAC GCGACGTCCACTAAACGGTATCTGCAAGAGGAACTCGGTTTACCGTATGAGGCGGCGA ATGACCTCTCTTTCTGGCAGCCAGGTTTCTTACCGGAGCAATTCGAGGTTTTGGACAAA CAGTCAGGTACGAGCCTTATAGAGCCGGGAGTCACCCCTCAGAGCCATACAGTGCTGG GGTGGTCTACGTGGGATGAGTTTTCAGACAAATGCGGCTTGTCAAGACTCTGGGCAGG AGTGCATTTCTACGATAGCATACCCGCAGGACAGTCTATTGGTGACCCAATCGGATCTG CCGCGTATGACTGGCTGAAGACCTATATTAACGGAACTGCCCAGTGA |

**Table S4 – Amino acid sequences for selected alkyl quinolone VHPOs**

| Accession no. | Organism | Abbreviation | Sequence |
|---|---|---|---|
| WP_052553269.<br>1 | *Enhygromyxa salina* | esVHPO | MQSTRFSTSSSLLLLGALATAACDPTLDPAATRVAASDEGVNDQECTTLLPANVPGLQAQL<br>PTQTMGADFDFDTGNAPIEIVIPAVLPVIAGSVAPGDATIVLRFTTMLSNAWFDATAPYHPTA<br>VGVYSNLGRRPASESTTHANMNIAILYASYRTLNSLAPQHAADWDALMVSLGLDPHDDHES<br>TTDPIGIGNAAAAALLAVRENDGFNQLGFEGGREYNPIPYADYTGYEPRNTRFEIKDERRW<br>QPAIVTSRYGITRAQHFVTPQYALTLPYSYDDPQDFGVPLPDKSLKKGSHAKKKYRAQADE<br>VLEVSANLTDEQKVTAELFEDKIRSLGFSALFVSLSSGHSLLDFVHYDFLTNLAAFDVGIVVW<br>QEKTQYDAVRPFTAIRHIYGDDEITAWGGPGQGTVNDLPANEWRSYLDVADHPEYPSASA<br>AFCAAHAQASRLFLGTDDLGWTVPIPAGSSIVEPAITPAADLNLHFPTFTDFATRCGYSRLW<br>GGVHFEDAILASFELGDEIGAGAYEFVQAHIDGTPP |
| WP_157898942.<br>1 | *Luteitalea pratensis* | lpVHPO | MCRVIPRIAFCTAALVLMPARPQAQAPPFDFSSGNIGIEVIIPAVIPALFQTTSPNDAPIILRHTT<br>LITNAWFDAIAPYHPTAKGVYTRLENRPPAEATTRNKNIAMAYATYRLLNRLMPRFAADWRQ<br>MLVSVGLDPDDASVDVRTPIGIGNVAGSAVAVARERDGMNQLGDEGGRVYDLRPYADYTG<br>YMPVNTPWKLSNPSRWQPLFTTPGNGTFLGQQFATPQWGATTPYSYGNPAAFGTPPPID<br>SNHHRRQAYEAQADEVMDAQATLTDYQKMVAELFDNKITGLGFAALFIAQSRNMSLDEFVH<br>YDFLTNVAAFDGGITTWRDKYLYDAVRPVTAIRYLNRGRTITGWAGPGRGIVNDLPANEWR<br>SYLNTANHPEYPSGSSCFCAAHAQASRRYLGSDQFGWSVPKPAGSSVIEPGVTPAADVVL<br>GPWQTFTEFEEECGMSRLWGGVHFRPAIEEARNSCRQVGDKAFEFLQRKLAGQ |
| WP_139559104.<br>1 | *Methylotetracoccus oryzae* | moVHPO | MNPLSRRSFATFWFTLASVVFAVGAESAQAQTFDFDNGSAPVEVIIPAIIPIVLNDVSAGAND<br>ATLVLRITSEIAVGWFDAIAPYHPTARGIYSNLPRRPPNESQSNSNKNIAMLYASRRILMSLLP<br>DRASDWDNLLTSVGLDPNNGSQNLASPVGIGNVAGNAVVAFREHDGMNQLGDEGGRTFN<br>RFPYMDSTGFKPVNTAYEILDPTRWQPLVVTQGNGIFRVQQYVTPQLARTRPYAIDPRNFS<br>SPEPIASLLKRGKNGRDAYKQQVDEVLAISASLNDYTKAQAEFFDNKFRSIGFVALFVAGVN<br>QFSLDELVQYDAMLNMAAFDGAIVSWKQKTRWNAVRPMTAVRHVYGDRTITAWGGPGKG<br>TQRLPANEWQSYLQTPDHPEYPSGSSCFCAAEAQATRRLLGSDELGWQVLVPKGQSRIEP<br>GITPSQDTTLVFNTWTDFENQCAQARVLGGVHFQASVAASIAMCRPAGDAAYEFIQRHVN<br>GLP |
| WP_183457526.<br>1 | *Microbulbifer rhizosphereae* | mrVHPO | MGKLFHKGVSKVLCCTMVIVIGCFTKSFASESETSYDFKNGNAVIEVAVPALLPVILSDVSST<br>AGDATLILRMSTLMSNAWFDASAPYHPTAVGVYSRLGRRPLSESETNTNINIAILYASYRVLS<br>SLLPQRTPVWRKMLTDIGLDPDDDSVDLATPVGIGNAAGLGVVRGRARDGMNQLGDESER<br>AFNPMPYMDYTNYRPINSAYRLFDPSRWQPDLQRKGMGLYKIQQFVTPQYALTEPYSYQA<br>PQRFNMPPPYASNHWNYPAYKQQADEVLAASADLNDERKLKAELFDNKIESLGFSALSTAI<br>SRGLSLQETIELDFLISMAAHDAGIVIWQEKRRYDAVRPFSAIRYLYGDEWVSAWGGPGRG<br>TIRLPANQWKSYLEEADHPEYPSASTCFCAAHAQAARRYLGTDVLNWEFSKAAGSSHVEP<br>GITPAEDTILRFDTWTDFVSDCGQSRVWAGVHFQAAVDVSTQMCDVFGDMAHEYLSSLID<br>GSAPVRRPAKGKRYK |

| | | | |
|---|---|---|---|
| WP_108732878.1 | *Microbulbifer* sp. A4B17 | A4B17-VHPO | MLKNIMYKNALKSIAGILLVSASWQAQAISCPQLTDVSLPPKCSSEELATCNAAVQNVIPAAACAIFEVSPAANDATLVLRFTTLITNSWFDAAAPYHPTAKGVYSDIARLTEPTDNTELNIALSYASHKVLSGLFPHLVSEWDDMLIELGLDPGNLSEDTTTPIGIGNYAGRKVLEGRANDGMNQLGNEGNQLYHRLPYADYTDFKPANSAYKLRFPSKWQPAVLMDRPGVFRVQQFVTPQIALTTPYSYESPEEFTSPVPLKSKIWNYPSYVSQVNDVLATSANLTDEQKMKAELFDMKIFSLGFAAVFASESQGLSLMDFIHYDFLTNMAAFDTAITIWKEKRRHNSVRPFSAVRYIYKDEPVTAWGGVGKGTVNDLPATQWTSYLPVADHPEYPSASASFCAAHAETSRLFLPQGDTLGYLVTALAGSSQIEPGITPQTDLNLYFPTWTSFEEDCGNSRVWGGVHFEPSVPAGQAIGRQIAHRAYEFYLSKLAGN |
| WP_156035230.1 | *Microbulbifer* sp. HZ11 | HZ11-VHPO | MKIGMLLRAVLCGTALAFCALNTQAQPTTPPPVDLDNGNAAIEAVIPYVAPVTFEYVSATGGDATLVLRITTQITNAWFDASAPYHPTAVGVYSRLGRRPANESANNRNINIALLFASYRVLNTLLPLQHATWRAMLEAQGLDPDDNSTDLTSPIGIGNAAGAAVAKGRLRDGMNQVGDETRSGPLRAANPMPYMDYTGYQPINTAYTLFNPSRWQPDIQRKGMGLYKIQQFVTPQYALTEPYSYKSPRKYRALPPFASFHWNFAAYKQQADEVLAASANLTDEQKLMAELFDNKIESLGFSAGFAALSQGLSLMEFIHFDFLTNMAAHDAGIFVWQEKRRYDAVRPFSAIQYIYGKQPVTAWGGPGQGTQSIPANTWKSYLEEADHPEYPSASTCFCTAHAQSARQYLGSDALNWSVPYTAGSSRIEPGVTPANDMTLKFPTWTSLEENCGQSRVWAGVHFQAAVDEGRRICGVFGDNAYNYLQTLVAGTAPEREPADRLRGRRK |
| WP_076514445.1 | *Oleibacter marinus* | omVHPO | MKLPLLKGRPLALAASALLSIAVNQASAYDLQTGNAPVELVISKVAPAIFQDISATAGDATLVLRVTAQVTNAWYDASAPYHPTAVGVYSRIPNRPASESVDNENINIATLYASYQVLKVLLPQRTAEWRGMLVEAGLDPDDTSTDLTTPVGIGNVAGAAVAEGRLNDGMNQAGWINEDVHPQPYADYTGYEPKNTAYDLKRPSKWQPDVQRKGLGLYKVQQFVTPQYANVEPYSYDDPERFSVPWPANSFAPFKHLYKQQADEVLVASANLTEEQKLKAELFDNKIFSLGFSAVFAAQSRGLSLLEFIQLDFLTNMAAFDAGIFVWQEKAEYDGVRPFSAIQHIYGDQPVEAWGGPGEGTVMLPANQWRAYMEEADHPEYPSASSCFCAAHSQSARAFLGDDMLGFPVEYPAGSSRIEPGLTPAADTTLVFNTWSEFEQDCGQSRVWAGVHFQAAVDESLALCGDFGDMAYDYMQSLIDGTAPARGPSQPLRDPATYSNKKWW |
| KZN74784.1 | *Pseudoalteromonas luteovioleacea* H33-S | plVHPO | MNKYLSKISLLHQIRKLMKLRNISVFLLSLICVTHVQANHLDEFDFDKDNGALDVVGWGSLDELKEHVSANYGDASLLFRFGVLITNAWYDASAPYHPTAVGVYSHLGRRPAEEATNRNINTAVIYASYRVLNSFMPTYKASWRKMLLDIGLDPDNNSTDLNTPIGIGNAAGFAVVAGRQFDGMNQEGDTNKQYNPMPYADYTQYKPLNTAYKLKSPSHWQPDMQRKGLGLYKIQQFVTPQFALAEPYSYDDPNDFEVPPPYNSNFKNKRAYRKQAKEVLAASANLTDEQKIKAELFDDKFRSLSYSLSSNIGPRNLSLLEFIQIGFMTNLAAFDAGIFVWQEKYRFDAVRPFSAIRKLYKDESVQAWGGPGKGTVSMQGEEWHSYLEEADHPEYPSATACFCNAHAQSLRKHFGDDNMVYYLPIPAGSSRVEPGVTPKNDIVISYNNWTDFAKECGQSRVWGGVHFQAAVDQSAEVCPVFGDLAYEYVSSLVDGSAKLRAPSQGRPLSELPERFSFENDDD |

| | | | |
|---|---|---|---|
| WP_141240123.1 | *Pseudoalteromonas* sp. HM-SA03 | SA03-VHPO | MAQVKQDFDFDVDNAALDVVGFGSLPALREHVSANYGDASLVFRYAILLTNAWYDATAAYHPTAVGVYSHLGRRPQSEATNRNINIAVFYASYHVLNSYMPTYKPTWRKLLLDVGLDPDNNSTDITTPIGLGNVAGKAVVAGRLNDGMNQYGDVGRTYNQTPYADYTNYKPVNTAYKLRNPSRWQPDMQRKGLGLYKIQQFVTPQFALAEPYSYEDPRDFEMPPPHNSNHRNRRAYRQQAKEVLDVSANLTDELKLKAELFDDKFSSLSYSVASNTTARNLPLLDFMQFEFVVNMATFDAGIFVWQEKYRYDAVRPFSAIQKLFKDNLVEAWGGPGKGTVTMKGSDWKSYLEEADHPEYPSASTCFCYAHAQAARLYFGDDNMVFNLTYPAGKSRIDPGVTPAKDTLITYNTWSKFAQECGQSRVWGGVHFQAAVDESAEFCPMFGNLAFDYMTKLVNGTAELRAPSKGRDLDTLPERFKFGKSSDKGDDDEDES |
| WP_128198684.1 | *Rubrivivax albus* | raVHPO | MCRVIPRIAFCTAALVLMPARPQAQAPPFDFSSGNIGIEVIIPAVIPALFQTTSPNDAPIILRHTTLITNAWFDAIAPYHPTAKGVYTRLENRPPAEATTRNKNIAMAYATYRLLNRLMPRFAADWRQMLVSVGLDPDDASVDVRTPIGIGNVAGSAVAVARERDGMNQLGDEGGRVYDLRPYADYTGYMPVNTPWKLSNPSRWQPLFTTPGNGTFLGQQFATPQWGATTPYSYGNPAAFGTPPPIDSNHHRRQAYEAQADEVMDAQATLTDYQKMVAELFDNKITGLGFAALFIAQSRNMSLDEFVHYDFLTNVAAFDGGITTWRDKYLYDAVRPVTAIRYLNRGRTITGWAGPGRGIVNDLPANEWRSYLNTANHPEYPSGSSCFCAAHAQASRRYLGSDQFGWSVPKPAGSSVIEPGVTPAADVVLGPWQTFTEFEEECGMSRLWGGVHFRPAIEEARNSCRQVGDKAFEFLQRKLAGQ |
| WP_132875434.1 | *Tamaricihabitans halophyticus* | thVHPO | MYSRRRIGLACAALAGLLAFSLGAPASTASKHTAAFDLDHGNAATEYAIPIVQPVIEELSPGGNDLTIILHWTSQITAAWFDAVAPYHPTAVGIASRLPRQERTEATNRNVNIALLYASYQIFSSLDPQESERWRKMLTDVGLDPDDQTTNPNTPVGIGNSAGRALAESRRHDGMNQHGDEGGRKYHRTPFADYTNYQPVNTADELTDPTRWQPAVVTDDNGIYRQQRFVTPQMRLADPYTYSSPEKFRAPAPIARKDGADSAEYRAQADNVLHHSAKLTERQKLVAETFDHKFVSIGNASLFLSEKLDLSIIESVQQEALFTLSNYDTTIAVWQEKHRYDAVRPFSAIAHLYGDKPVQAWGGPGKGTVADMPANQWRSYINSADHPEYPSGSAAVCASQAQMARNMYQDDQLGFPVEYAKGSSRIEPGSTPSSDTTLRFDTWSELEEQCGQSRVWGGVHFQQAVDAGADLGHQIADVAYRTMRGYVAGTAPPLTAGK |
| WP_093031253.1 | *Thiocapsa roseopersicina* | trVHPO | MKKGKALSISMLSLVLLQAPNITDAFDFDSGNAAVEIVIPTVAPVIFTVVSPSGGDATLVLRVTTMTTNAWFDATAPYHPTGVGVYTRLDRRPSAESATNRNPNIALLYASYHVFMSLLPDQEQTWRDMLSNLGLDPDAEADDPDPAIRLGILGGTGVVQGRQNDGMNQLGFDGGQLYNPTPYRDYTGYAPVNTAYELKDPSRWQPDLQRQGMGLYKVQQFVTPQYASVEPYSFVDASEFSVPPPINSKKAPKFRYKNQADEVLQASANLTDEQKLRAELFDNKIESLGFSAVFAAQSRGLSLLDFIHLDFLTNMAAFDAGIVVWQEKRRYDAVRPFSAIREVYGDDPVTAWGGPGQGTVFDLPATEWKSYLEVADHPEYPSGSACFCAAHAEAARLFLGTDDLTVGGAFPGFVIERPAGSSRIEPGVTPAIATSLVFPNWTQFAADCGQSRVWAGVHFQSAVDESVKLCDVFGEKAYDYVNALIEGVAPERGPSQGRF |

| | | | |
|---|---|---|---|
| WP_017029669.1 | *Vibrio breoganii* 1C10 | vbVHPO | MNRTSLSSIVTTGLFTAFFSTSSQAFECPAGAPNICAPATVIDNTVPRLLSTVSPGDAPIILRTTTMITAAWFDAIAPYSESTVGVHSNLGRIESNYGDSDRNIAIMYASHKVLNSLMPQYAADWDQMLLSLGLDPYNQSTDLNTPVGVGNTAGNAVVTTRVQDGMNQLGTDCNIYKVRPYSDCSGYKPTNKAEKLVNSRKWQPDTLTSGGGIFFSQQFVTPQYGQTTPFSYDQPTLMAPIPTKSYAVTSDGSPLPQYRDQADEVLQSQLELTDEQKLLAEFYDNKIISLGFSTLAASIHHQLTLEQFVQLDFLVNVAAFDTGITIWDNKRHYDAVRPFSAIAYIYGSQPISAWGGPGKGNVNDMPGSDWRPYLQSANHPEYPSASASFCRAHATSTKRYLREIVGLNKKRSNDLSFWTPDLLPPEFNVLQKPAGSSVVEPGVTPAAELTLGWKTWDDFSDDCGISRLWAGVHFYDSIPAGQDIGQSIGSDAFDWLETHINGNP |
| WP_155759790.1 | *Vibrio natriegens* CCUG 16374 | vnVHPO | MKLKPLSLKLATSTLLLSVPGVSYAFECPQNFPICAPATVIEAAVPRLLSTVSPGDAPLILRTTTLITAAWFDAIAPYSDTAIGVHSDLGRLNAGDSDSARNIAIMYASHKVLKSLMPQFASDWDAMLLTVGLDPDDNTLDANTSIGVGNLAGNAVVAARVDDGMNQLGTDCNQYYPKPYADCTGYRPVNTAEKLTDPRRWQPDTLTSGSGIFFSQQFVTPQLGSTQPFSYDKPTITVPKPTRSYAINPSGKAFPDYIEQANEVLEAQADLTDEQKLFSEFYDNKIISLGYSTLAATFYHHLSLEQFVQLDYLVNVAAFDAGITVWDNKRKYDAVRPFSAIGYLYKNDNLTAWGGPGMGTVSDLKGSEWRPYLQSANHPEYPSASASFCRAHATSTKRYLQEELGLPYEAANDLSFWQPGFLPEQFEVLDKQSGTSLIEPGVTPQSHTVLGWSTWDEFSDKCGLSRLWAGVHFYDSIPAGQSIGDPIGSAAYDWLKTYINGTAQ |

**Table S5 – Primers used in this study**

| Sequence (5'->3') | Description |
|---|---|
| CATATGGCTGCCGCGCGGCAC | Amplification of pET28a (+), N-terminal 6x His tag for gene insertion, Forward |
| CTCGAGCACCACCACCACCACCACTGA | Amplification of pET28a (+), N-terminal 6x His tag for gene insertion, Reverse |
| GGTGCCGCGCGGCAGCCATATGAAAATTGGAATGCTGCTTCGTGCT | Amplification of synthetic Hz11-VHPO with Gibson assembly overhangs for N-terminal 6x His tag insertion, ATG Start, forward |
| GGTGGTGGTGGTGCTCGAGTCACTTGCGGCGGCC | Amplification of synthetic Hz11-VHPO with Gibson assembly overhangs for N-terminal 6x His tag insertion, maintains TAA Stop, reverse |
| GGTGCCGCGCGGCAGCCATATGCAGTCTACGCGCTTTAGCAC | Amplification of synthetic esVHPO with Gibson assembly overhangs for N-terminal 6x His tag insertion, ATG Start, forward |
| GGTGGTGGTGGTGCTCGAGTCAGGGTGGAGTGCCGTC | Amplification of synthetic esVHPO with Gibson assmebly overhangs for N-terminal 6x His tag insertion, maintains TAA Stop, reverse |
| GGTGCCGCGCGGCAGCCATATGTGCAGAGTTATCCCGCGTATTG | Amplification of synthetic lpVHPO with Gibson assmebly overhangs for N-terminal 6x His tag insertion, ATG Start, forward |
| GGTGGTGGTGGTGCTCGAGTCATTGACCTGCGAGTTTACGTTGAAG | Amplification of synthetic lpVHPO with Gibson assmebly overhangs for N-terminal 6x His tag insertion, maintains TAA Stop, reverse |
| CTTTAAGAAGGAGATATACCATGTGCAGAGTTATCCCGCG | Amplification of synthetic lpVHPO with Gibson assmebly overhangs for C-terminal 6x His tag insertion, ATG Start, forward |
| GGTGGTGGTGGTGCTCGAGTTGACCTGCGAGTTTACGTTGAAG | Amplification of synthetic lpVHPO with Gibson assmebly overhangs for C-terminal 6x His tag insertion, removes TAA Stop, reverse |
| GGTGCCGCGCGGCAGCCATATGAATCCGCTCTCTCGTCG | Amplification of synthetic moVHPO with Gibson assmebly overhangs for N-terminal 6x His tag insertion, ATG Start, forward |

| | |
|---|---|
| GGTGGTGGTGGTGCTCGAGTTAAGGGAGACCATTAACGTGACG | Amplification of synthetic moVHPO with Gibson assmebly overhangs for N-terminal 6x His tag insertion, maintains TAA Stop, reverse |
| CTTTAAGAAGGAGATATACCATGGGGAAACTGTTCCATAAGGG | Amplification of synthetic mrVHPO with Gibson assmebly overhangs for C-terminal 6x His tag insertion, ATG Start, forward |
| GGTGGTGGTGGTGCTCGAGCTTATATCTCTTTCCTTTCGCGGGG | Amplification of synthetic mrVHPO with Gibson assmebly overhangs for C-terminal 6x His tag insertion, removes TAA Stop, reverse |
| GGTGCCGCGCGGCAGCCATATGCTGAAGAATATCATGTATAAGAACG | Amplification of synthetic A4B17-VHPO with Gibson assmebly overhangs for N-terminal 6x His tag insertion, ATG Start, forward |
| GGTGGTGGTGGTGCTCGAGTCAGTTACCCGCCAGTTTA | Amplification of synthetic A4B17-VHPO with Gibson assmebly overhangs for N-terminal 6x His tag insertion, maintains TAA Stop, reverse |
| CTTTAAGAAGGAGATATACCATGAAATTGCCCTTGTTGAAAGGTC | Amplification of synthetic omVHPO with Gibson assmebly overhangs for N-terminal 6x His tag insertion, ATG Start, forward |
| GGTGGTGGTGGTGCTCGAGCCACCATTTCTTATTGCTATATGTCGC | Amplification of synthetic omVHPO with Gibson assmebly overhangs for N-terminal 6x His tag insertion, maintains TAA Stop, reverse |
| GGTGCCGCGCGGCAGCCATATGAAATTGCCCTTGTTGAAAGGTC | Amplification of synthetic omVHPO with Gibson assmebly overhangs for C-terminal 6x His tag insertion, ATG Start, forward |
| GGTGGTGGTGGTGCTCGAGTCACCACCATTTCTTATTGCTATATGTCG | Amplification of synthetic omVHPO with Gibson assmebly overhangs for C-terminal 6x His tag insertion, removes TAA Stop, reverse |
| GGTGCCGCGCGGCAGCCATATGAACAAGTACTTAAGCAAAATTTCACT | Amplification of synthetic plVHPO with Gibson assmebly overhangs for N-terminal 6x His tag insertion, ATG Start, forward |
| GGTGGTGGTGGTGCTCGAGCTAGTCGTCGTCGTTCTCG | Amplification of synthetic plVHPO with Gibson assmebly overhangs for N-terminal 6x |

| | |
|---|---|
| | His tag insertion, maintains TAA Stop, reverse |
| GGTGCCGCGCGGCAGCCATATGGCACAGGTCAAACAGG | Amplification of synthetic SA03-VHPO with Gibson assmebly overhangs for N-terminal 6x His tag insertion, ATG Start, forward |
| GGTGGTGGTGGTGCTCGAGTCACGACTCATCTTCGTCATCAT | Amplification of synthetic SA03-VHPO with Gibson assmebly overhangs for N-terminal 6x His tag insertion, maintains TAA Stop, reverse |
| GGTGCCGCGCGGCAGCCATATGAAAAGTCCGATTAGCTTCCGTTTAGG | Amplification of synthetic raVHPO with Gibson assmebly overhangs for N-terminal 6x His tag insertion, ATG Start, forward |
| GGTGGTGGTGGTGCTCGAGCTAACGGCGATCAACTCGGG | Amplification of synthetic raVHPO with Gibson assmebly overhangs for N-terminal 6x His tag insertion, maintains TAA Stop, reverse |
| CTTTAAGAAGGAGATATACCATGAAAAGTCCGATTAGCTTCCGTTTAG | Amplification of synthetic raVHPO with Gibson assmebly overhangs for C-terminal 6x His tag insertion, ATG Start, forward |
| GGTGGTGGTGGTGCTCGAGACGGCGATCAACTCGGG | Amplification of synthetic raVHPO with Gibson assmebly overhangs for C-terminal 6x His tag insertion, removes TAA Stop, reverse |
| GGTGCCGCGCGGCAGCCATATGTACTCACGAAGACGGATAGG | Amplification of synthetic thVHPO with Gibson assmebly overhangs for N-terminal 6x His tag insertion, ATG Start, forward |
| GGTGGTGGTGGTGCTCGAGTTATTTTCCTGCGGTTAAAGGCGG | Amplification of synthetic thVHPO with Gibson assmebly overhangs for N-terminal 6x His tag insertion, maintains TAA Stop, reverse |
| CTTTAAGAAGGAGATATACCATGTACTCACGAAGACGGATAGGATTG | Amplification of synthetic thVHPO with Gibson assmebly overhangs for C-terminal 6x His tag insertion, ATG Start, forward |
| GGTGGTGGTGGTGCTCGAGTTTTCCTGCGGTTAAAGGCGG | Amplification of synthetic thVHPO with Gibson assmebly overhangs for C-terminal 6x His tag insertion, removes TAA Stop, reverse |

| | |
|---|---|
| GGTGCCGCGCGGCAGCCATATGAAGAAGGGAAAGGCATTAAGCA | Amplification of synthetic trVHPO with Gibson assmebly overhangs for N-terminal 6x His tag insertion, ATG Start, forward |
| GGTGGTGGTGGTGCTCGAGTTAAAACCGACCCTGGGAAGG | Amplification of synthetic trVHPO with Gibson assmebly overhangs for N-terminal 6x His tag insertion, maintains TAA Stop, reverse |
| GGTGCCGCGCGGCAGCCATATGAACAGAACGAGCCTTTCCTCG | Amplification of synthetic vbVHPO with Gibson assmebly overhangs for N-terminal 6x His tag insertion, ATG Start, forward |
| GGTGGTGGTGGTGCTCGAGTCATGGGTTGCCGTTTATGTGC | Amplification of synthetic vbVHPO with Gibson assmebly overhangs for N-terminal 6x His tag insertion, maintains TAA Stop, reverse |
| GGTGCCGCGCGGCAGCCATATGAAGTTAAAGCCTTTGAGCCTG | Amplification of synthetic vnVHPO with Gibson assmebly overhangs for N-terminal 6x His tag insertion, ATG Start, forward |
| GGTGGTGGTGGTGCTCGAGTCACTGGGCAGTTCCGTTAATATAG | Amplification of synthetic vnVHPO with Gibson assmebly overhangs for N-terminal 6x His tag insertion, maintains TAA Stop, reverse |
| ATGAAACTCCCTCTTCTTAAGGGGCG | Sequencing of genomically encoded omVHPO, forward |
| TTACCACCACTTCTTGTTGCTGTATGTCG | Sequencing of genomically encoded omVHPO, reverse |
| TCTAAGGAGACTGCCGGTGA | Sequencing of genomically encoded O. marinus 16s rRNA, forward |
| TGTGATCCAGCCCCAGGTTC | Sequencing of genomically encoded O. marinus 16s rRNA, reverse |
| TCTAAGGAGACTGCCGGTGA | O. marinus qPCR 16S rRNA primer, forward |
| TGTGATCCAGCCCCAGGTTC | O. marinus qPCR 16S rRNA primer, reverse |
| ACTGGCCTCGAGCATGAAAA | O. marinus qPCR gyrA primer, forward |
| GCCGGTGTGGGATAAAGTCA | O. marinus qPCR gyrA primer, reverse |
| ACCCACAGCCGTATGCCGAC | O. marinus qPCR omVHPO primer, forward |
| TTCGCAGGCCAAGGCACACT | O. marinus qPCR omVHPO primer, reverse |

| | |
|---|---|
| ATG AAA CTT AAA CCA CTT TCT TTA AAG | Sequencing of genomically encoded vnVHPO, forward |
| TTA TTG TGC CGT ACC ATT TAT G | Sequencing of genomically encoded vnVHPO, reverse |
| TAA ACA TGG CAG TCG AGC GG | Sequencing of genomically encoded V. natriegens 16s rRNA, forward |
| AAC CAA AGT GGG TAA GCG TCC | Sequencing of genomically encoded V. natriegens 16s rRNA, reverse |
| TTG GGA AAG TTA TTT CAC AAA GGG | Sequencing of genomically encoded mrVHPO, forward |
| TTA TTT GTA CCT TTT ACC CTT GGC | Sequencing of genomically encoded mrVHPO, reverse |
| ACA TGC AAG TCG AGC GCG AAA | Sequencing of genomically encoded M. rhizosphaerae 16s rRNA, forward |
| ACT TCA CCC CAG TCA TGA ATC ACT CC | Sequencing of genomically encoded M. rhizosphaerae 16s rRNA, reverse |

**Table S6 – Predicted secretion signal peptides**

| Accession Number | Source organism | Likelihood Signal Peptide | Cleavage Site |
|---|---|---|---|
| MBM3759267.1 | Acidobacteria bacterium | 99.83% Sec/SPI | 97% btw 29 and 30 |
| MCC6989002.1 | Acidobacteria bacterium | 76.66% Sec/SPI, 13.93% Sec/SPII | 48% btw 25 and 26 |
| WP_086933921.1 | Agarilytica rhodophyticola | 99.88% Sec/SPI | 97% btw 29 and 30 |
| WP_246970656.1 | Alcanivorax sp. S6407 | 99.89% Sec/SPI | 95% btw 25 and 26 |
| WP_162906947.1 | Allorhizocola rhizosphaerae | 99.87% Sec/SPI | 97% btw 30 and 31 |
| MCE2471135.1 | Anaerolineae bacterium | 99.43% Sec/SPI | 96% btw 29 and 30 |
| MCE2473632.1 | Anaerolineae bacterium | No Predicted Signal | No Predicted Signal |
| WP_130969110.1 | Aquabacterium lacunae | 97.97% Sec/SPI | 93% btw 31 and 32 |
| MBM4778199.1 | Archangiaceae bacterium | No Predicted Signal | No Predicted Signal |
| MBL8938882.1 | Archangium sp. | 99.91% Sec/SPI | 98% btw 21 and 22 |
| MCR9160379.1 | bacterium | 99.99% Sec/SPII | 99% btw 06 and 07 |
| MBE0658962.1 | Bryobacteraceae bacterium | No Predicted Signal | No Predicted Signal |

| | | | |
|---|---|---|---|
| MBX2809551.1 | Cellvibrionaceae bacterium | 99.75% Sec/SPI | 96% btw 34 and 35 |
| MXV92527.1 | Chloroflexi bacterium | 66.09% Sec/SPI, 16.10% Sec/SPII | 65% btw 24 and 25 |
| MXX84763.1 | Chloroflexi bacterium | 99.92% Sec/SPI | 99% btw 19 and 20 |
| MYD09778.1 | Chloroflexi bacterium | 72.19% Sec/SPI, 27.15% Sec/SPII | 93% btw 31 and 32 |
| MYD09780.1 | Chloroflexi bacterium | 99.85% Sec/SPI | 97% btw 28 and 29 |
| MYD09944.1 | Chloroflexi bacterium | 90.82% Sec/SPI, 08.69% Sec/SPII | 88% btw 28 and 29 |
| MYD38911.1 | Chloroflexi bacterium | 99.92% Sec/SPI | 99% btw 19 and 20 |
| MBA2454044.1 | Chloroflexia bacterium | 99.86 Tat/SPI | 65% btw 41 and 42 |
| MBA2454144.1 | Chloroflexia bacterium | 100 Tat/SPI | 57% btw 39 and 40 |
| MBA2758935.1 | Chloroflexia bacterium | No Predicted Signal | No Predicted Signal |
| WP_052553269.1 | Enhygromyxa salina | 90.46% Sec/SPII, 09.51% Sec/SPI | 87% btw 22 and 23 |
| MBC8211257.1 | Gammaproteobacteria bacterium | 99.89% Sec/SPI | 82% btw 25 and 26 |
| MBL4851085.1 | Gammaproteobacteria bacterium | 98.16% Sec/SPI | 96% btw 32 and 33 |
| MBR9909232.1 | Gammaproteobacteria bacterium | 99.91% Sec/SPI | 98% btw 21 and 22 |

| | | | |
|---|---|---|---|
| MBU2115646.1 | Gammaproteobacteria bacterium | 99.30% Sec/SPI | 97% btw 30 and 31 |
| MCB1723870.1 | Gammaproteobacteria bacterium | 99.90% Sec/SPI | 98% btw 24 and 25 |
| MCB1800213.1 | Gammaproteobacteria bacterium | 99.91% Sec/SPI | 96% btw 25 and 26 |
| NCC26771.1 | Gammaproteobacteria bacterium | 99.89% Sec/SPI | 95% btw 25 and 26 |
| WP_157898942.1 | Luteitalea pratensis | 99.92% Sec/SPI | 98% btw 24 and 25 |
| WP_236985789.1 | Marinagarivorans sp. GE09 | 99.71% Sec/SPI | 94% btw 27 and 28 |
| WP_137437665.1 | Marinobacter sp. PJ-38 | 99.79% Sec/SPI | 97% btw 30 and 31 |
| WP_188862589.1 | Marinobacterium nitratireducens | 99.88% Sec/SPI | 56% btw 24 and 25 |
| WP_139559104.1 | Methylotetracoccus oryzae | 99.80% Sec/SPI | 97% btw 30 and 31 |
| WP_231757960.1 | Microbulbifer elongatus | 82.92% Sec/SPI, 16.95% Sec/SPII | 80% btw 25 and 26 |
| WP_183457526.1 | Microbulbifer rhizosphaerae | 70.57% Sec/SPII, 59.34% Sec/SPI | 67% btw 22 and 23 |
| WP_108732878.1 | Microbulbifer sp. A4B17 | 99.83% Sec/SPI | 94% btw 28 and 29 |
| WP_156035230.1 | Microbulbifer sp. HZ11 | 76.50% Sec/SPI, 23.37% Sec/SPII | 74% btw 25 and 26 |
| WP_152453784.1 | Microbulbifer sp. THAF38 | 99.89% Sec/SPI | 97% btw 28 and 29 |

| | | | |
|---|---|---|---|
| WP_040629207.1 | Microbulbifer variabilis | 99.88% Sec/SPI | 97% btw 28 and 29 |
| TPV96993.1 | Myxococcales bacterium FL481 | 99.89% Sec/SPI | 97% btw 25 and 26 |
| MBE2251792.1 | Myxococcus sp. | 99.30% Sec/SPI | 91% btw 12 and 13 |
| MAE34203.1 | Oceanospirillaceae bacterium | 99.87% Sec/SPI | 96% btw 28 and 29 |
| WP_076514445.1 | Oleibacter marinus | 99.89% Sec/SPI | 97% btw 28 and 29 |
| TNC83410.1 | Oleibacter sp. | 99.86% Sec/SPI | 97% btw 28 and 29 |
| KZZ11471.1 | Oleibacter sp. HI0075 | 99.89% Sec/SPI | 97% btw 28 and 29 |
| MBM3996742.1 | Planctomycetes bacterium | No Predicted Signal | No Predicted Signal |
| WP_171626078.1 | Pseudoalteromonas caenipelagi | 99.88% Sec/SPI | 97% btw 23 and 24 |
| WP_063362279.1 | Pseudoalteromonas luteoviolacea | 98.36% Sec/SPI | 93% btw 38 and 39 |
| KZN74784.1 | Pseudoalteromonas luteoviolacea H33-S | 99.89% Sec/SPI | 98% btw 21 and 22 |
| WP_192540197.1 | Pseudoalteromonas prydzensis | 75.48% Sec/SPI, 23.34% Sec/SPII | 70% btw 36 and 37 |
| WP_141240123.1 | Pseudoalteromonas sp. HM-SA03 | No Predicted Signal | No Predicted Signal |
| WP_166165103.1 | Pseudomaricurvus alcaniphilus | 99.91% Sec/SPI | 98% btw 21 and 22 |

| | | | |
|---|---|---|---|
| MCG8311860.1 | Pseudomonadales bacterium | 99.68% Sec/SPI | 96% btw 34 and 35 |
| WP_128198684.1 | Rubrivivax albus | 58.71% Sec/SPII, 40.54% Sec/SPI | 53% btw 25 and 26 |
| WP_230564822.1 | Saccharothrix Luteola | No Predicted Signal | No Predicted Signal |
| NUS64381.1 | Saccharothrix sp. | No Predicted Signal | No Predicted Signal |
| MCH1931814.1 | Shewanella shenzhenensis | 96.91% Sec/SPI | 93% btw 31 and 32 |
| WP_237443824.1 | Sinobacterium norvegicum | 99.88% Sec/SPI | 93% btw 26 and 27 |
| TVR74215.1 | Sphaerobacteraceae bacterium | 99.53% Tat/SPI | 83% btw 45 and 46 |
| WP_132875434.1 | Tamaricihabitans halophyticus | 96.01% Sec/SPI | 87% btw 29 and 30 |
| WP_093031253.1 | Thiocapsa roseopersicina | 99.90% Sec/SPI | 97% btw 25 and 26 |
| WP_200249935.1 | Thiococcus pfennigii | No Predicted Signal | No Predicted Signal |
| WP_207188716.1 | Thiocystis minor | No Predicted Signal | No Predicted Signal |
| MCF7989346.1 | Thiohalocapsa sp. | 99.91% Sec/SPI | 97% btw 25 and 26 |
| WP_017029669.1 | Vibrio breoganii 1C10 | 99.89% Sec/SPI | 97% btw 25 and 26 |
| OED83209.1 | Vibrio breoganii ZF-55 | 99.89% Sec/SPI | 97% btw 25 and 26 |

| | | | |
|---|---|---|---|
| WP_155759790.1 | Vibrio natriegens CCUG16374 | 99.89% Sec/SPI | 97% btw 25 and 26 |
| WP_118119916.1 | Vibrio sp. dhg | 99.89% Sec/SPI | 97% btw 25 and 26 |
| WP_014234363.1 | Vibrio sp. EJY3 | 99.88% Sec/SPI | 97% btw 25 and 26 |
| WP_237314952.1 | Vibrio sp. J1-1 | 98.59% Sec/SPI | 94% btw 25 and 26 |
| WP_248006878.1 | Vibrio amylolyticus | 99.87% Sec/SPI | 96% btw 25 and 26 |
| MBR9787475.1 | Vibrionaceae bacterium | 99.70% Sec/SPI | 96% btw 25 and 26 |
| MBR9875570.1 | Vibrionaceae bacterium | 98.59% Sec/SPI | 94% btw 25 and 26 |

**Table S7 – Representative VHPO accession numbers from across kingdoms**

| VHPO | Accession Number | Source Organism | General Classification |
|---|---|---|---|
| NapH1 | ABS50486.1 | Streptomyces sp. CNQ525 | Actinobacteria |
| NapH4 | ABS50492.1 | Streptomyces sp. CNQ525 | Actinobacteria |
| Mcl24 | AGH68909.1 | Streptomyces sp. CNH189 | Actinobacteria |
| Mcl40 | AGH68925.1 | Streptomyces sp. CNH189 | Actinobacteria |
| MarH1 | WP_047018063 | Streptomyces sp. CNQ-509 | Actinobacteria |
| ZgVIPO1 | CAZ96246.1 | Zobellia galactanivorans | Flavobacteria |
| ZgVIPO2 | CAZ96406.1 | Zobellia galactanivorans | Flavobacteria |
| | WP_011620573.1 | Synechococcus sp. CC9311 | Cyanobacteria |
| | WP_047157019.1 | Trichodesmium erythraeum | Cyanobacteria |
| | WP_012165216.1 | Acaryochloris marina | Cyanobacteria |
| | P81701.1 | Ascophyllum nodosum | Brown algae |
| | AJ491786.1 | Laminaria digitata | Brown algae |
| | AAC35279 | Fucus distichus | Brown algae |
| | PXF43007.1 | Gracilariopsis chorda | Red algae |
| | 1QHB_A | Corallina officinalis | Red algae |
| | XP_005714237.1 | Chondrus crispus | Red algae |
| | CAA59686 | Curvularia inaequalis | Fungi |
| | CAA72622 | Alternaria didymospora | Fungi |
| | XP_003708974.1 | Pyricularia oryzae | Fungi |
| | WP_156035230.1 | Microbulbifer sp. Hz11 | Gammaproteobacteria |
| | WP_155759790.1 | Vibrio natriegens CCUG 16374 | Gammaproteobacteria |
| | WP_076514445.1 | Oleibacter marinus | Gammaproteobacteria |
| | WP_093031253.1 | Thiocapsa roseopersicina | Gammaproteobacteria |
| | WP_171626078.1 | Pseudoalteromonas caenipelagi | Gammaproteobacteria |
| | WP_052553269.1 | Enhygromyxa salina | Deltaproteobacteria |
| | WP_157898942.1 | Luteitalea pratensis | Acidobacteria |
| | WP_128198684.1 | Rubrivivax albus | Betaproteobacteria |

| | | | |
|---|---|---|---|
| | WP_132875434.1 | Tamaricihabitans halophyticus | Actinobacteria |
| | WP_230564822.1 | Saccharothrix Luteola NEAU S10 | Actinobacteria |
| | MYD09780.1 | Chloroflexi bacterium | Chloroflexota |

**Table S8 – Halogenases used to query bacterial genomes for halogenation potential**

| Halogenase family | Protein Signifier | UniProt Accession |
|---|---|---|
| flavin-dependent halogenase | ThaL | A1E280 |
| non-heme iron/α-ketoglutarate dependent halogenase | BesD | G8XHD5 |
| SAM-dependent adenosyl-chloride synthase | SalL | A4X3Q0 |
| heme chloroperoxidase | CPO | P04963 |
| Vanadium-dependent haloperoxidase | HZ11-VHPO | *see table S1* |

**Table S9 – Halogenases in Oleibacter marinus, Vibrio natriegens, and Microbulbifer rhizosphaerae**

| Oleibacter marinus | JGI Gene ID | % identity to query |
|---|---|---|
| flavin-dependent halogenase | N/A | N/A |
| non-heme iron/α-ketoglutarate dependent halogenase | N/A | N/A |
| SAM-dependent adenosyl-chloride synthase | N/A | N/A |
| heme chloroperoxidase | N/A | N/A |
| Vanadium-dependent haloperoxidase | 2684411802 | 46 |

| Vibrio natriegens | JGI Gene ID | % identity to query |
|---|---|---|
| flavin-dependent halogenase | N/A | N/A |
| non-heme iron/α-ketoglutarate dependent halogenase | N/A | N/A |
| SAM-dependent adenosyl-chloride synthase | N/A | N/A |
| heme chloroperoxidase | N/A | N/A |
| Vanadium-dependent haloperoxidase | 2813788644 | 41 |

| Microbulbifer rhizosphaerae | JGI Gene ID | % identity to query |
|---|---|---|
| flavin-dependent halogenase | 2824283566 | 35 |
| flavin-dependent halogenase | 2824283088 | 33 |
| flavin-dependent halogenase | 2824283085 | 38 |
| flavin-dependent halogenase | 2824283090 | 34 |
| flavin-dependent halogenase | 2824282115 | 30 |
| flavin-dependent halogenase | 2824283797 | 32 |
| flavin-dependent halogenase | 2824283089 | 24 |
| flavin-dependent halogenase | 2824283589 | 37 |
| non-heme iron/α-ketoglutarate dependent halogenase | N/A | N/A |
| SAM-dependent adenosyl-chloride synthase | N/A | N/A |
| heme chloroperoxidase | N/A | N/A |
| Vanadium-dependent haloperoxidase | 2824281323 | 49 |
